# CCC-GPU: A graphics processing unit (GPU)-accelerated nonlinear correlation coefficient for large-scale transcriptomic analyses

**DOI:** 10.1101/2025.06.03.657735

**Authors:** Haoyu Zhang, Kevin Fotso, Marc Subirana-Granés, Milton Pividori

## Abstract

**Motivation:** Identifying meaningful patterns in complex biological data necessitates correlation coefficients capable of capturing diverse relationship types beyond simple linearity. Furthermore, efficient computational tools are crucial for handling the ever-increasing scale of biological datasets.

**Results:** We introduce CCC-GPU, a high-performance, GPU-accelerated implementation of the Clustermatch Correlation Coefficient (CCC). CCC-GPU computes correlation coefficients for mixed data types, effectively detects nonlinear relationships, and offers significant speed improvements over its predecessor.

**Availability and Implementation:** CCC-GPU is openly available on GitHub (https://github.com/pivlab/ccc-gpu) and distributed under the BSD-2-Clause Plus Patent License.

## Introduction

Correlation coefficients are fundamental tools for uncovering meaningful patterns within data. While traditional measures like Pearson and Spearman are adept at capturing linear and monotonic relationships, newer methodologies, such as the Clustermatch Correlation Coefficient (CCC) [1] and the Maximal Information Coefficient (MIC) [2,3] have emerged to detect a broader spectrum of associations.

The CCC is a clustering-based statistic designed to identify both linear and nonlinear patterns. Previously, we demonstrated CCC’s utility in analyzing gene expression data from GTEx, showcasing its robustness to outliers and its ability to detect both strong linear relationships and biologically significant nonlinear patterns often missed by conventional coefficients [1]. Unlike MIC, CCC offers the distinct advantage of accommodating both numerical and categorical data types and offers speeds up to a two orders of magnitude faster. However, despite leveraging CPU multi-threading for acceleration, the original CCC implementation can still be computationally expensive for large datasets. For instance, in the original study [1], we used only the top 5,000 most variable genes in a single tissue (whole blood) to ease computation.

Here, we introduce a new implementation, CCC-GPU, that harnesses the power of NVIDIA CUDA for GPU programming. We have computed CCC-GPU values for all ~50,000 genes in GTEx v8 [4] across all 54 tissues in a fraction of the time that would have been needed using the original CCC implementation. This advancement achieves a substantial speedup over the original CPU-based implementation, making comprehensive correlation analysis of large biological datasets more practical and efficient.

## Methods

The original CCC was developed purely in Python. While several NVIDIA CUDA-based GPU acceleration Python libraries now exist, achieving maximum flexibility and leveraging CUDA’s complete feature set requires native C++ code [5]. Rather than fully re-implementing CCC in another programming language, we strategically integrated the original Python codebase with high-performance C++ components. This hybrid approach preserves Python interfaces and maintains the original implementation’s comprehensive feature set, ensuring robust testing and evaluation.

Profiling of the original CCC revealed that Adjusted Rand Index (ARI) computation [6] was the primary performance bottleneck. We addressed this by implementing the coefficient computation module in CUDA C++, enabling distribution of computationally intensive ARI calculations across GPU parallel processing cores. We then used pybind11 [7] to seamlessly integrate the CUDA C++ backend with the existing Python framework. This ensures full compatibility while allowing users to interact with CCC-GPU through the same Python interfaces as the original CCC.

## Results

We conducted a comprehensive comparative evaluation of Pearson, Spearman, and CCC-GPU using simulated data and all genes and tissues in the GTEx v8 dataset. The evaluation was performed on an AMD Ryzen Threadripper 7960X CPU with an NVIDIA RTX 4090 GPU, representing a typical high-performance workstation configuration.

We compared the computational complexity of CCC-GPU (using the GPU) and the original CCC (using 24 CPU cores) when analyzing each tissue in the GTEx dataset. For whole blood (56,200 genes, 755 samples), CCC-GPU demonstrated a 37x speedup. On average, the original CCC required approximately 6 hours to process one GTEx tissue. This acceleration enabled the complete analysis of all 54 GTEx tissues in just 8 hours on a single machine, compared to ~312 hours (~13 days) that would be needed with the original implementation. Consistent performance gains were observed across all tissue types, confirming CCC-GPU’s broad applicability.

To further understand CCC-GPU’s scalability with increasing data size, we performed benchmarks using synthesized input, comparing its speedup against the original CPU implementation. The results, illustrated in Figure 1a, demonstrate that CCC-GPU maintains a stable speedup trend as input size increases, confirming its robust scaling capabilities when handling large datasets. The different curves in Figure 1a simulate CCC-GPU’s prospective speedup over CPU hardware with varying numbers of cores. We also compared the runtime of the original CCC, CCC-GPU, Pearson, and Spearman methods, as shown in Figure 1b. While Pearson and Spearman are inherently faster due to their reliance on simple statistics, CCC-GPU’s runtime is remarkably close to these methods, showcasing its efficiency while maintaining its advanced capabilities for capturing complex relationships.

**Figure 1.**
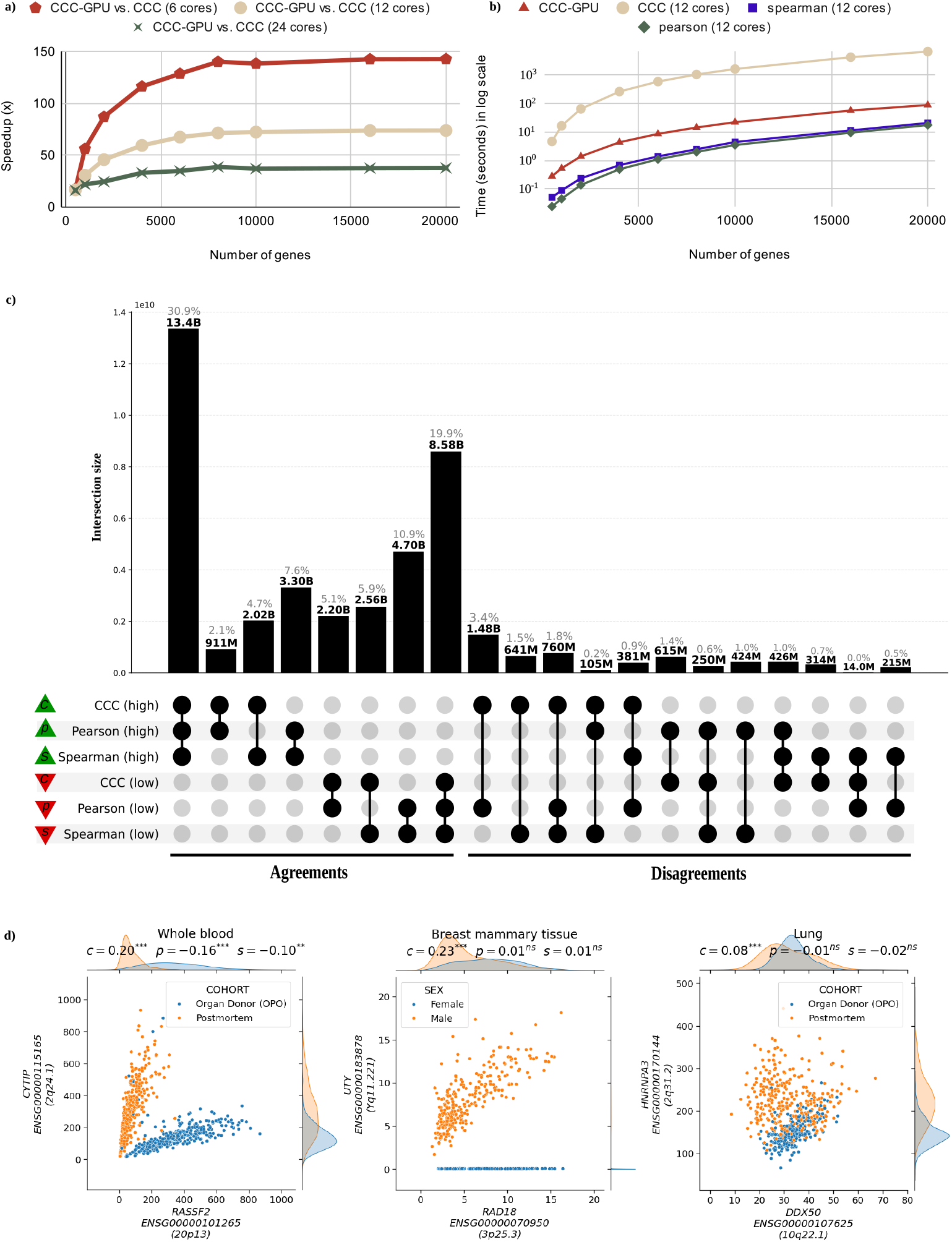
a) CCC-GPU scalability analysis showing speedup relative to CPU-based CCC across varying gene counts (1,000 samples fixed) and CPU core configurations. See Supplementary Table S1 for detailed metrics. b) Runtime comparison of correlation methods (Pearson, Spearman, original CCC, CCC-GPU) demonstrating CCC-GPU’s competitive performance despite its complexity to capture nonlinear patterns. Analysis used 12 CPU cores with varying gene counts (1,000 samples fixed). See Supplementary Table S2. c) UpSet plot showing correlation agreement (left) and disagreement (right) patterns of all gene pairs in all 54 GTEx v8 tissues among methods. Green triangles indicate high correlations (top 30% gene pairs per tissue), and red triangles show low correlations (bottom 30%). We also created UpSet plots using permutation-based thresholds (see Supplementary Information and Supplementary Figure S1). UpSet plots for individual tissues are provided in Supplementary Figures S2 to S55 in panels c and d. d) Expression levels of selected gene pairs with associations across GTEx tissues: *RASSF2*-*CYTIP*, in whole blood, shows two clear and strong linear patterns across different sample subsets (postmortem versus organ donor). Although all coefficients are statistically significant, Pearson and Spearman detect the wrong pattern (i.e., their slopes are negative); *UTY*-*RAD18* shows a linear relationship only in male samples, whereas female samples exhibit a constant value of zero (since *UTY* is located in chromosome Y). *HNRNPA3*-*DDX50* demonstrates a case where one sample subset (organ donor) shows a linear relationship that is masked by another subset (postmortem) with no or weak relationship.

The significant performance enhancement provided by CCC-GPU expands the scope of transcriptomic analyses and allowed us to uncover more nonlinear patterns. In the UpSet analysis [8] shown in Figure 1c, we compared how Pearson, Spearman, and CCC-GPU agreed or disagreed in prioritizing gene pairs across all genes in the 54 GTEx tissues. We found that ~2.9 billion gene pairs found by CCC-GPU only (Disagreements in Figure 1c, where CCC-GPU is “high” and any of the others “low”) likely have biologically meaningful nonlinear patterns, as it was found in the original CCC study [1]. Likewise, gene pairs where Pearson is “high” and the rest are “low” are likely driven by outliers [1].

Leveraging CCC-GPU’s mixed data type capability, we correlated gene expression with GTEx metadata to interpret nonlinear patterns. For this, we included categorical variables such as sex, mortality status and numerical variables such as BMI, age, among others. From the five intersection groups where CCC values were high but Pearson or Spearman remained low, we selected the top 100 gene pairs per tissue with the largest CCC value. Figure 1d highlights gene pairs with biologically interpretable nonlinear patterns explained by a metadata variable, such as *UTY*-*KDM6A* (sex) and *RASSF2*-*CYTIP* (pre/post-mortem status; these two genes were previously found to be differentially expressed in these conditions [9]). We provide the CCC values computed for all gene pairs, and the top gene-metadata correlation results across all GTEx v8 tissues (see Supplementary Note 1).

## Conclusions

We present CCC-GPU, a GPU-accelerated implementation of the original Clustermatch Correlation Coefficient (CCC). Our work demonstrates that CCC-GPU delivers a remarkable acceleration over its CPU-based predecessor, enabling the rapid and efficient computation of correlation coefficients in large transcriptomic datasets. This performance leap transforms analyses that previously required weeks into tasks achievable within hours on standard research hardware.

Challenges often reside not only in detecting but also in interpreting complex, nonlinear patterns between genes. CCC-GPU’s ability to correlate different data types allows researchers to easily incorporate available metadata into their analyses. For example, a previously highlighted nonlinear relationship for the gene pair *RASSF2*-*CYTIP*, detected by CCC, was explained by GTEx metadata field “COHORT” (pre-vs. post-mortem status). These two genes were previously reported to be differentially expressed under these conditions [9]. CCC also detected a strong linear pattern between these genes regardless of the organism’s mortality status.

Our new CCC-GPU delivers a next-generation correlation coefficient at a fraction of the computational cost without sacrificing its accessibility, accuracy or reliability. This significant performance enhancement makes comprehensive correlation analysis of large genomic data practical on standard research hardware. Beyond standard correlation analyses, CCC-GPU empowers biologists to perform sophisticated tasks such as advanced feature selection prior to machine learning model training with large datasets. Moreover, by accelerating the discovery of novel nonlinear relationships in expression data, CCC-GPU helps researchers quickly identify patterns beyond conventional, linear-only relationships. This can potentially uncover new biological insights that would otherwise remain hidden and drive forward our understanding of complex biological systems.

## Acknowledgements

This work is supported by the National Human Genome Research Institute (R00 HG011898 to M.P.), and The Eunice Kennedy Shriver National Institute of Child Health and Human Development (R01 HD109765 to M.P.).

## Notes

https://github.com/pivlab/ccc-gpu

https://doi.org/10.5281/zenodo.17156519

### Competing Interest Statement

The authors have declared no competing interest.

### Summary of Updates

Figure 1 has been updated with new results that confirm previous findings. Text has been revised and improved. Links to Zenodo with datasets have been added. Supplementary material has been significantly extended.

