## Supplementary Information, Notes, Figures and Tables for "CCC-GPU: A graphics processing unit (GPU)-accelerated nonlinear correlation coefficient for large-scale transcriptomic analyses"

#### Supplementary Note 1: Datasets

In our Zenodo archive (https://doi.org/10.5281/zenodo.17156519), we provide: 1) the CCC values for all gene pairs across all 54 GTEx v8 tissues, 2) the threshold tables for top and bottom genes using both the 30% threshold and the permutation-based thresholds (see section below for an explanation of these thresholds), and 3) the top gene-metadata correlation results for all tissues.

#### Supplementary Note 2: high and low correlation thresholds

We used two approaches to define correlation values that are "high" or "low" for each coefficient per tissue. The first approach captures, for each tissue and correlation coefficient, the top 30% of gene pairs by using the 70th percentile of correlation values, and the bottom 30% of gene pairs by using the 30th percentile (Figure 1c, in the main text, for all tissues combined; and panel c for individual tissues in the supplementary figures section below). The second approach computes the null distribution of coefficient values per tissue by taking a random subset of 10,000 genes and shuffling samples. Then, we define genes with "high" correlation as those with coefficient values larger than the 95th percentile ($P < 0.05$), and "low" correlation as those with coefficient values smaller than the 80th percentile ($P >$ 0.20) (Figure S1 for all tissues, and panel d for individual tissues, in the supplementary figures section below).

#### Tables

**Table S1: Speedup using different CPU configurations (1,000 fixed samples).**

| Number of genes | CCC-GPU vs. CCC (6 cores) | CCC-GPU vs. CCC (12 cores) | CCC-GPU vs. CCC (24 cores) |
|---|---|---|---|
| 500 | 17.6x | 16.52x | 16.1x |
| 1,000 | 56.17x | 30.65x | 21.82x |
| 2,000 | 87.06x | 45.72x | 24.45x |
| 4,000 | 116.39x | 59.46x | 33.03x |
| 6,000 | 128.74x | 67.46x | 34.77x |
| 8,000 | 140.24x | 71.48x | 38.67x |
| 10,000 | 138.51x | 72.38x | 37.02x |
| 16,000 | 142.7x | 73.83x | 37.53x |
| 20,000 | 142.83x | 73.88x | 37.73x |

**Table S2: Execution time of different methods (1,000 fixed samples).**

| Number of genes | CCC-GPU | CCC (12 cores) | Spearman (12 cores) | Pearson (12 cores) |
|---|---|---|---|---|
| 500 | 0.279s | 4.603s | 0.051s | 0.024s |
| 1,000 | 0.529s | 16.198s | 0.089s | 0.045s |
| 2,000 | 1.384s | 63.263s | 0.235s | 0.138s |
| 4,000 | 4.290s | 255.071s | 0.679s | 0.489s |

| Number of genes | CCC-GPU | CCC (12 cores) | Spearman (12 cores) | Pearson (12 cores) |
|---|---|---|---|---|
| 6,000 | 8.441s | 569.409s | 1.379s | 1.093s |
| 8,000 | 14.116s | 1009.069s | 2.443s | 1.973s |
| 10,000 | 21.694s | 1570.269s | 4.379s | 3.448s |
| 16,000 | 55.469s | 4051.114s | 11.169s | 9.310s |
| 20,000 | 86.286s | 6374.781s | 20.282s | 17.238s |

### Figures

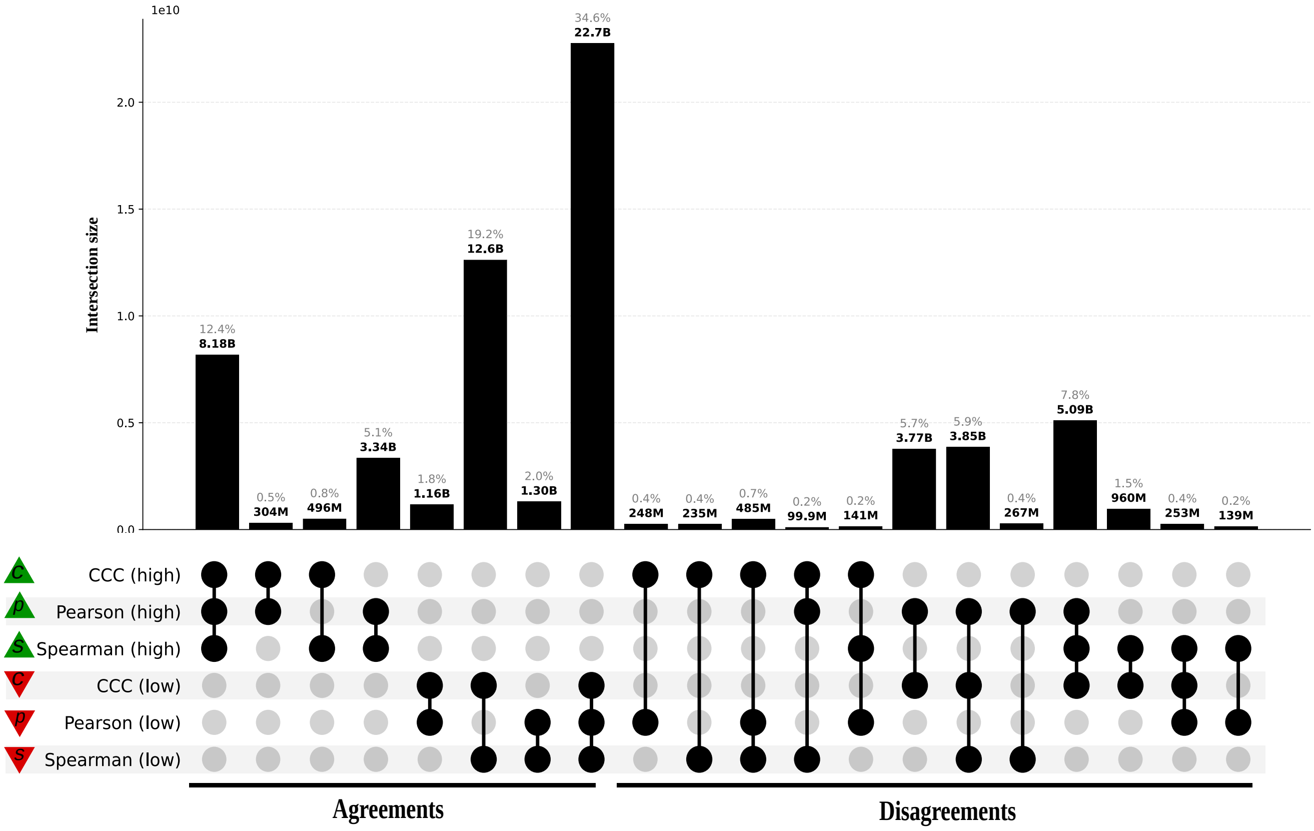


**Figure S1: UpSet plot for gene pairs in all 54 tissues in GTEx using permutation-based thresholds for each coefficient for grouping.**

#### Adipose Subcutaneous

**a)** Correlation coefficient distributions between gene pairs within GTEx v8 Adipose Subcutaneous

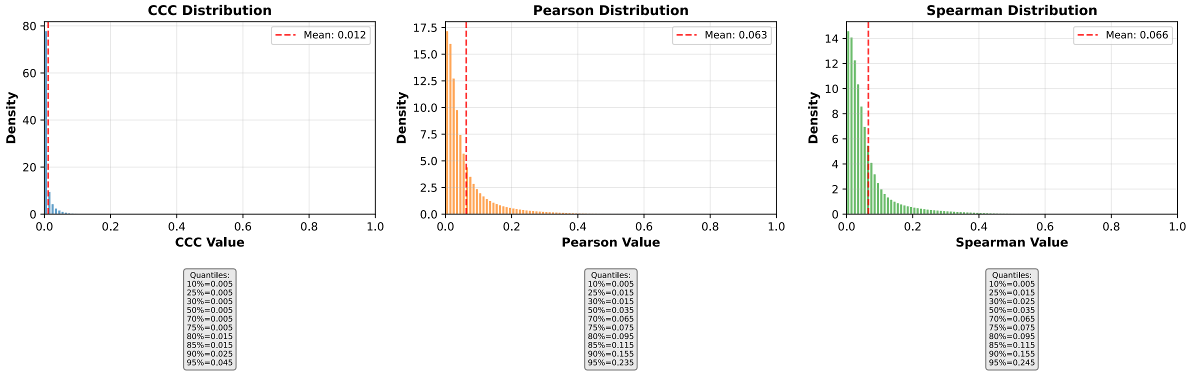


**b)** Corresponding cumulative histogram

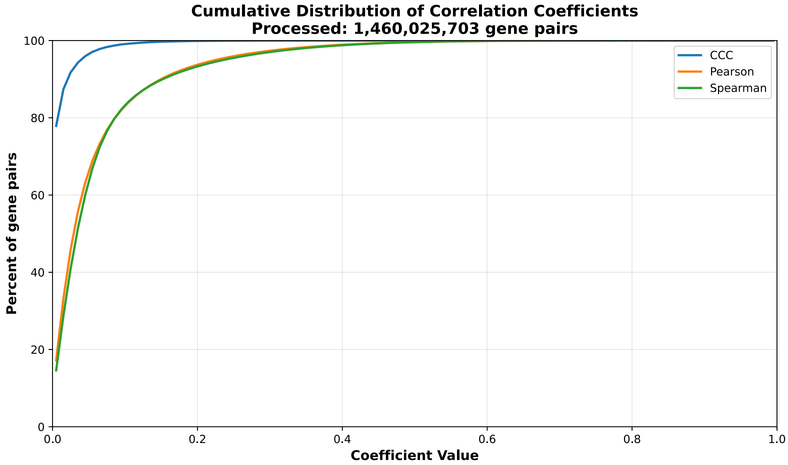


**c)** UpSet plot using top and bottom 30% correlations

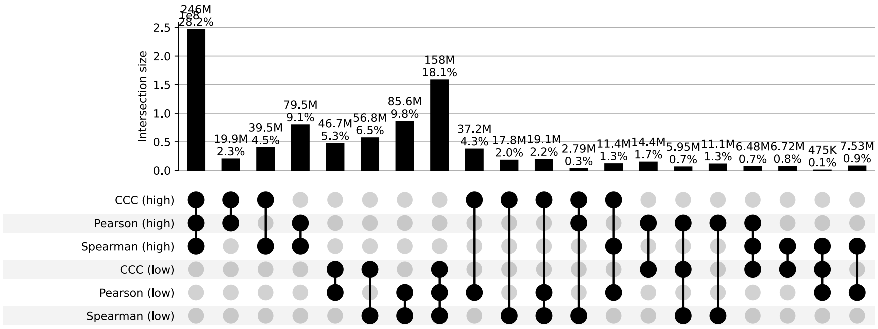


**d)** UpSet plot using permutation-based statistical thresholds

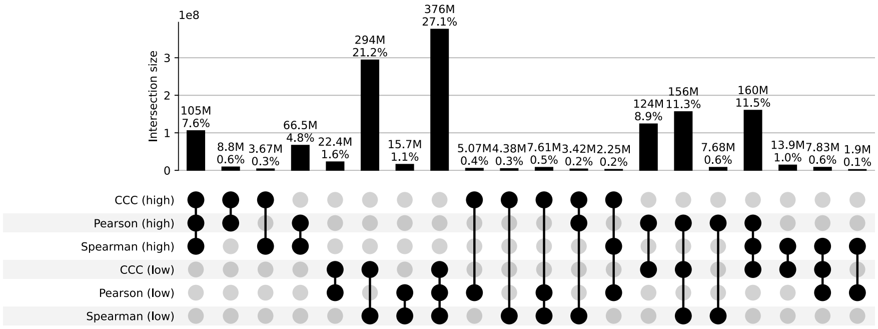


**Figure S2: Distribution and UpSet plots for GTEx v8 adipose subcutaneous.**

#### Adipose Visceral Omentum

**a)** Correlation coefficient distributions between gene pairs within GTEx v8 Adipose Visceral Omentum

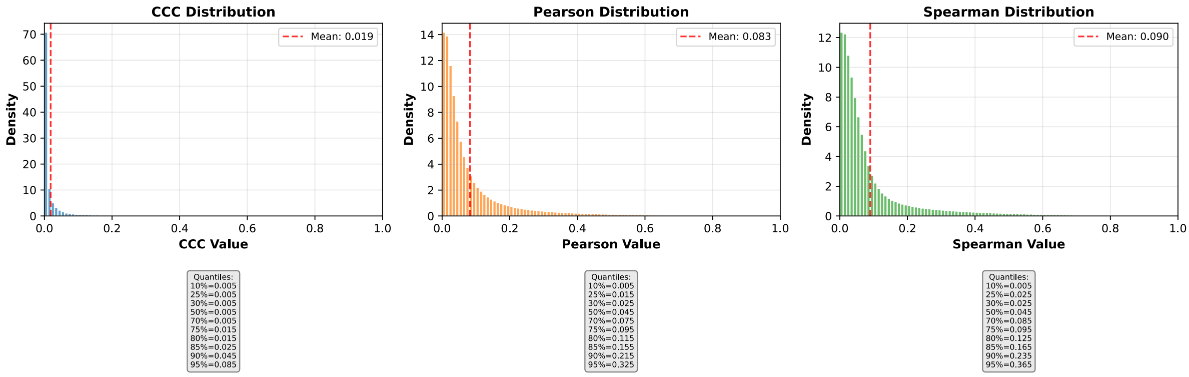


**b)** Corresponding cumulative histogram

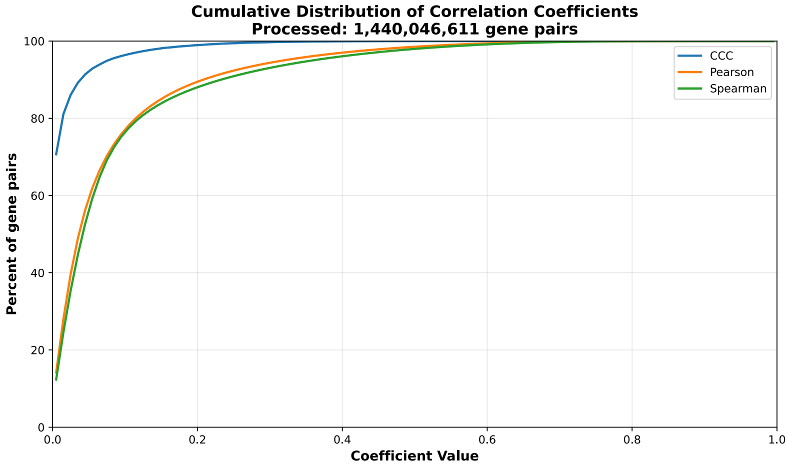


**c)** UpSet plot using top and bottom 30% correlations

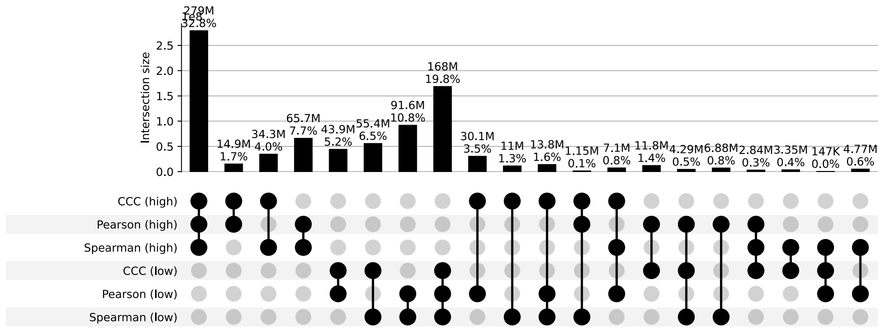


**d)** UpSet plot using permutation-based statistical thresholds

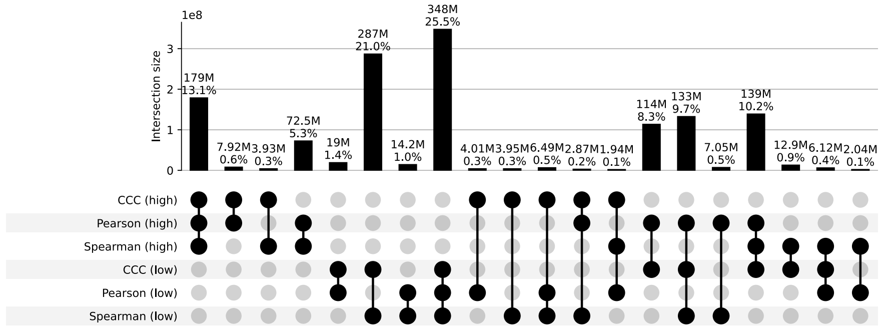


**Figure S3: Distribution and UpSet plots for GTEx v8 adipose visceral omentum.**

#### Adrenal Gland

**a)** Correlation coefficient distributions between gene pairs within GTEx v8 Adrenal Gland

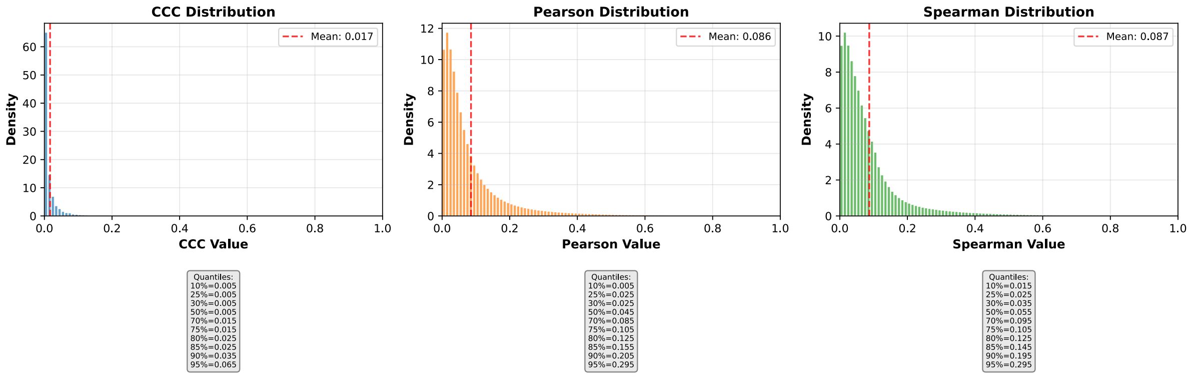


**b)** Corresponding cumulative histogram

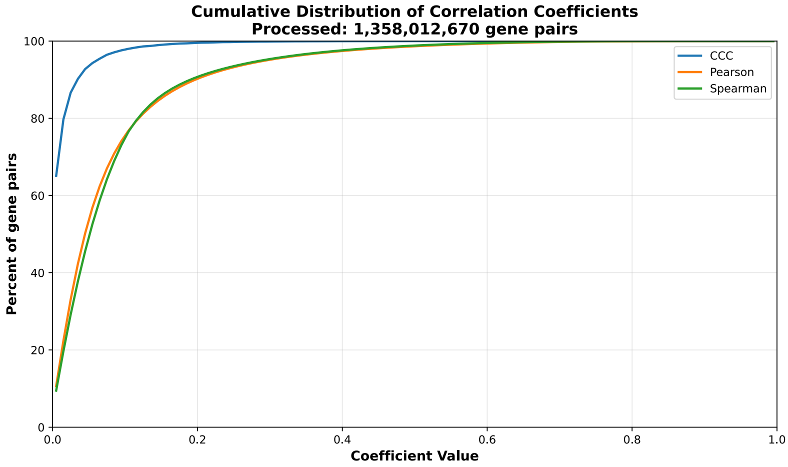


**c)** UpSet plot using top and bottom 30% correlations

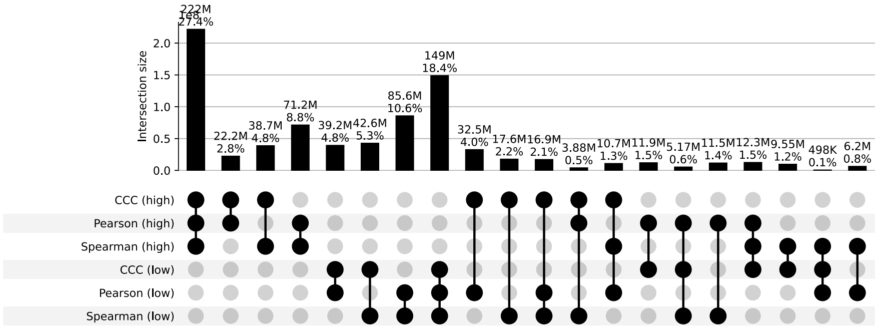


**d)** UpSet plot using permutation-based statistical thresholds

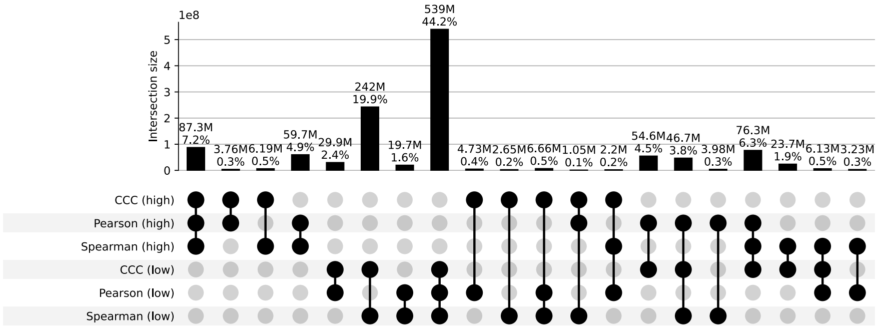


**Figure S4: Distribution and UpSet plots for GTEx v8 adrenal gland.**

#### Artery Aorta

**a)** Correlation coefficient distributions between gene pairs within GTEx v8 Artery Aorta

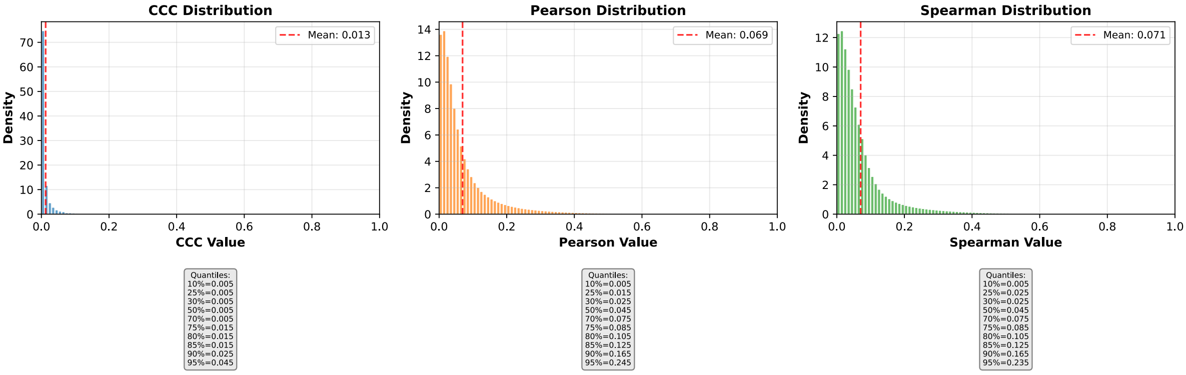


**b)** Corresponding cumulative histogram

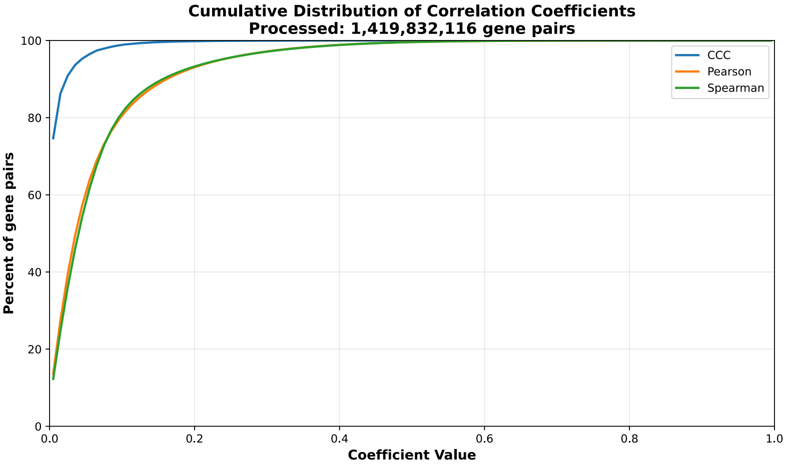


**c)** UpSet plot using top and bottom 30% correlations

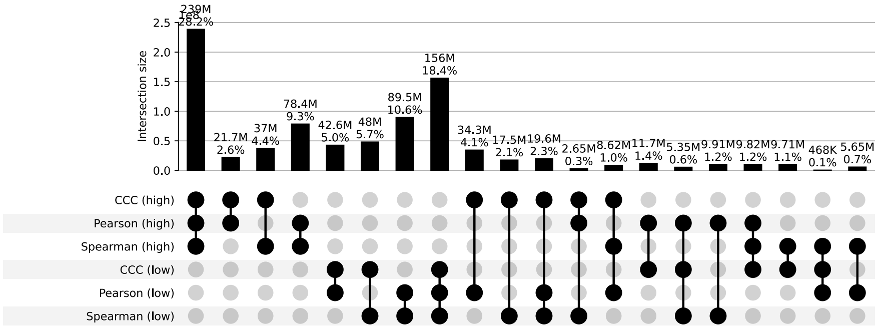


**d)** UpSet plot using permutation-based statistical thresholds

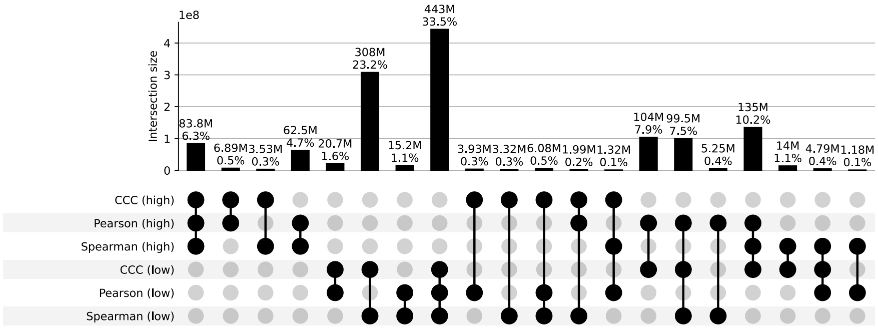


**Figure S5: Distribution and UpSet plots for GTEx v8 artery aorta.**

#### Artery Coronary

**a)** Correlation coefficient distributions between gene pairs within GTEx v8 Artery Coronary

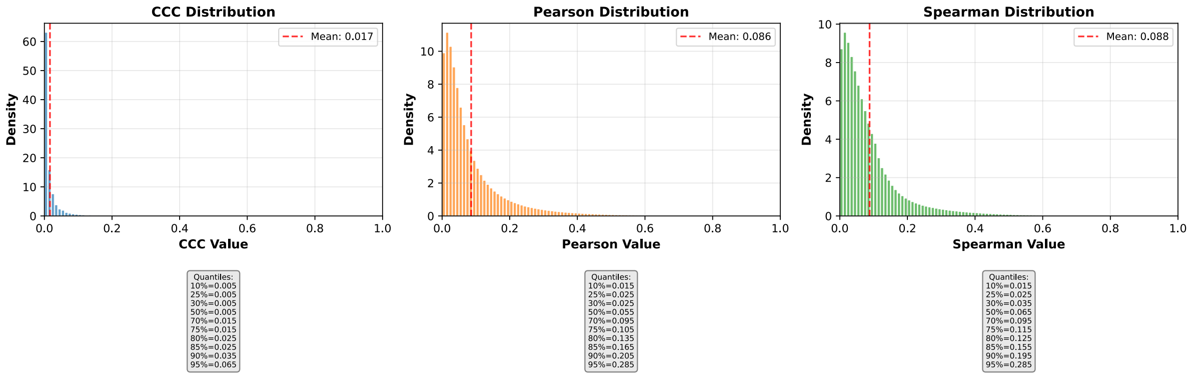


**b)** Corresponding cumulative histogram

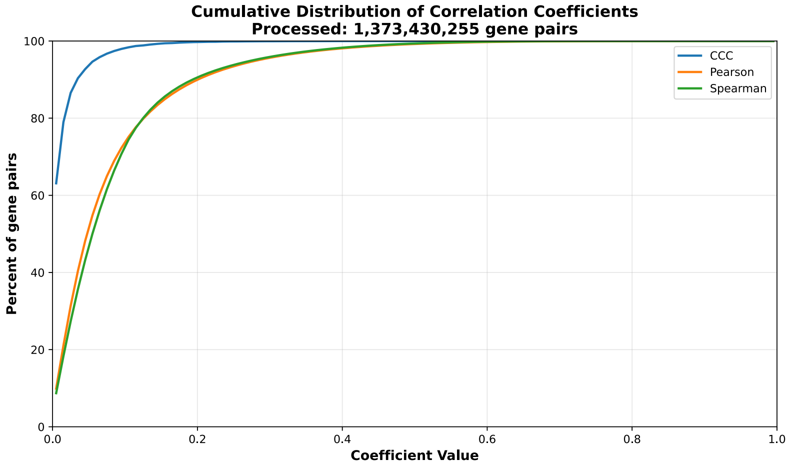


**c)** UpSet plot using top and bottom 30% correlations

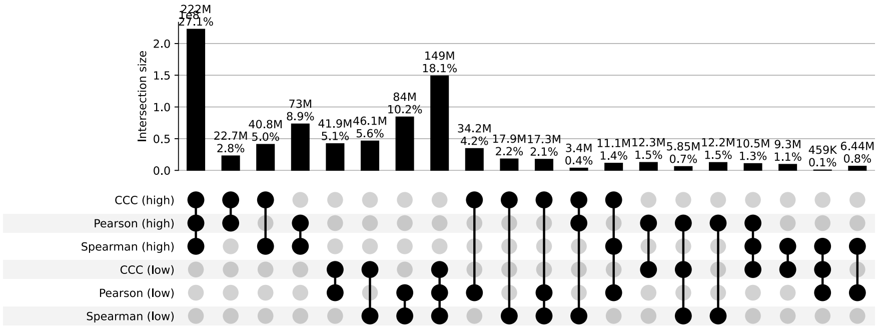


**d)** UpSet plot using permutation-based statistical thresholds

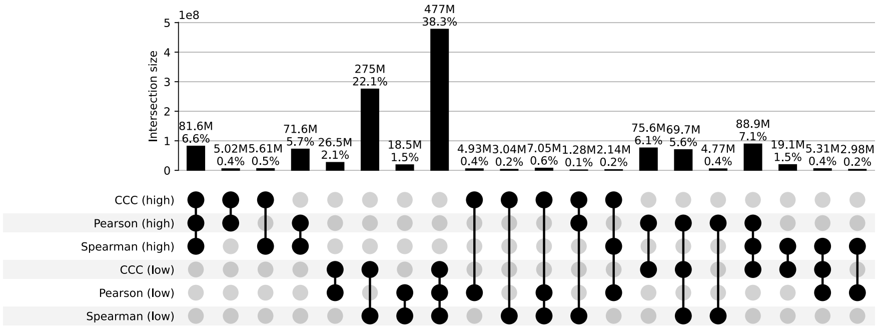


**Figure S6: Distribution and UpSet plots for GTEx v8 artery coronary.**

#### Artery Tibial

**a)** Correlation coefficient distributions between gene pairs within GTEx v8 Artery Tibial

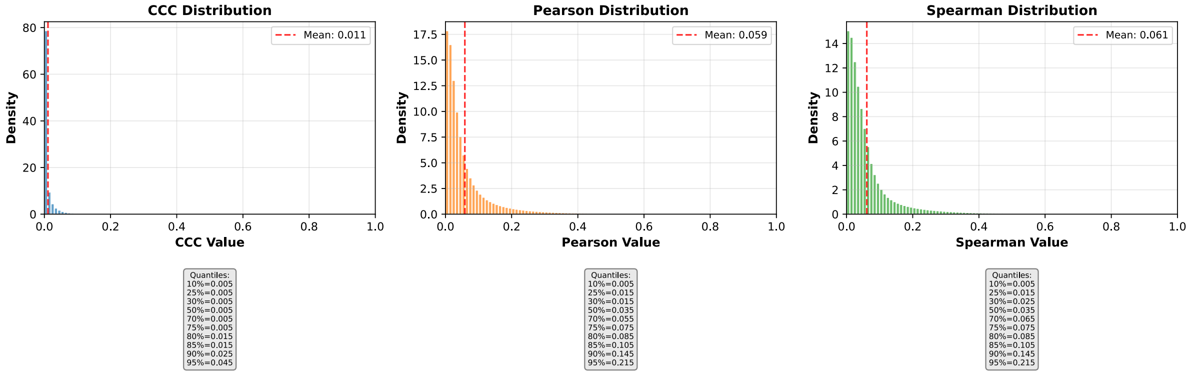


**b)** Corresponding cumulative histogram

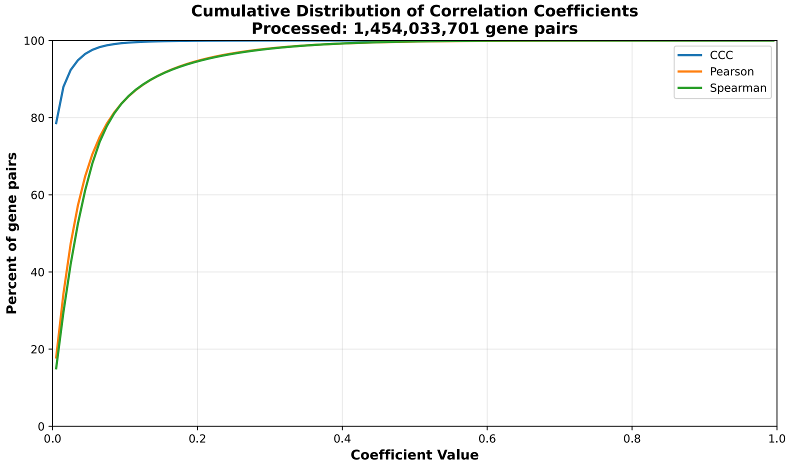


**c)** UpSet plot using top and bottom 30% correlations

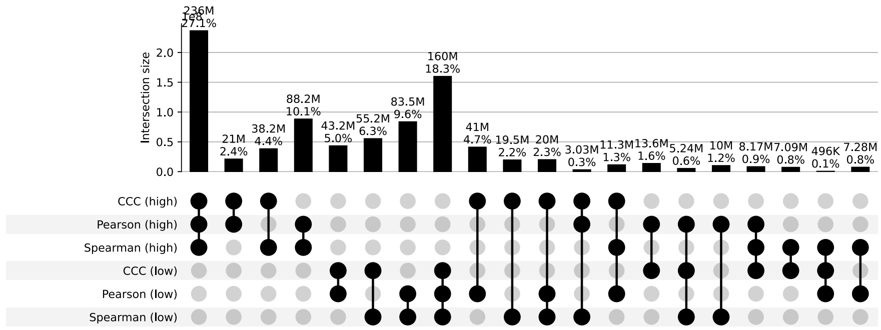


**d)** UpSet plot using permutation-based statistical thresholds

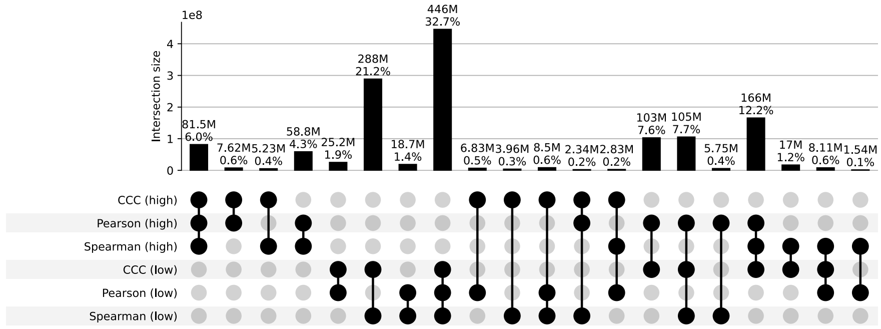


**Figure S7: Distribution and UpSet plots for GTEx v8 artery tibial.**

#### Bladder

**a)** Correlation coefficient distributions between gene pairs within GTEx v8 Bladder

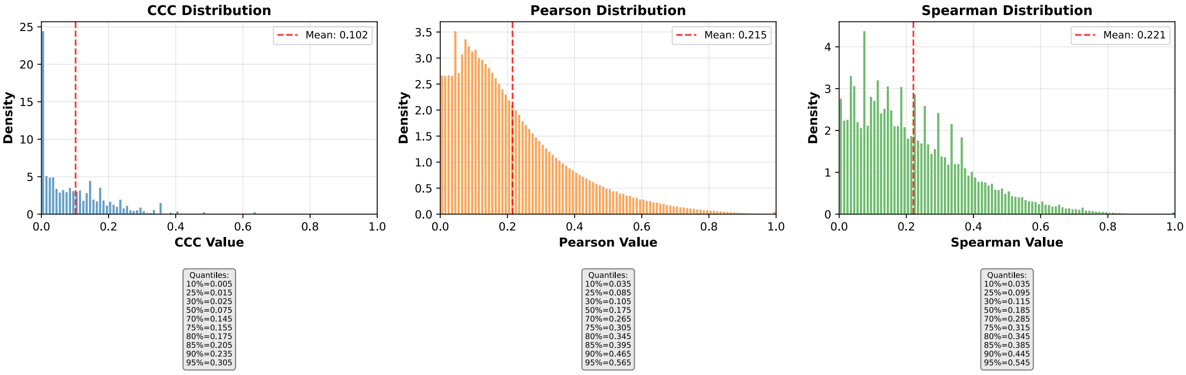


**b)** Corresponding cumulative histogram

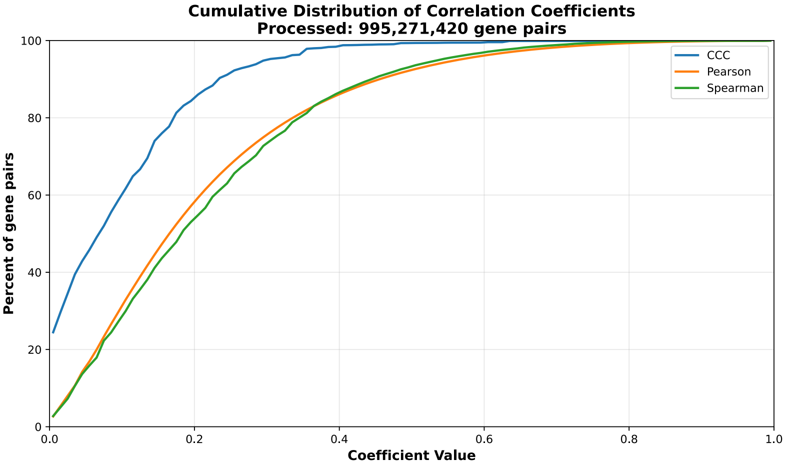


**c)** UpSet plot using top and bottom 30% correlations

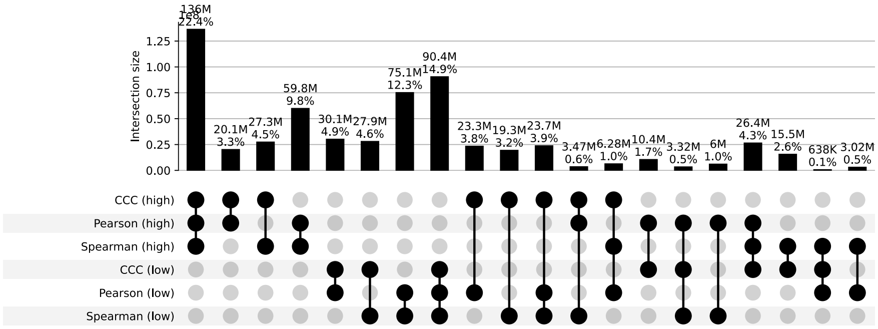


**d)** UpSet plot using permutation-based statistical thresholds

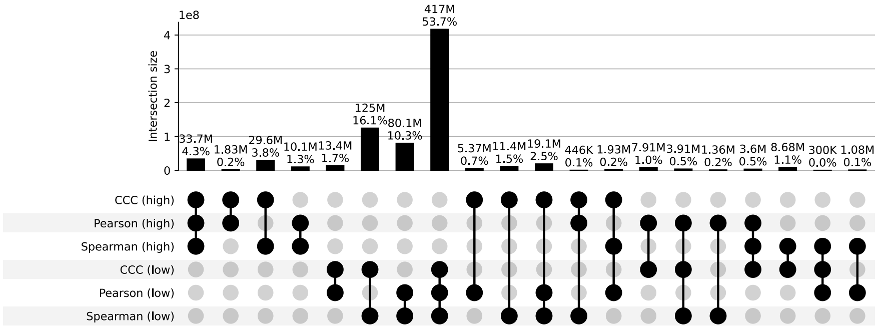


**Figure S8: Distribution and UpSet plots for GTEx v8 bladder.**

#### Brain Amygdala

**a)** Correlation coefficient distributions between gene pairs within GTEx v8 Brain Amygdala

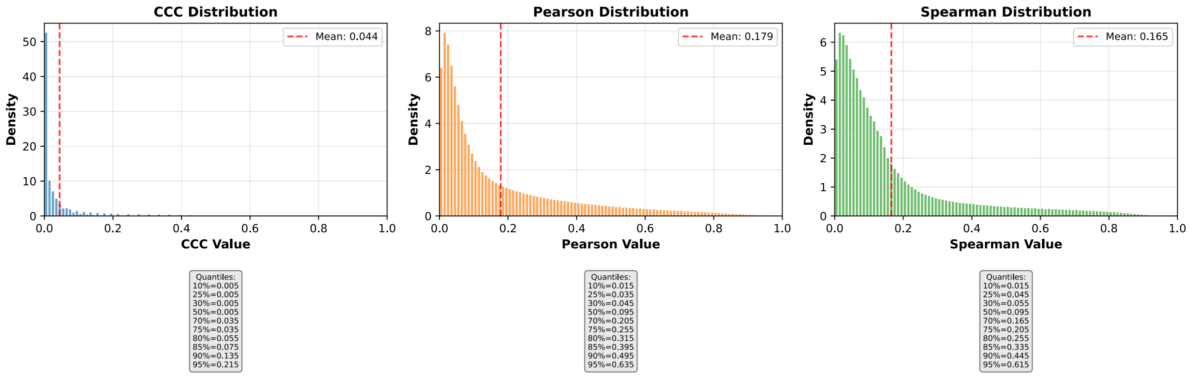


**b)** Corresponding cumulative histogram

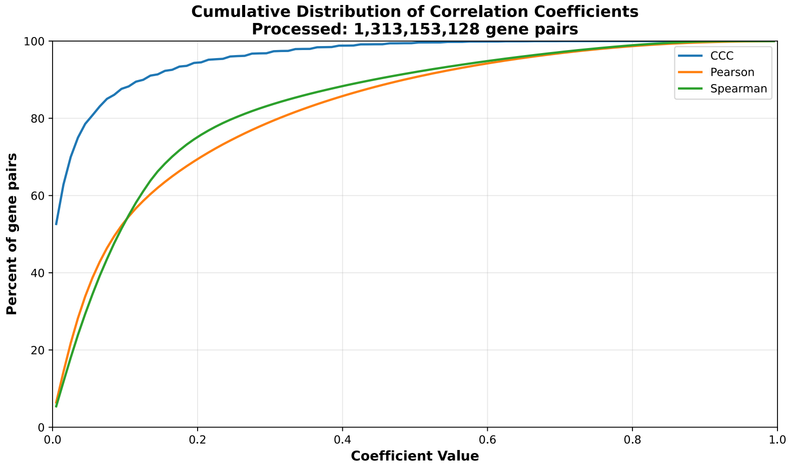


**c)** UpSet plot using top and bottom 30% correlations

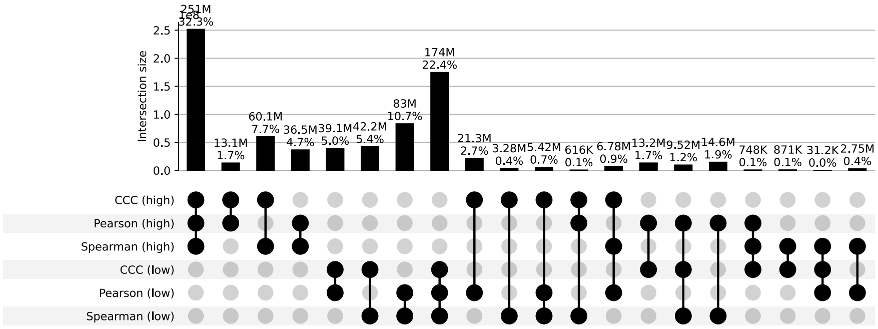


**d)** UpSet plot using permutation-based statistical thresholds

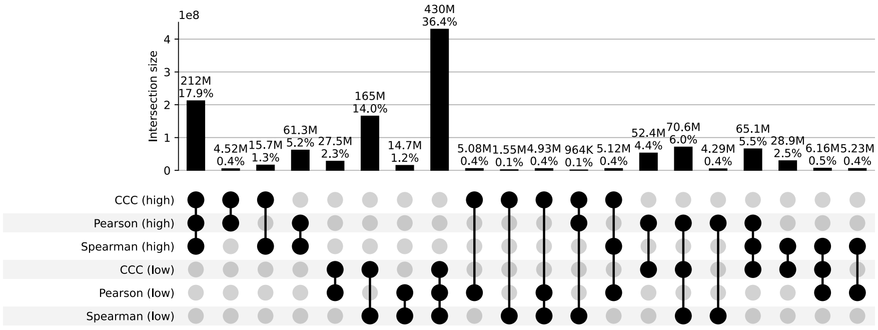


**Figure S9: Distribution and UpSet plots for GTEx v8 brain amygdala.**

#### Brain Anterior Cingulate Cortex Ba24

**a)** Correlation coefficient distributions between gene pairs within GTEx v8 Brain Anterior Cingulate Cortex Ba24

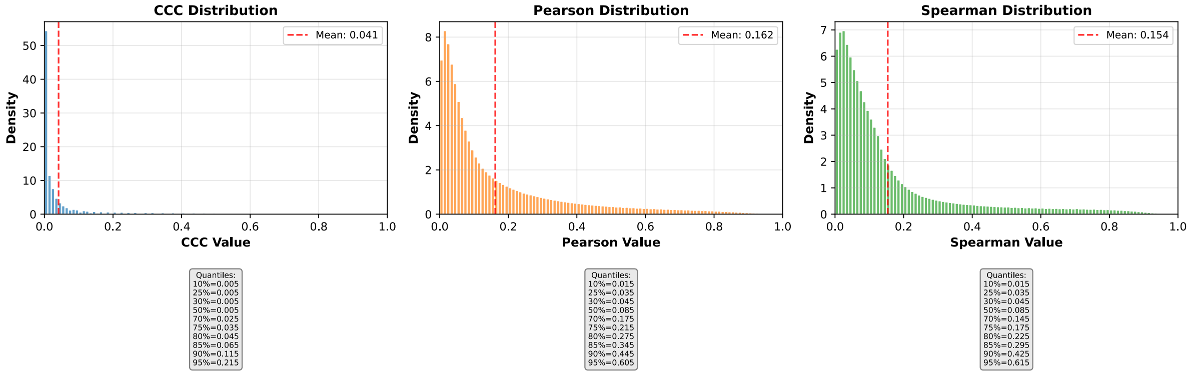


**b)** Corresponding cumulative histogram

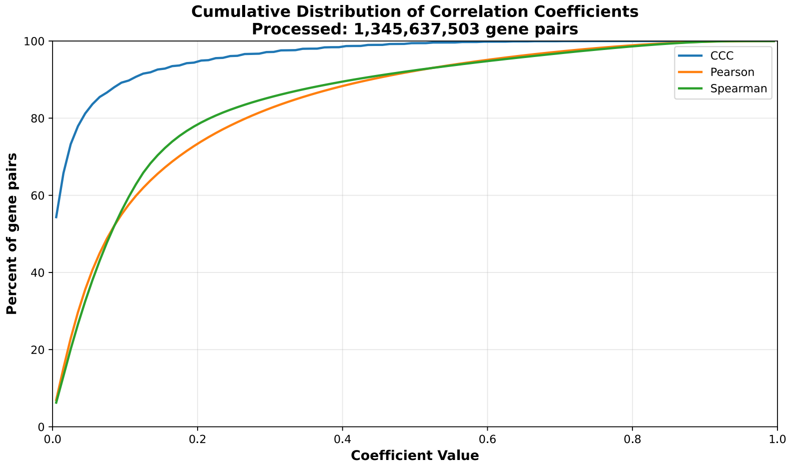


**c)** UpSet plot using top and bottom 30% correlations

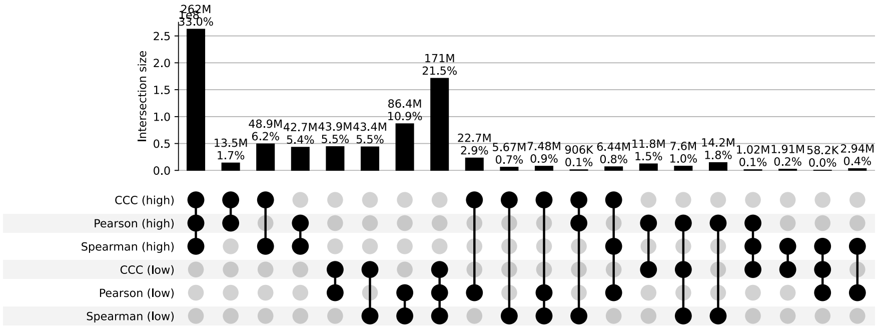


**d)** UpSet plot using permutation-based statistical thresholds

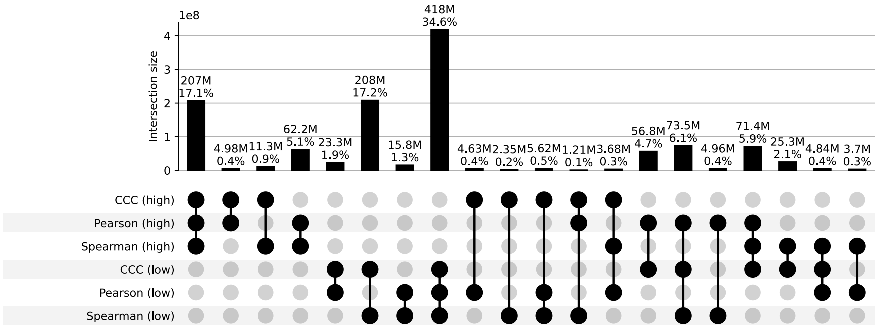


**Figure S10: Distribution and UpSet plots for GTEx v8 brain anterior cingulate cortex BA24.**

#### Brain Caudate Basal Ganglia

**a)** Correlation coefficient distributions between gene pairs within GTEx v8 Brain Caudate Basal Ganglia

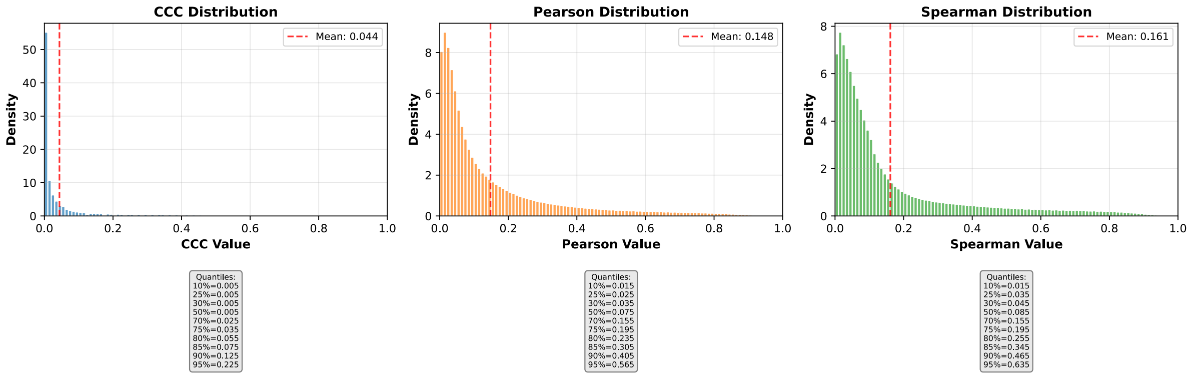


**b)** Corresponding cumulative histogram

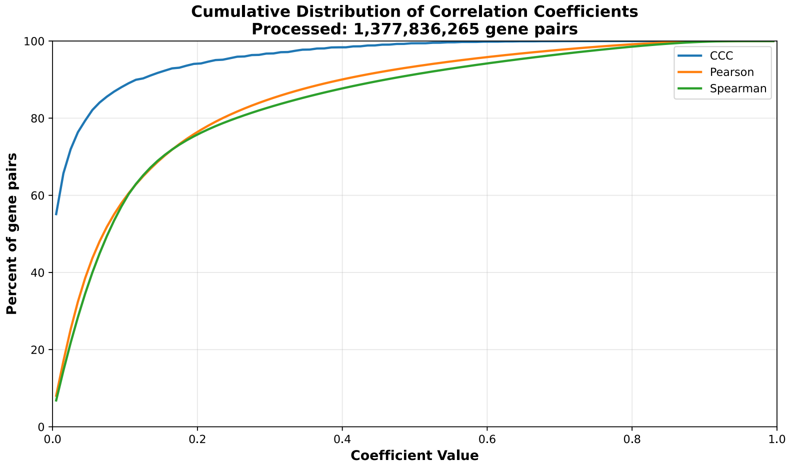


**c)** UpSet plot using top and bottom 30% correlations

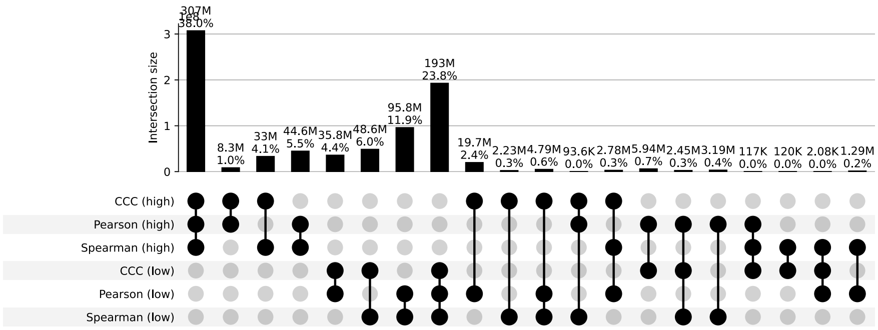


**d)** UpSet plot using permutation-based statistical thresholds

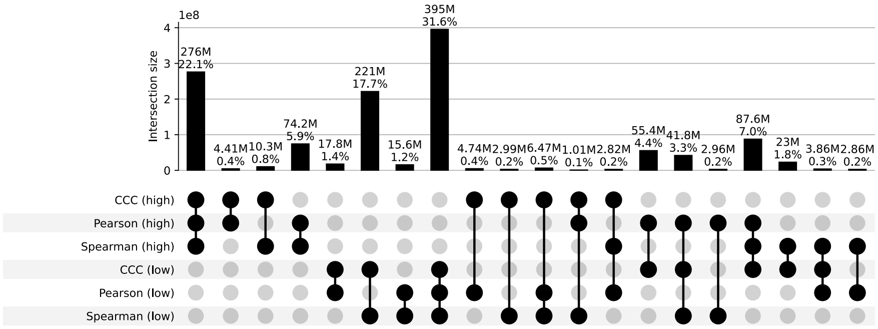


**Figure S11: Distribution and UpSet plots for GTEx v8 brain caudate basal ganglia.**

#### Brain Cerebellar Hemisphere

**a)** Correlation coefficient distributions between gene pairs within GTEx v8 Brain Cerebellar Hemisphere

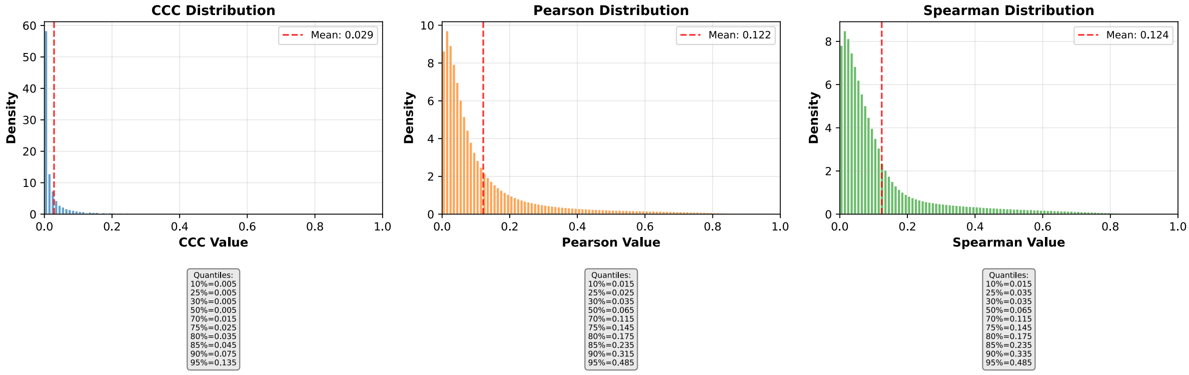


**b)** Corresponding cumulative histogram

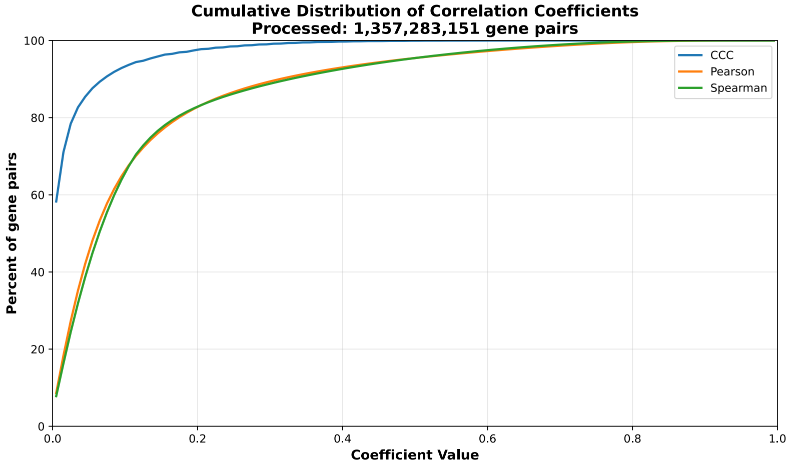


**c)** UpSet plot using top and bottom 30% correlations

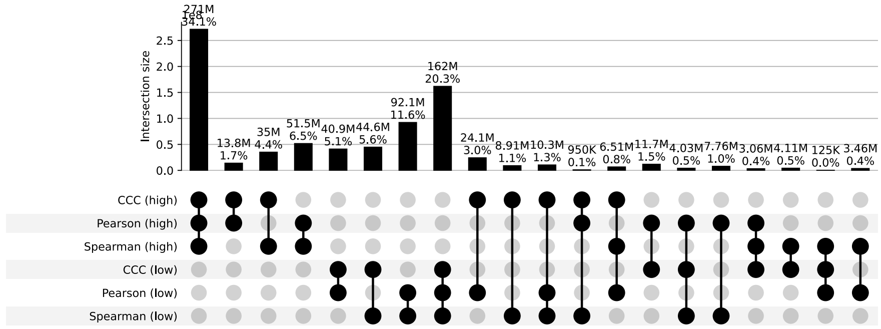


**d)** UpSet plot using permutation-based statistical thresholds

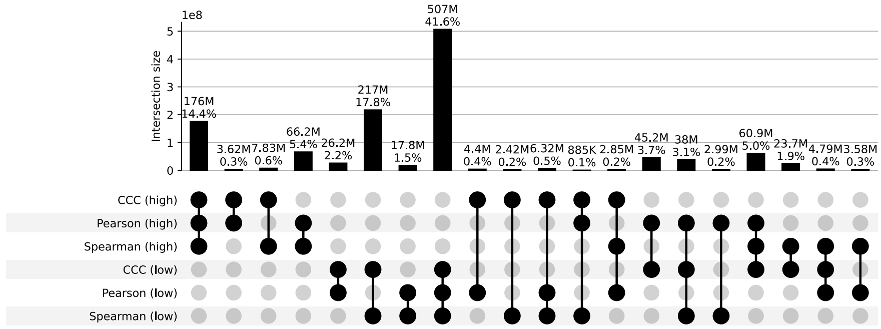


**Figure S12: Distribution and UpSet plots for GTEx v8 brain cerebellar hemisphere.**

#### Brain Cerebellum

**a)** Correlation coefficient distributions between gene pairs within GTEx v8 Brain Cerebellum

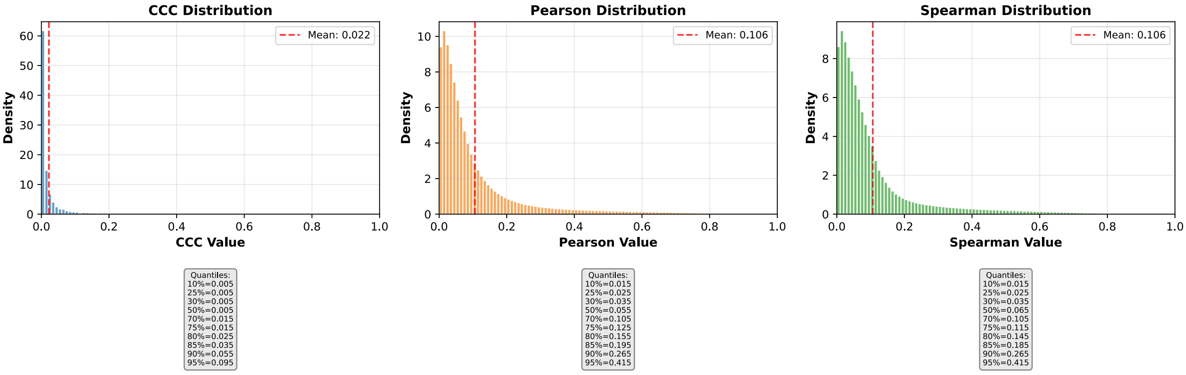


**b)** Corresponding cumulative histogram

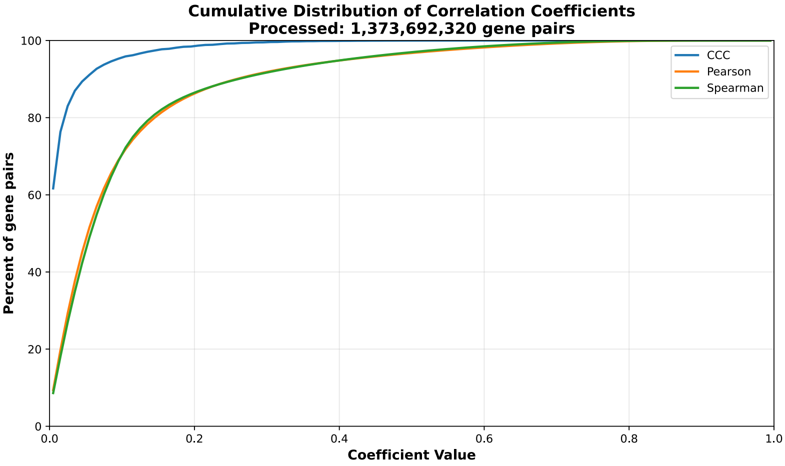


**c)** UpSet plot using top and bottom 30% correlations

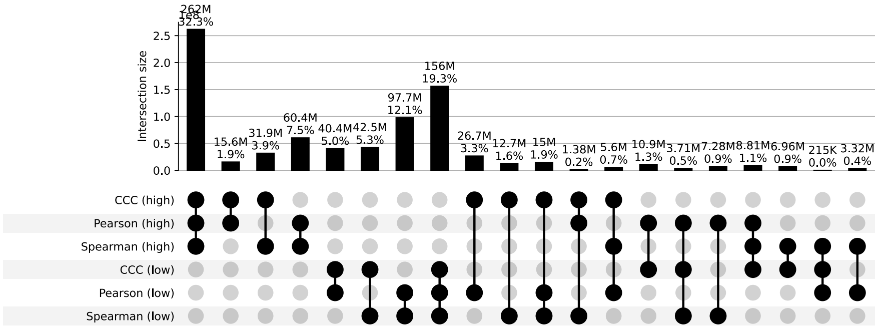


**d)** UpSet plot using permutation-based statistical thresholds

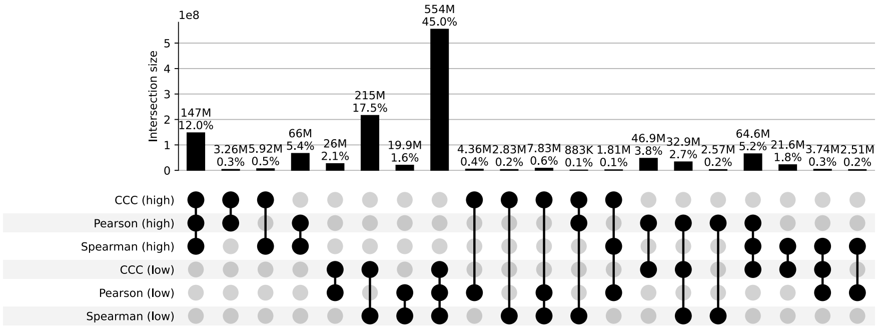


**Figure S13: Distribution and UpSet plots for GTEx v8 brain cerebellum.**

#### Brain Cortex

**a)** Correlation coefficient distributions between gene pairs within GTEx v8 Brain Cortex

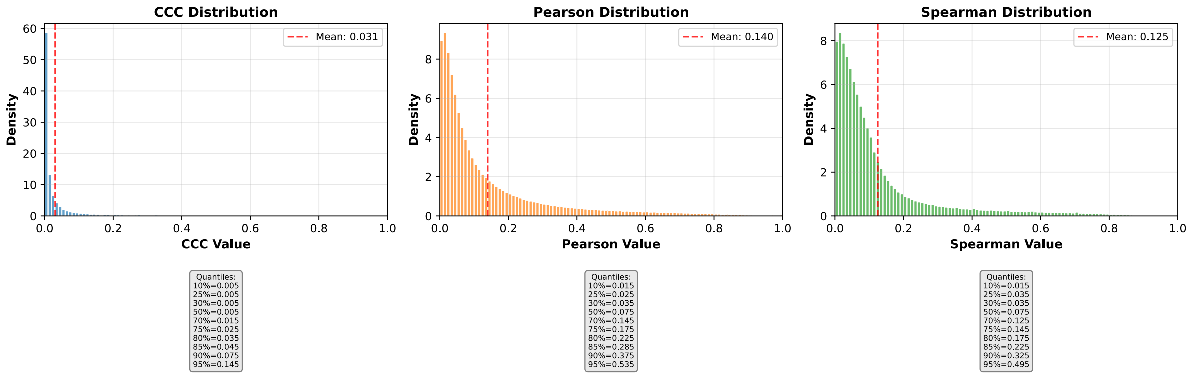


**b)** Corresponding cumulative histogram

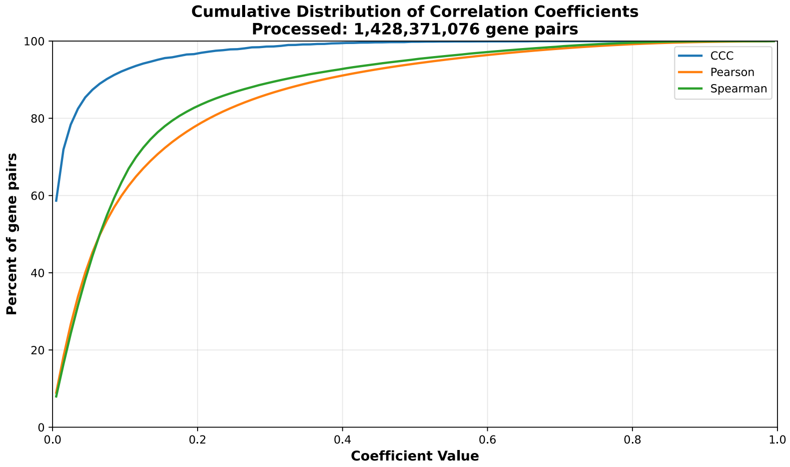


**c)** UpSet plot using top and bottom 30% correlations

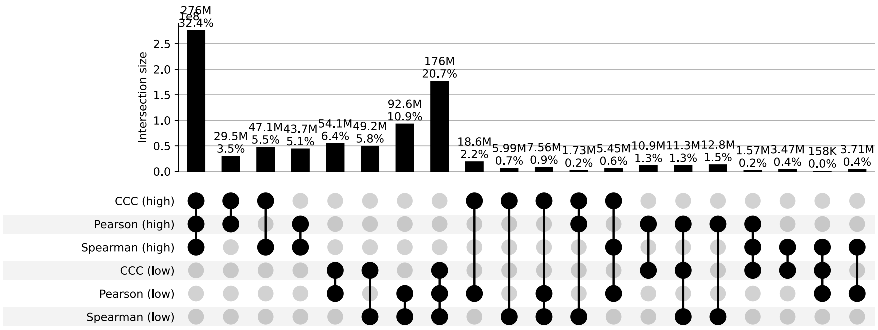


**d)** UpSet plot using permutation-based statistical thresholds

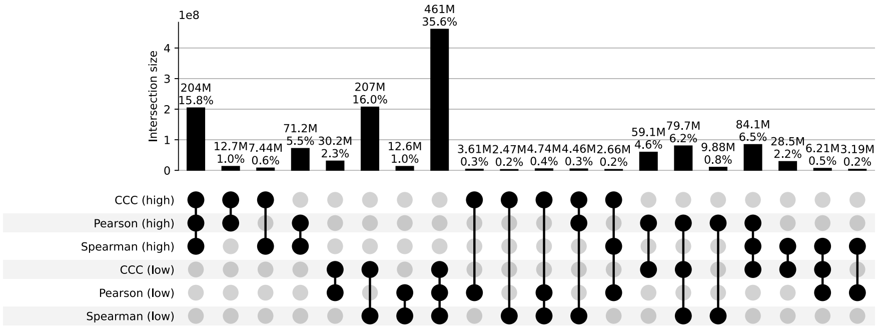


**Figure S14: Distribution and UpSet plots for GTEx v8 brain cortex.**

#### Brain Frontal Cortex Ba9

**a)** Correlation coefficient distributions between gene pairs within GTEx v8 Brain Frontal Cortex Ba9

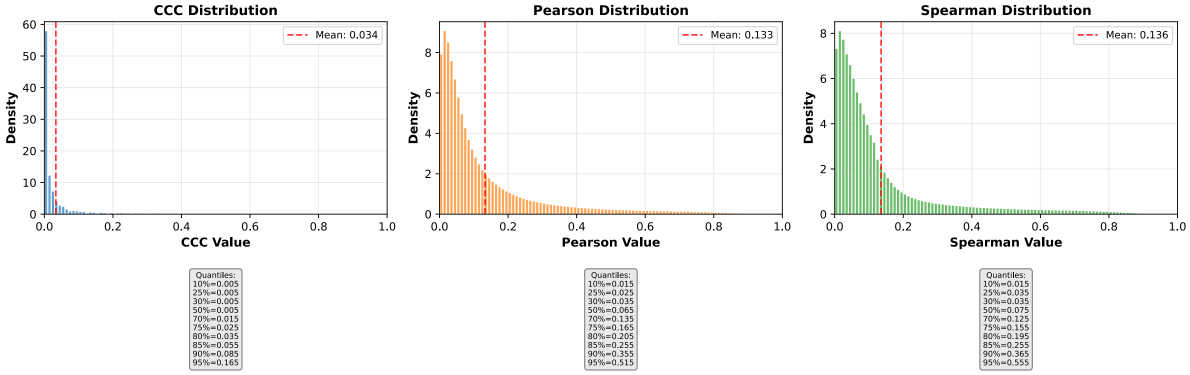


**b)** Corresponding cumulative histogram

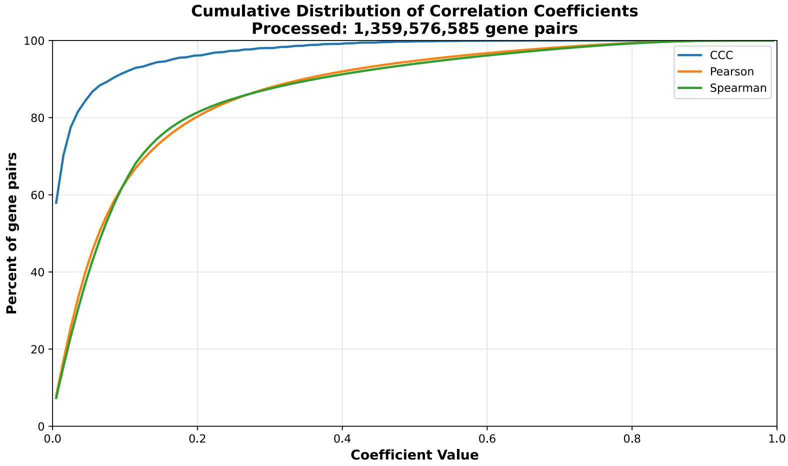


**c)** UpSet plot using top and bottom 30% correlations

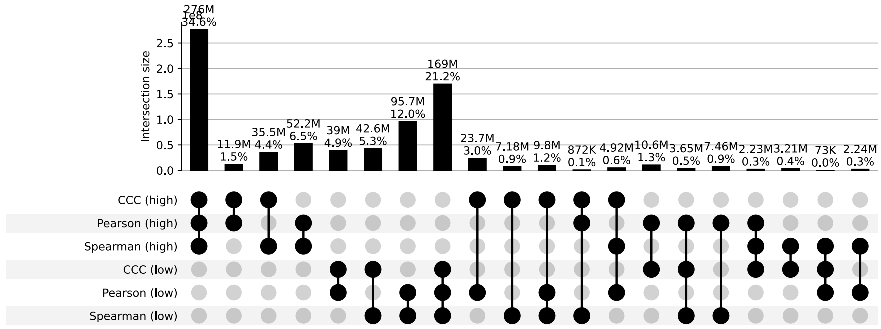


**d)** UpSet plot using permutation-based statistical thresholds

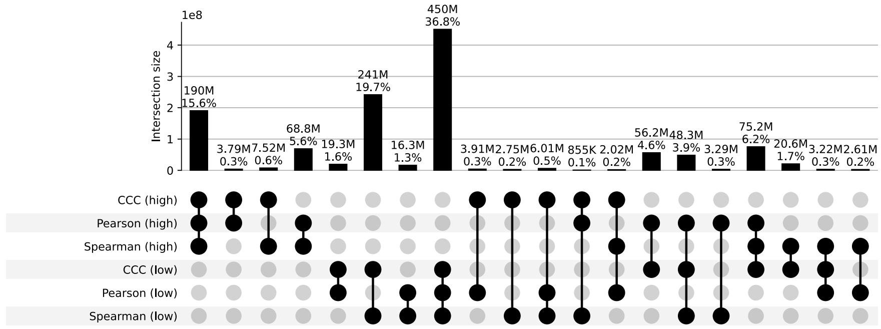


**Figure S15: Distribution and UpSet plots for GTEx v8 brain frontal cortex BA9.**

#### Brain Hippocampus

**a)** Correlation coefficient distributions between gene pairs within GTEx v8 Brain Hippocampus

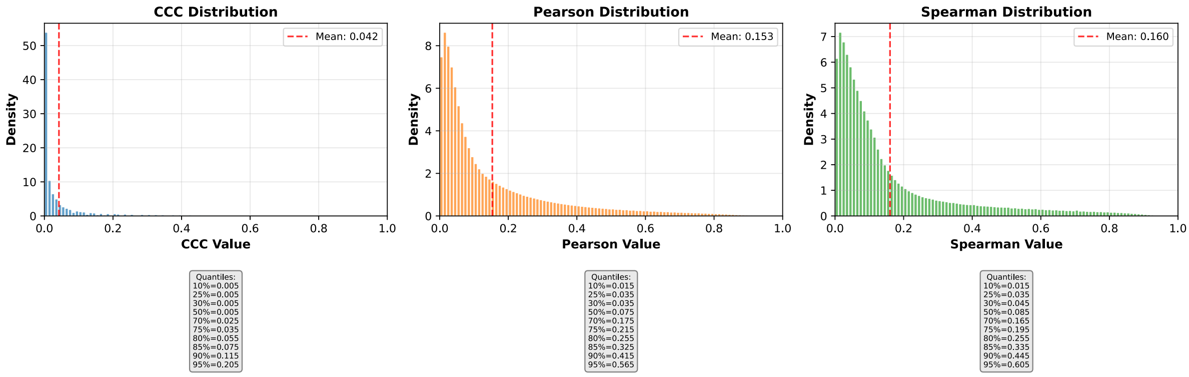


**b)** Corresponding cumulative histogram

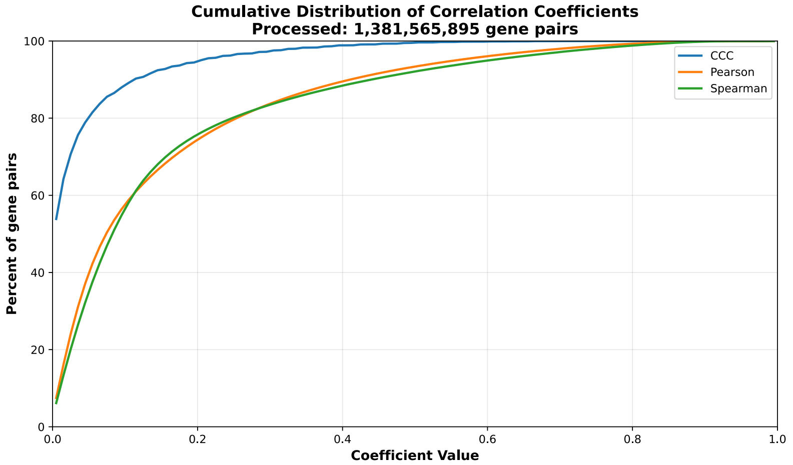


**c)** UpSet plot using top and bottom 30% correlations

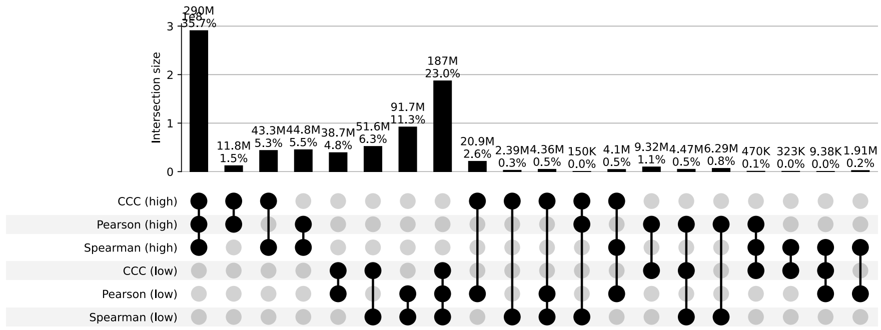


**d)** UpSet plot using permutation-based statistical thresholds

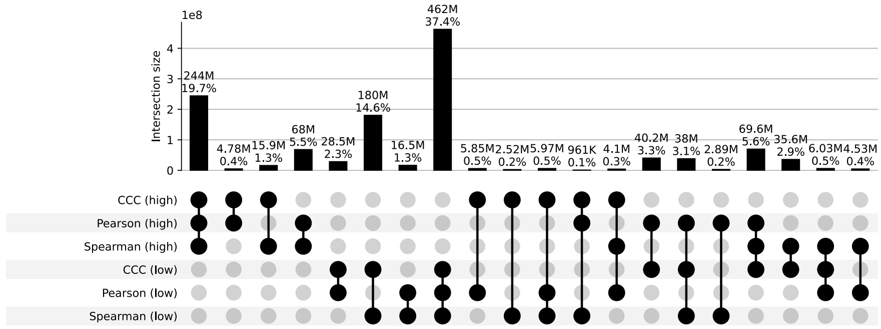


**Figure S16: Distribution and UpSet plots for GTEx v8 brain hippocampus.**

#### Brain Hypothalamus

**a)** Correlation coefficient distributions between gene pairs within GTEx v8 Brain Hypothalamus

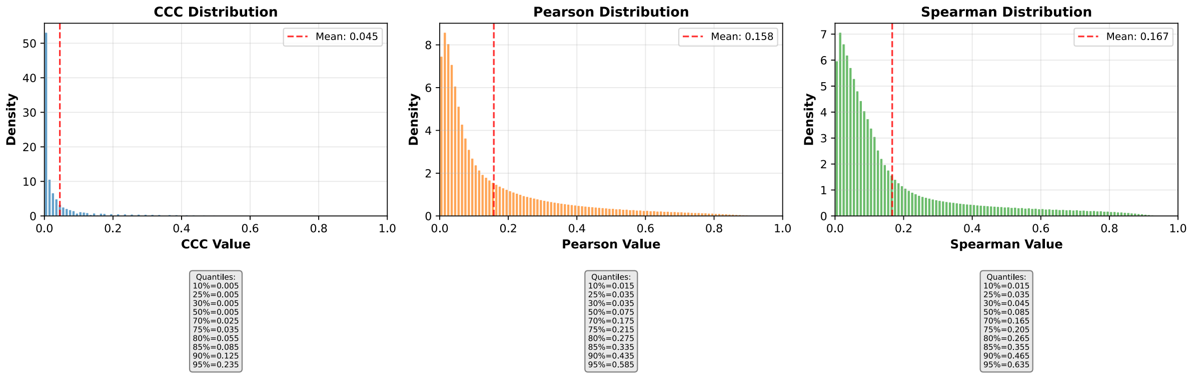


**b)** Corresponding cumulative histogram

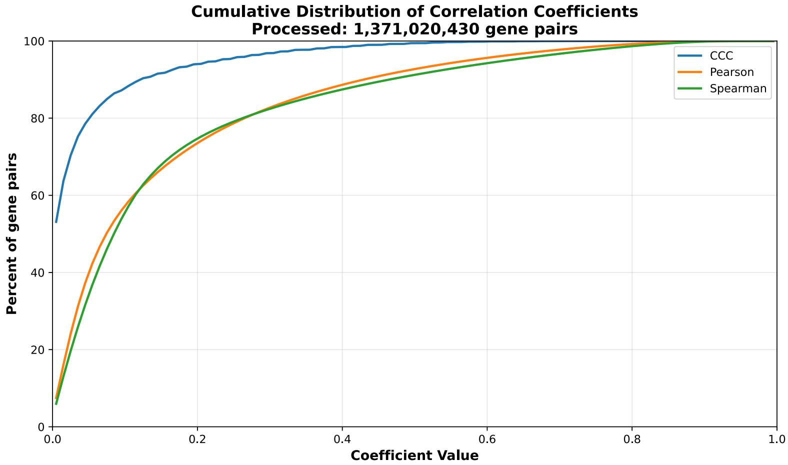


**c)** UpSet plot using top and bottom 30% correlations

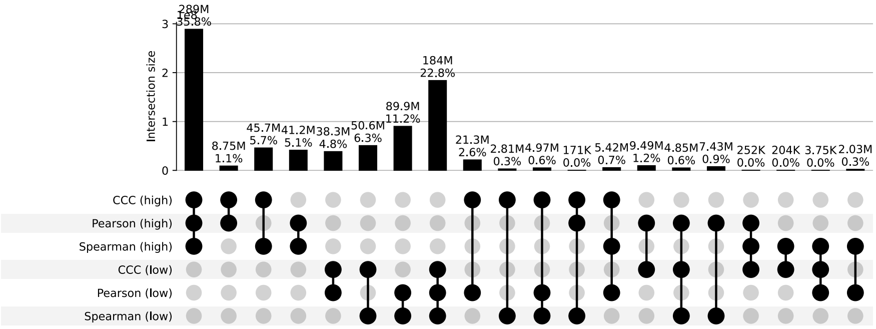


**d)** UpSet plot using permutation-based statistical thresholds

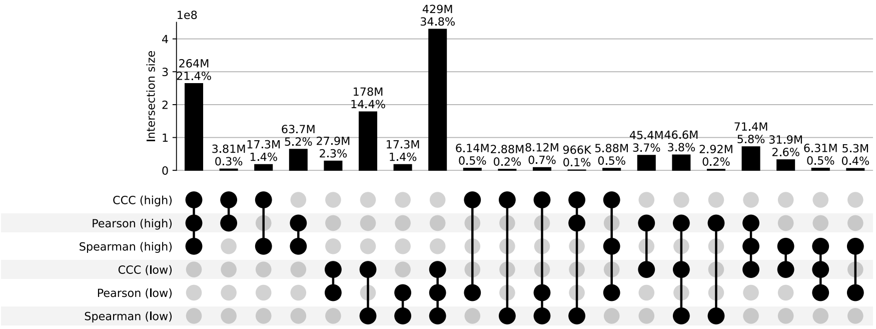


**Figure S17: Distribution and UpSet plots for GTEx v8 brain hypothalamus.**

#### Brain Nucleus Accumbens Basal Ganglia

**a)** Correlation coefficient distributions between gene pairs within GTEx v8 Brain Nucleus Accumbens Basal Ganglia

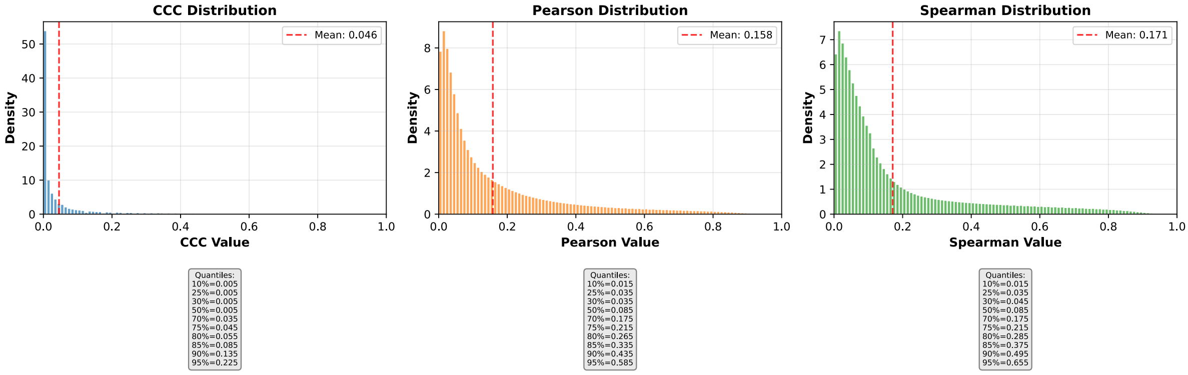


**b)** Corresponding cumulative histogram

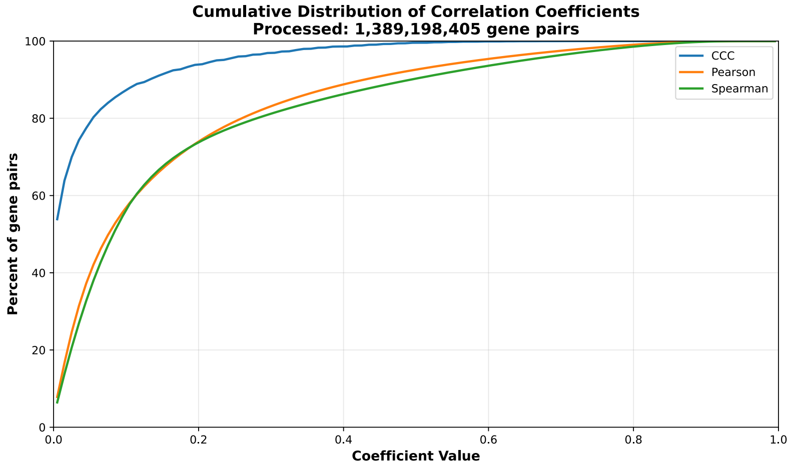


**c)** UpSet plot using top and bottom 30% correlations

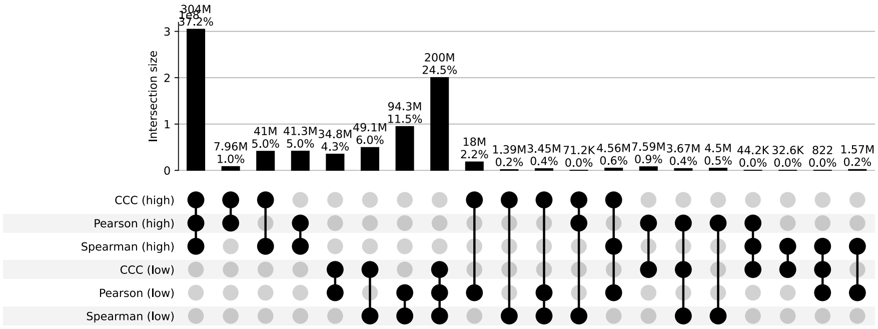


**d)** UpSet plot using permutation-based statistical thresholds

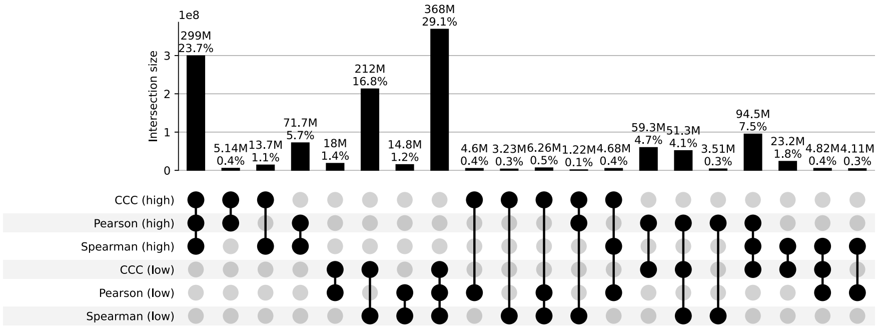


**Figure S18: Distribution and UpSet plots for GTEx v8 brain nucleus accumbens basal ganglia.**

#### Brain Putamen Basal Ganglia

**a)** Correlation coefficient distributions between gene pairs within GTEx v8 Brain Putamen Basal Ganglia

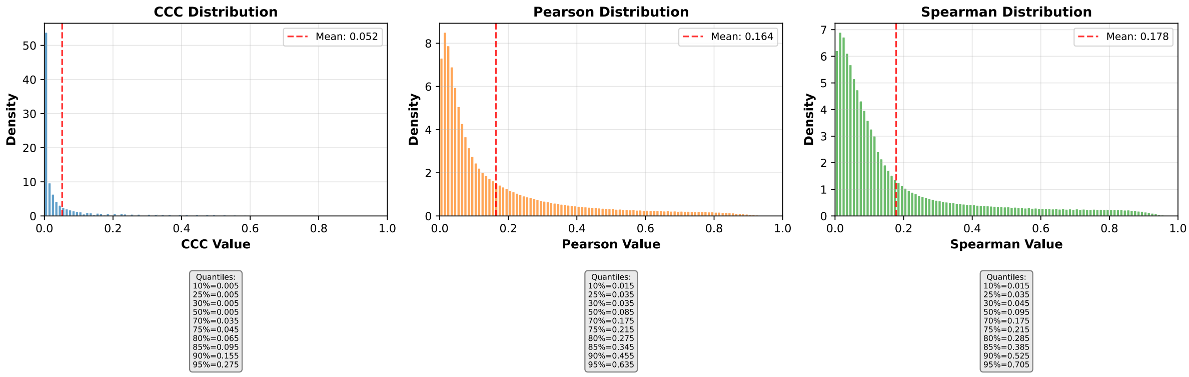


**b)** Corresponding cumulative histogram

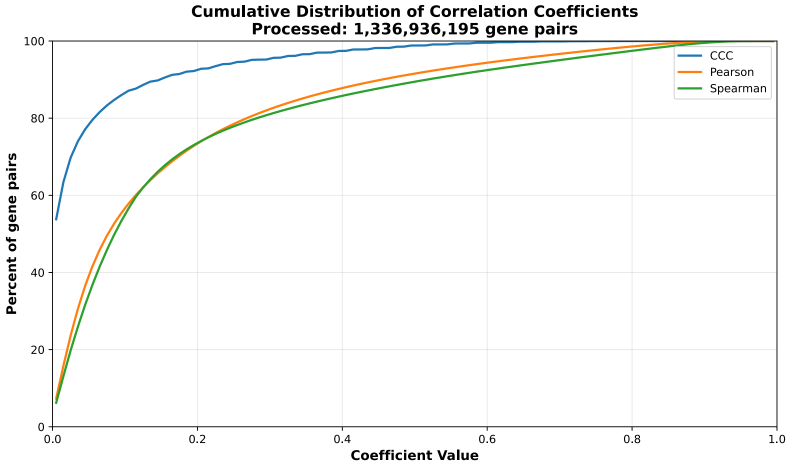


**c)** UpSet plot using top and bottom 30% correlations

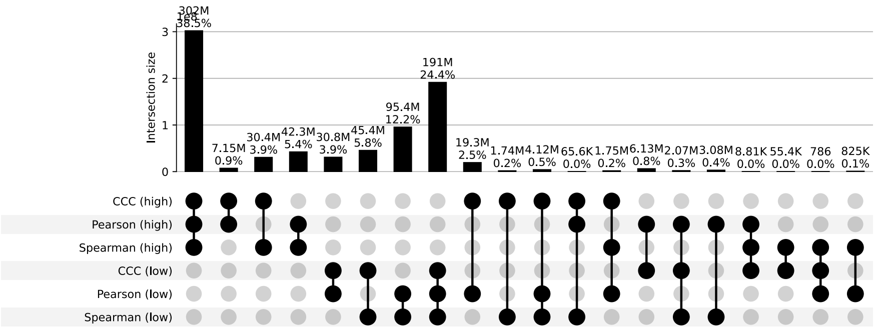


**d)** UpSet plot using permutation-based statistical thresholds

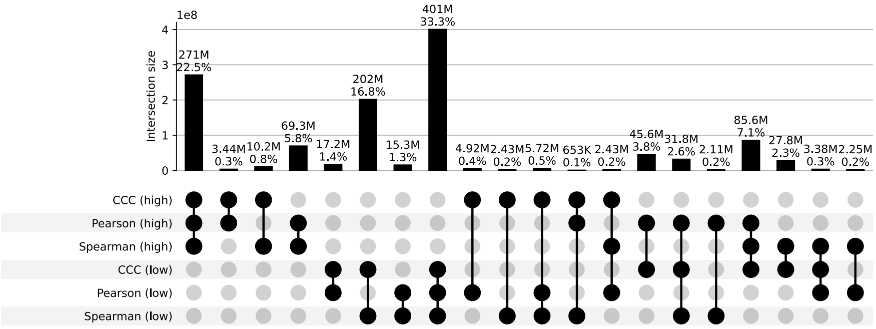


**Figure S19: Distribution and UpSet plots for GTEx v8 brain putamen basal ganglia.**

#### Brain Spinal Cord Cervical C1

**a)** Correlation coefficient distributions between gene pairs within GTEx v8 Brain Spinal Cord Cervical C1

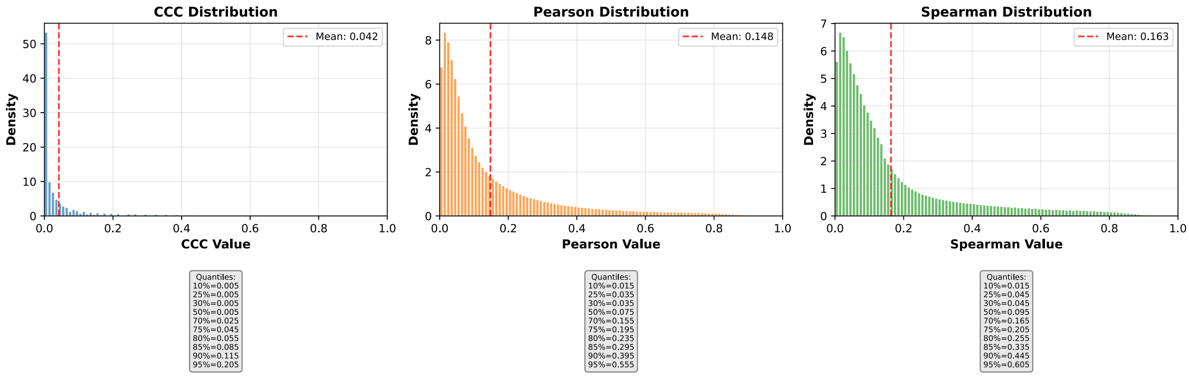


**b)** Corresponding cumulative histogram

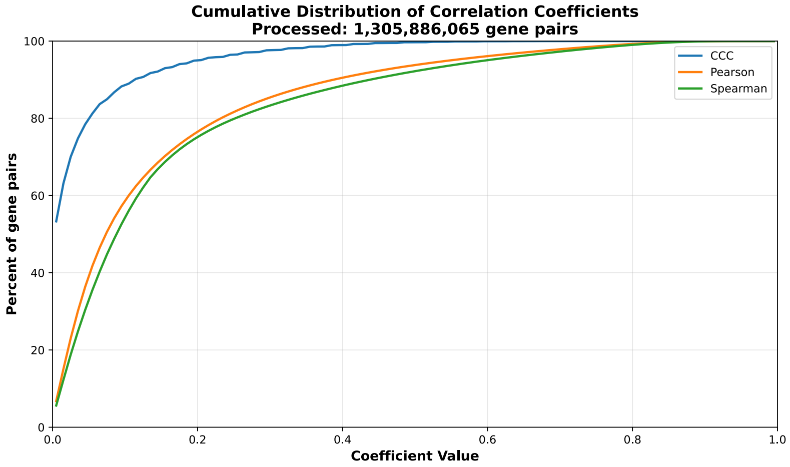


**c)** UpSet plot using top and bottom 30% correlations

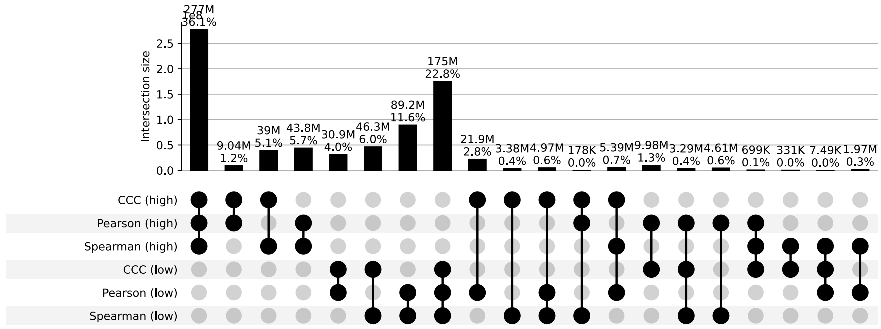


**d)** UpSet plot using permutation-based statistical thresholds

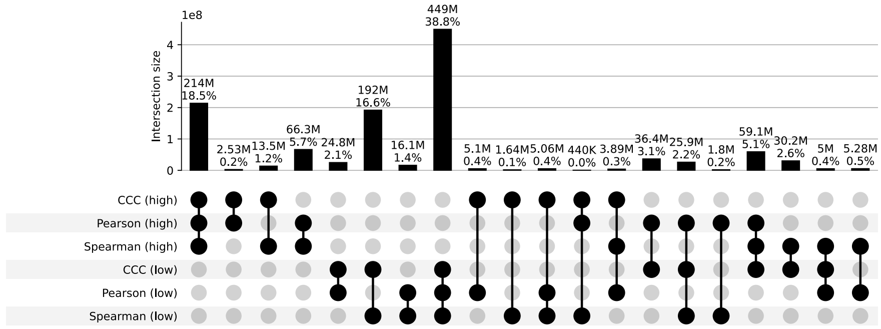


**Figure S20: Distribution and UpSet plots for GTEx v8 brain spinal cord cervical C1.**

#### Brain Substantia Nigra

**a)** Correlation coefficient distributions between gene pairs within GTEx v8 Brain Substantia Nigra

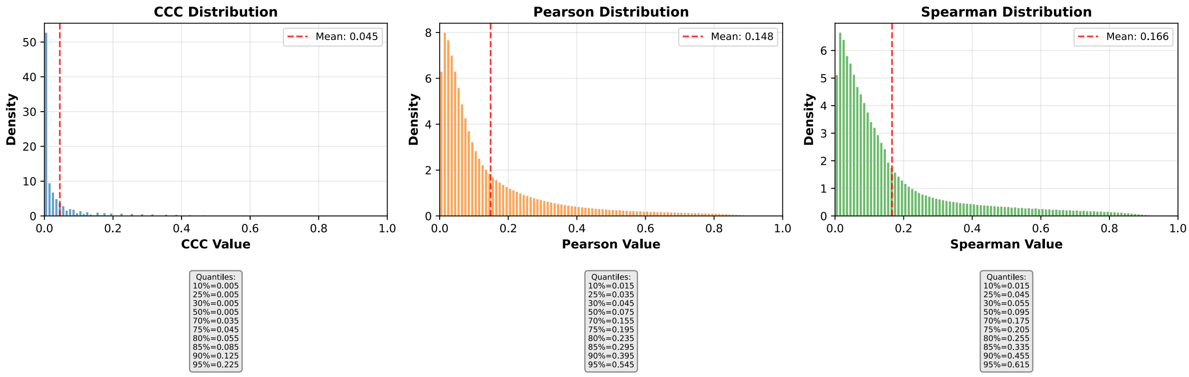


**b)** Corresponding cumulative histogram

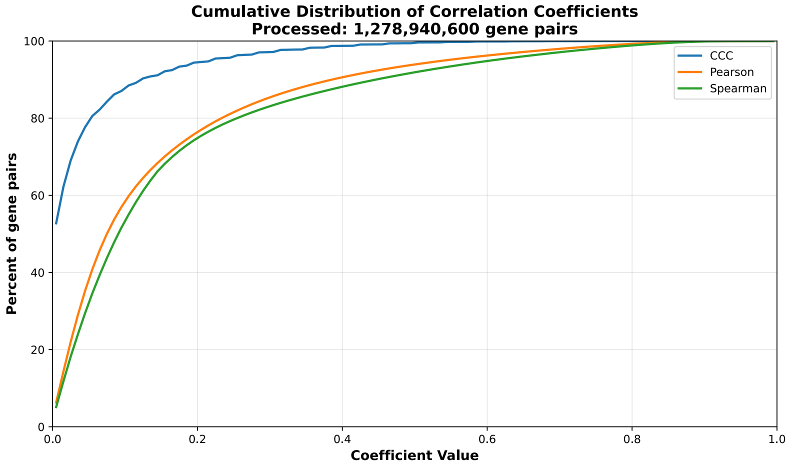


**c)** UpSet plot using top and bottom 30% correlations

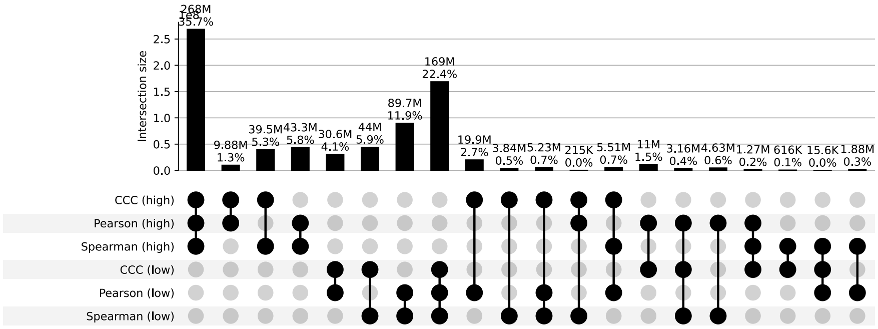


**d)** UpSet plot using permutation-based statistical thresholds

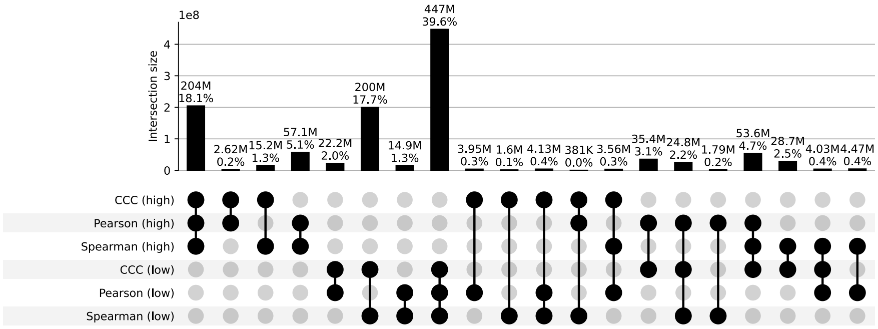


**Figure S21: Distribution and UpSet plots for GTEx v8 brain substantia nigra.**

#### Breast Mammary Tissue

**a)** Correlation coefficient distributions between gene pairs within GTEx v8 Breast Mammary Tissue

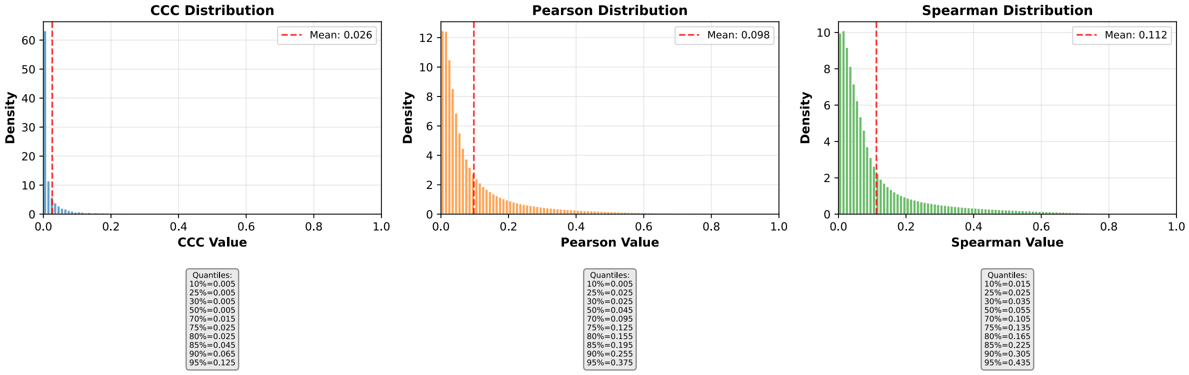


**b)** Corresponding cumulative histogram

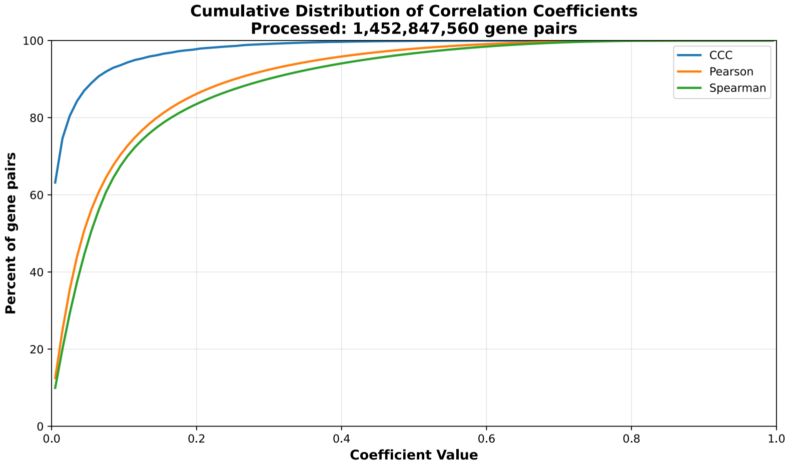


**c)** UpSet plot using top and bottom 30% correlations

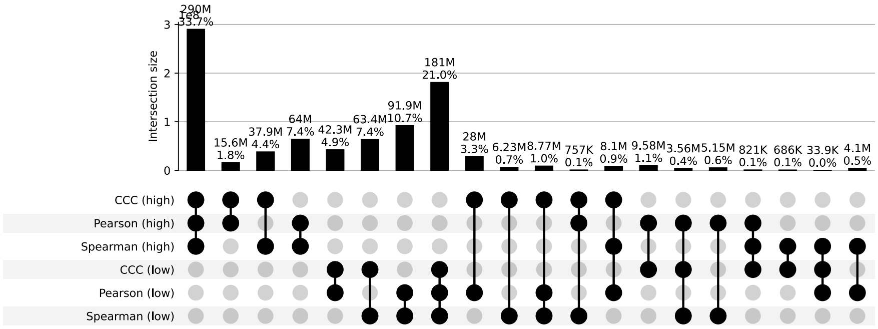


**d)** UpSet plot using permutation-based statistical thresholds

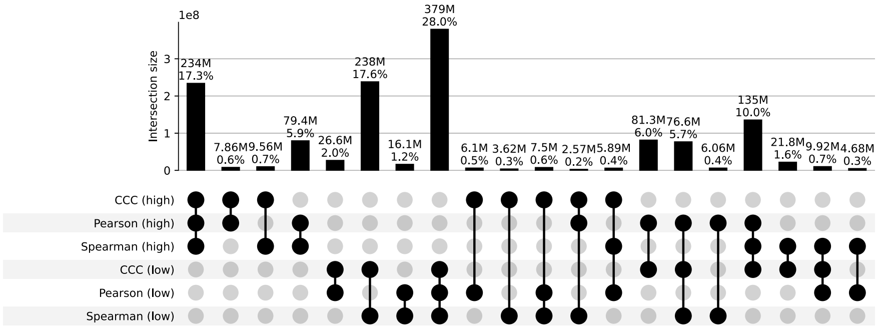


**Figure S22: Distribution and UpSet plots for GTEx v8 breast mammary tissue.**

#### Cells Cultured Fibroblasts

**a)** Correlation coefficient distributions between gene pairs within GTEx v8 Cells Cultured Fibroblasts

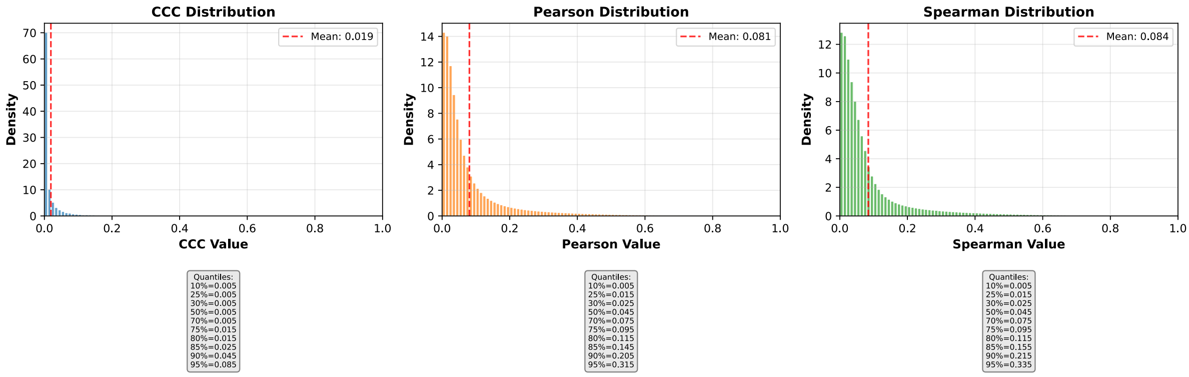


**b)** Corresponding cumulative histogram

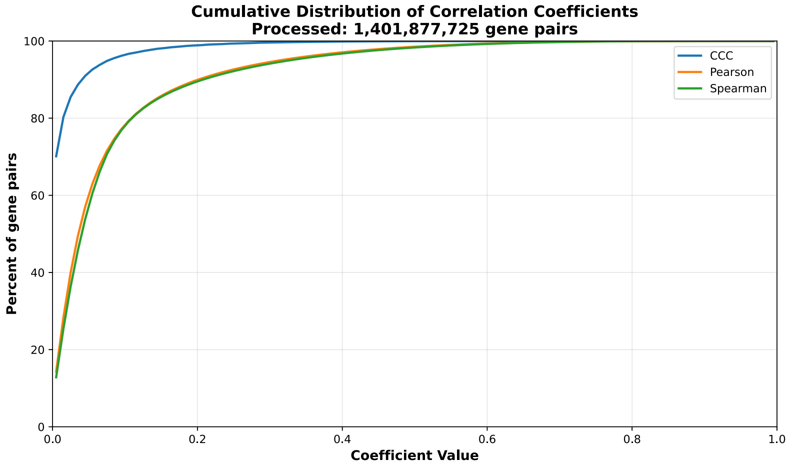


**c)** UpSet plot using top and bottom 30% correlations

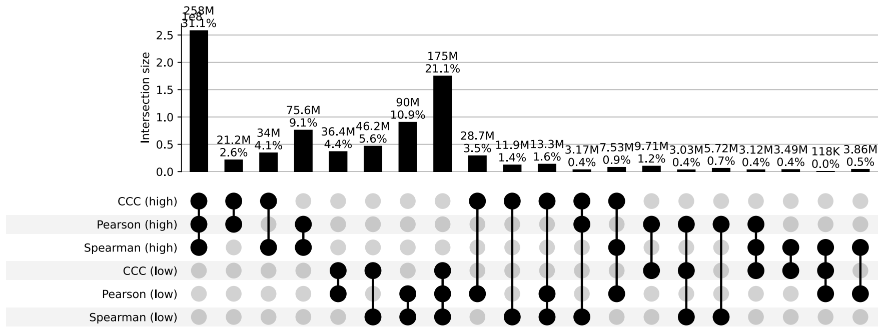


**d)** UpSet plot using permutation-based statistical thresholds

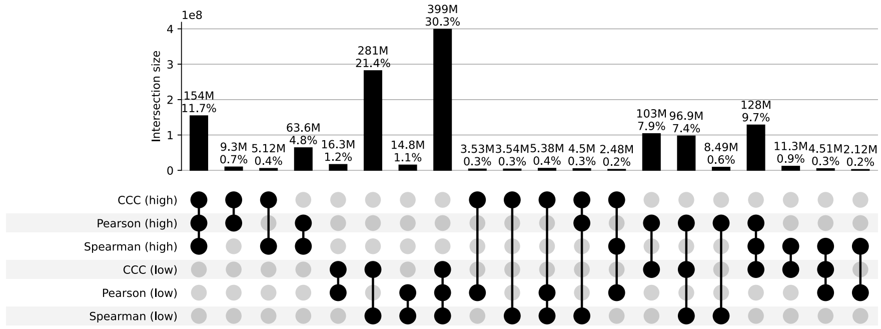


**Figure S23: Distribution and UpSet plots for GTEx v8 cells cultured fibroblasts.**

#### Cells Ebvtransformed Lymphocytes

**a)** Correlation coefficient distributions between gene pairs within GTEx v8 Cells Ebvtransformed Lymphocytes

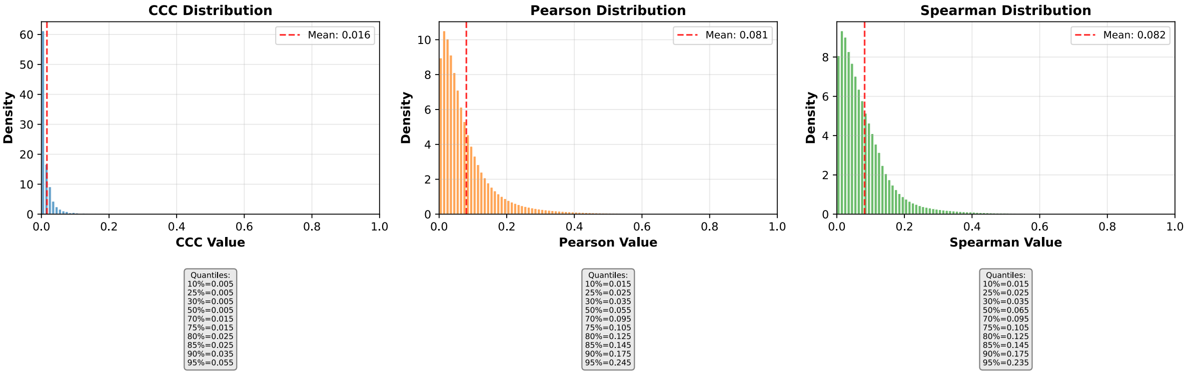


**b)** Corresponding cumulative histogram

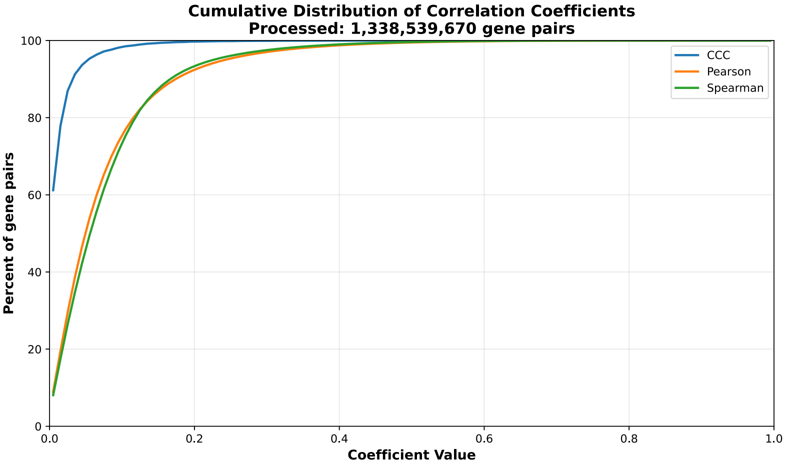


**c)** UpSet plot using top and bottom 30% correlations

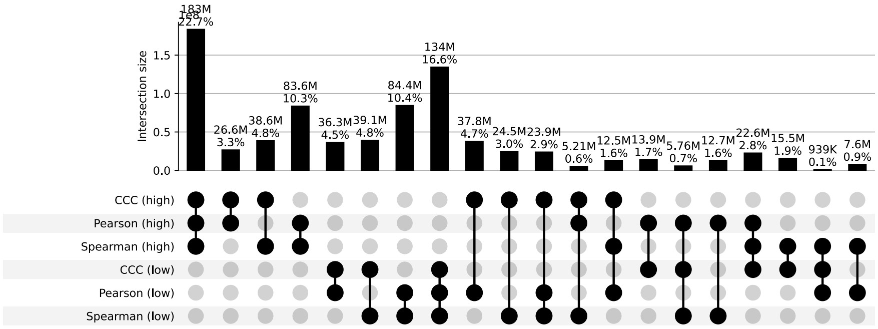


**d)** UpSet plot using permutation-based statistical thresholds

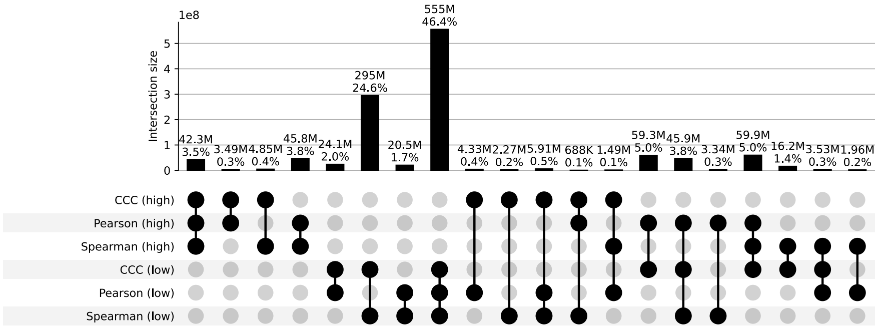


**Figure S24: Distribution and UpSet plots for GTEx v8 cells EBV-transformed lymphocytes.**

#### Cervix Ectocervix

**a)** Correlation coefficient distributions between gene pairs within GTEx v8 Cervix Ectocervix

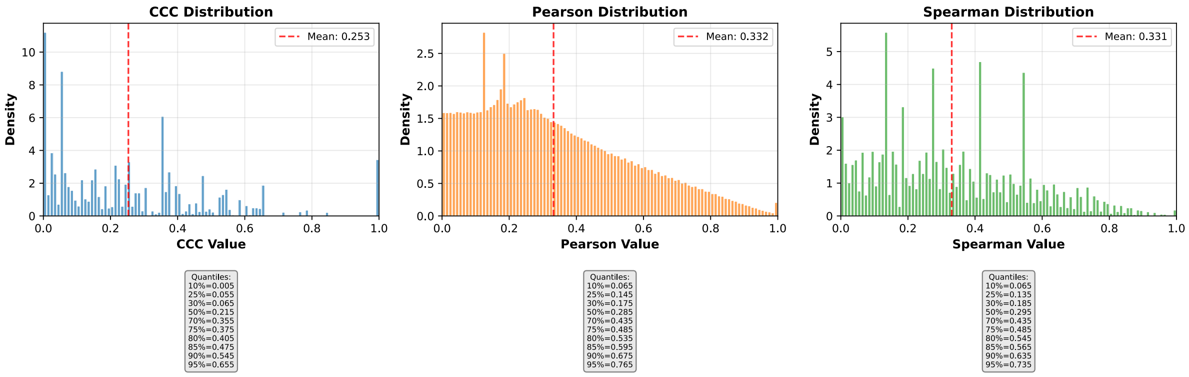


**b)** Corresponding cumulative histogram

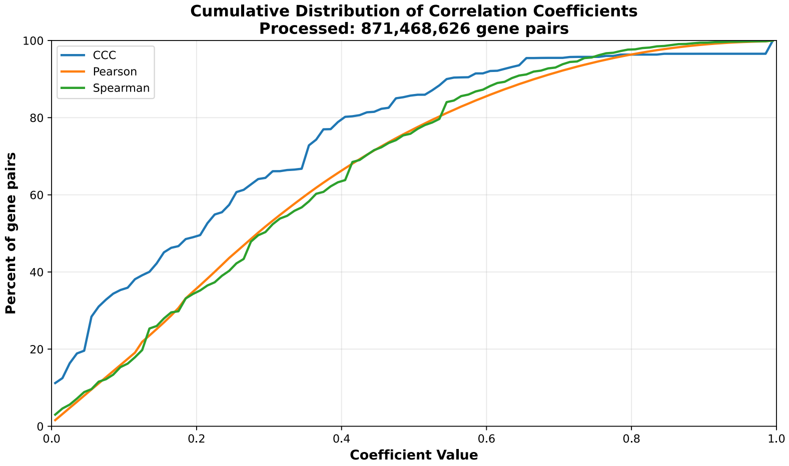


**c)** UpSet plot using top and bottom 30% correlations

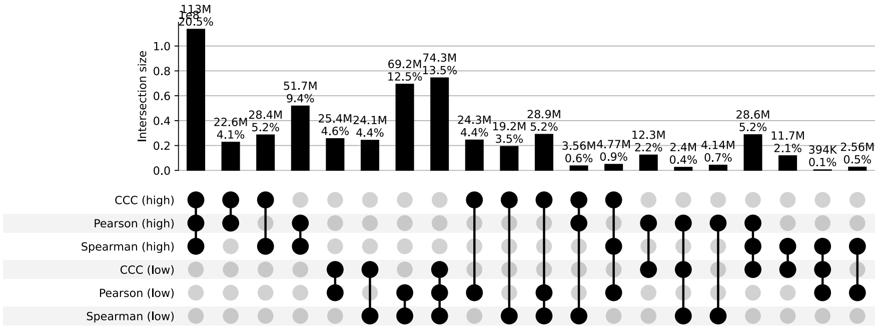


**d)** UpSet plot using permutation-based statistical thresholds

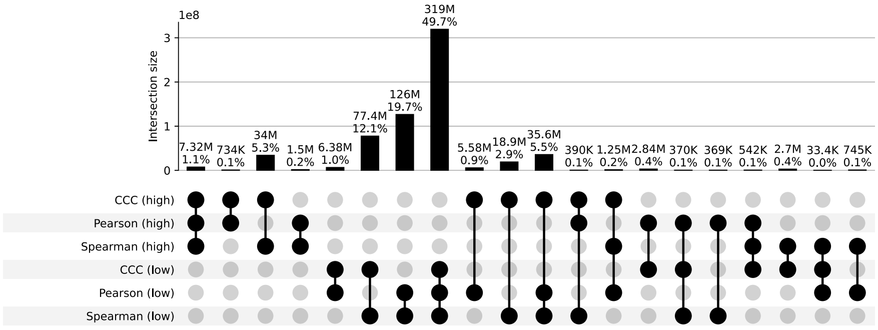


**Figure S25: Distribution and UpSet plots for GTEx v8 cervix ectocervix.**

#### Cervix Endocervix

**a)** Correlation coefficient distributions between gene pairs within GTEx v8 Cervix Endocervix

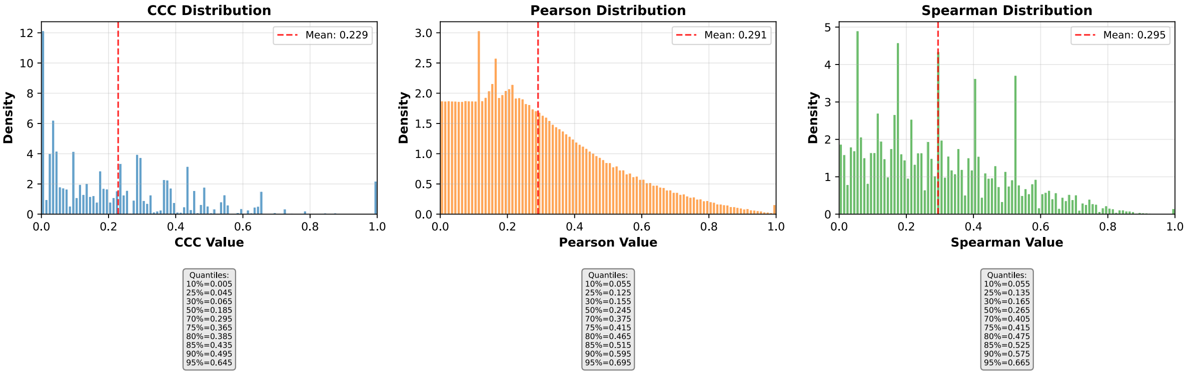


**b)** Corresponding cumulative histogram

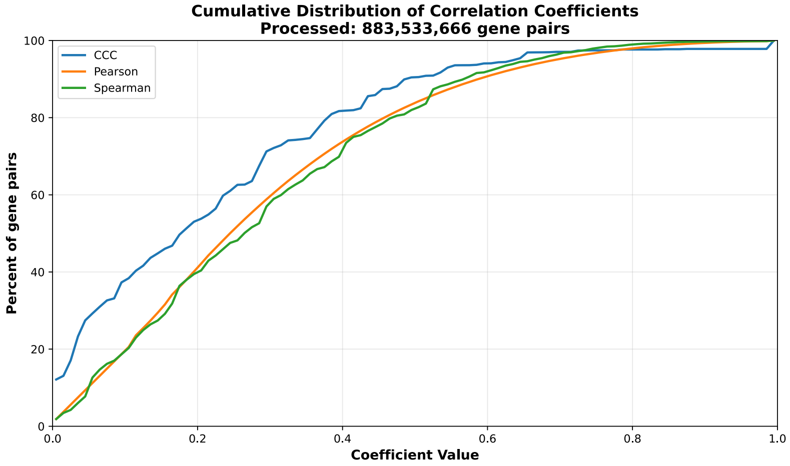


**c)** UpSet plot using top and bottom 30% correlations

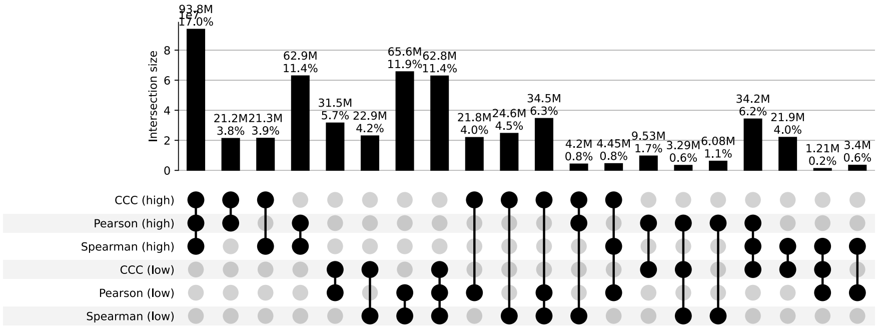


**d)** UpSet plot using permutation-based statistical thresholds

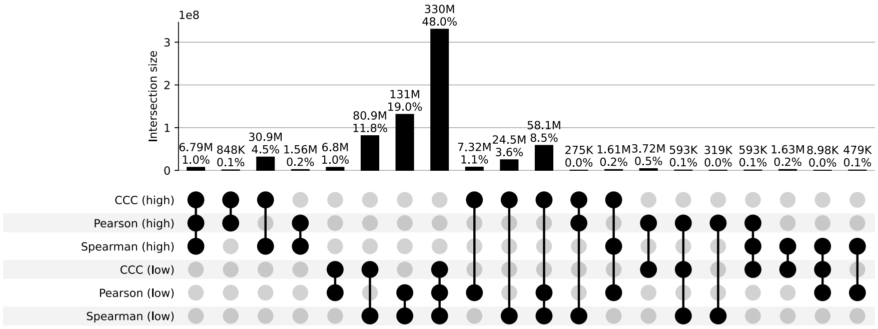


**Figure S26: Distribution and UpSet plots for GTEx v8 cervix endocervix.**

#### Colon Sigmoid

**a)** Correlation coefficient distributions between gene pairs within GTEx v8 Colon Sigmoid

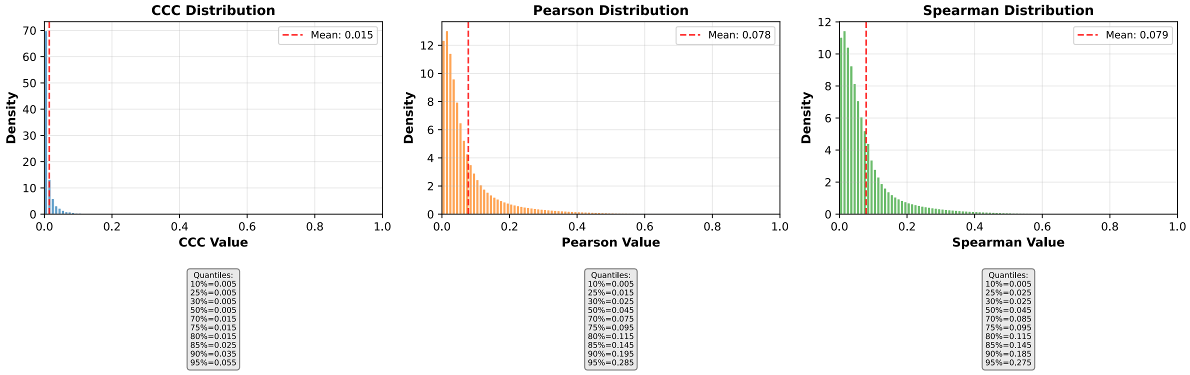


**b)** Corresponding cumulative histogram

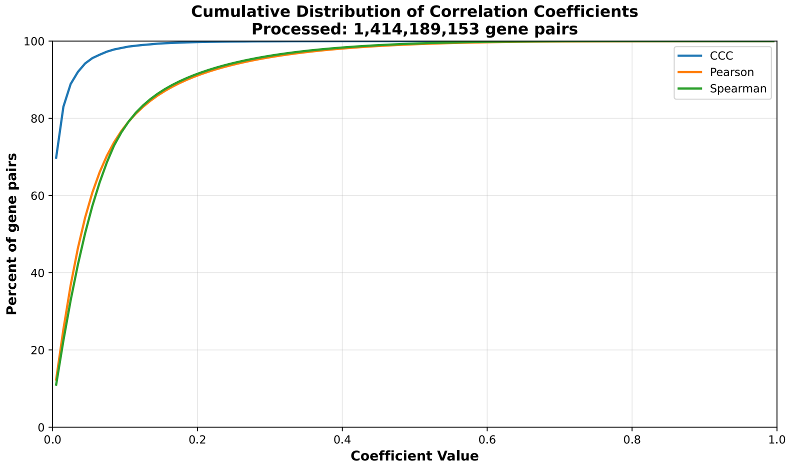


**c)** UpSet plot using top and bottom 30% correlations

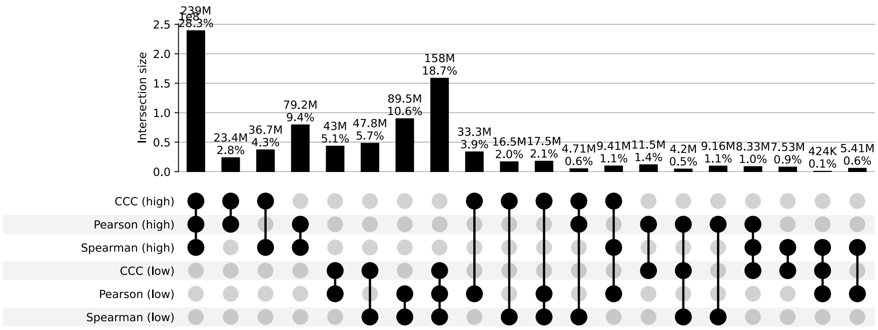


**d)** UpSet plot using permutation-based statistical thresholds

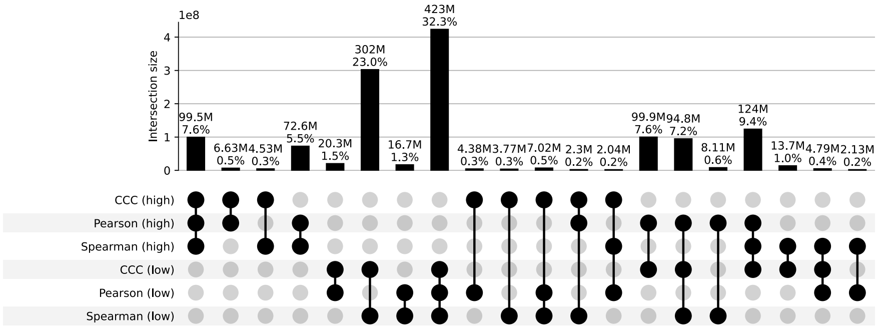


**Figure S27: Distribution and UpSet plots for GTEx v8 colon sigmoid.**

#### Colon Transverse

**a)** Correlation coefficient distributions between gene pairs within GTEx v8 Colon Transverse

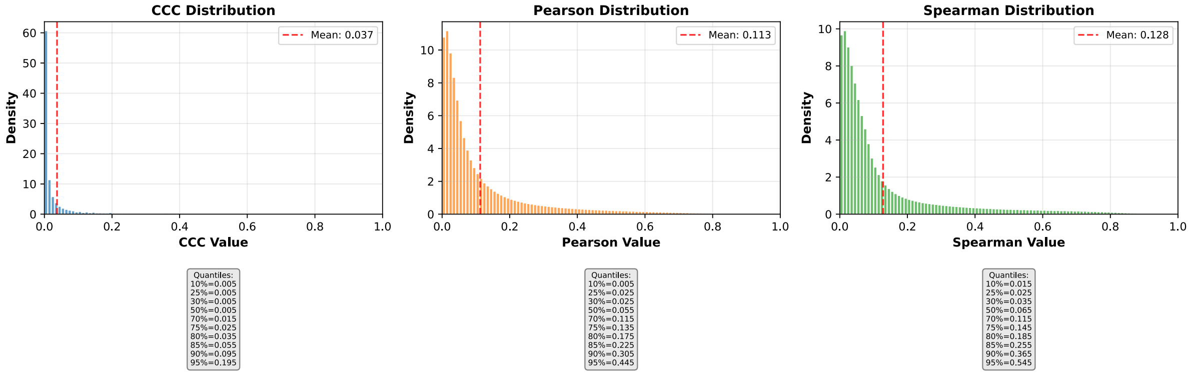


**b)** Corresponding cumulative histogram

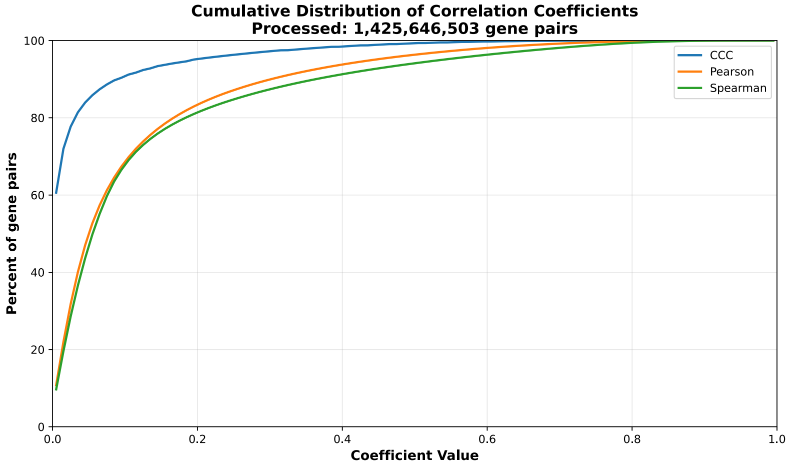


**c)** UpSet plot using top and bottom 30% correlations

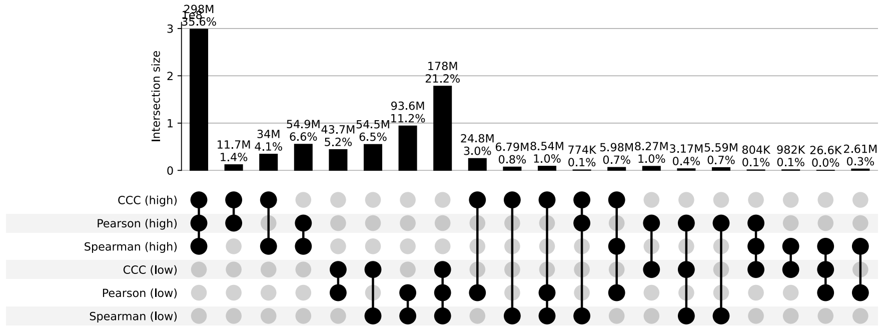


**d)** UpSet plot using permutation-based statistical thresholds

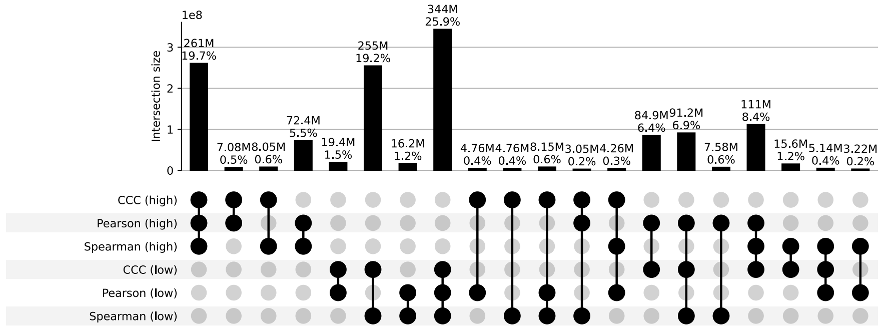


**Figure S28: Distribution and UpSet plots for GTEx v8 colon transverse.**

#### Esophagus Gastroesophageal Junction

**a)** Correlation coefficient distributions between gene pairs within GTEx v8 Esophagus Gastroesophageal Junction

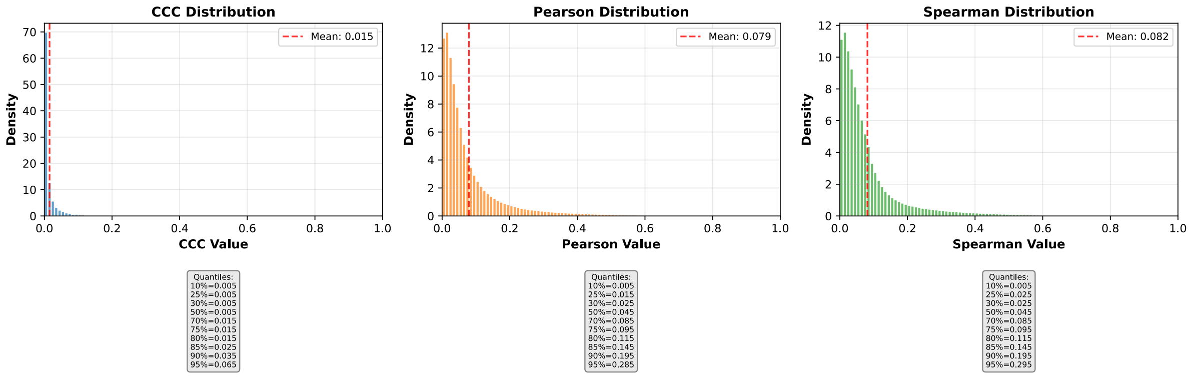


**b)** Corresponding cumulative histogram

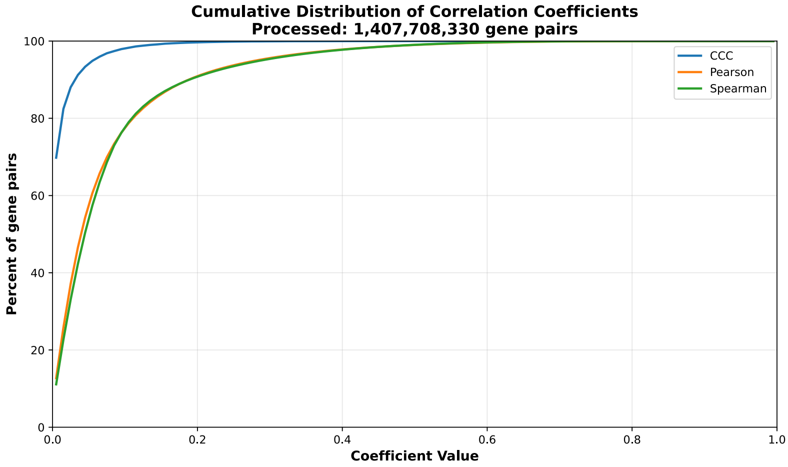


**c)** UpSet plot using top and bottom 30% correlations

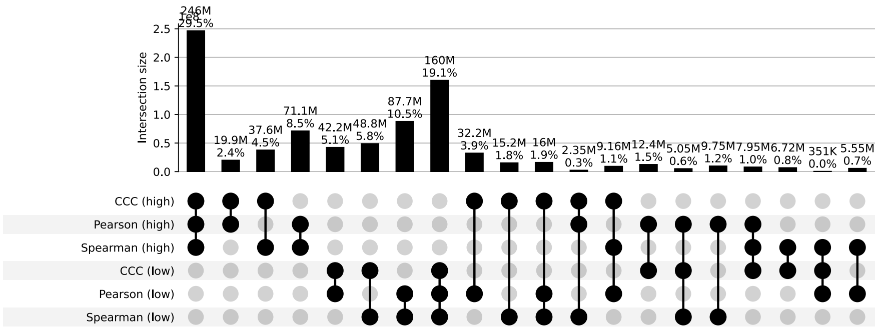


**d)** UpSet plot using permutation-based statistical thresholds

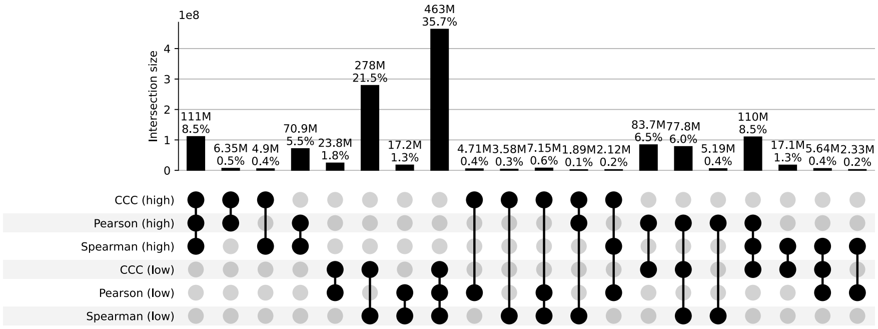


**Figure S29: Distribution and UpSet plots for GTEx v8 esophagus gastroesophageal junction.**

#### Esophagus Mucosa

**a)** Correlation coefficient distributions between gene pairs within GTEx v8 Esophagus Mucosa

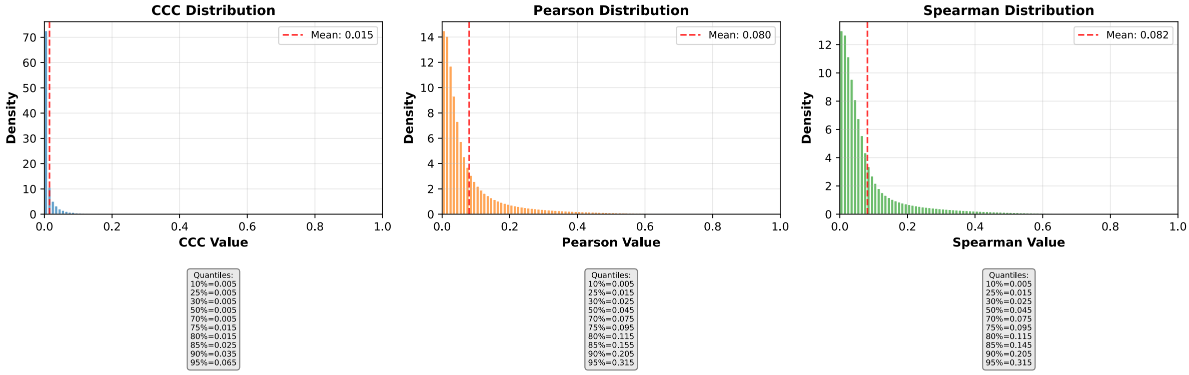


**b)** Corresponding cumulative histogram

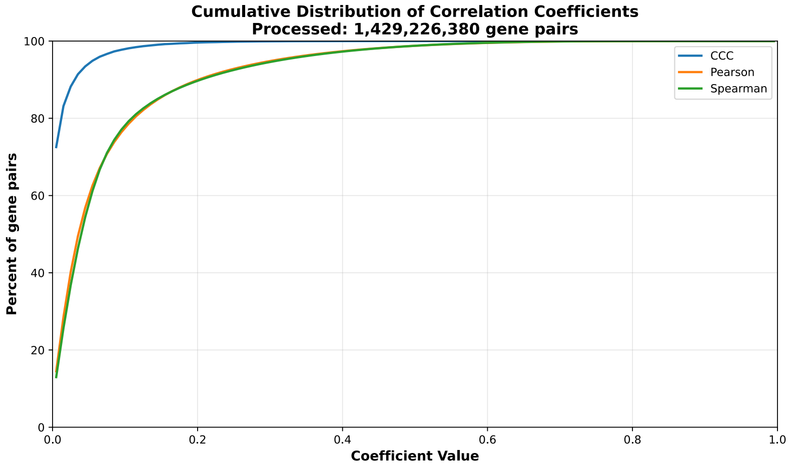


**c)** UpSet plot using top and bottom 30% correlations

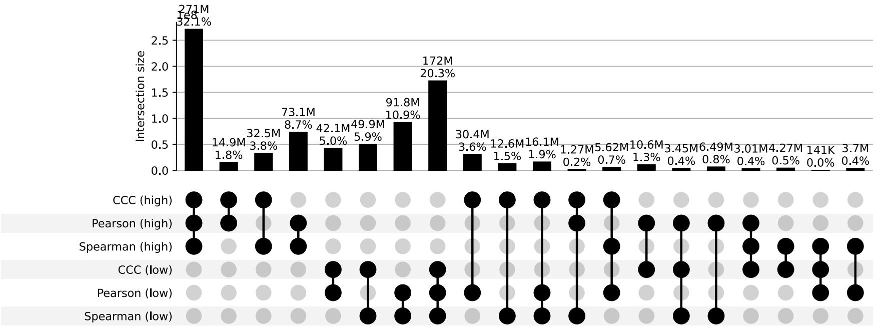


**d)** UpSet plot using permutation-based statistical thresholds

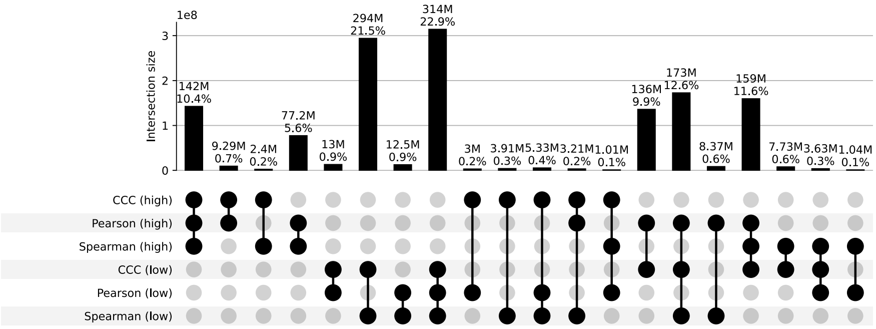


**Figure S30: Distribution and UpSet plots for GTEx v8 esophagus mucosa.**

#### Esophagus Muscularis

**a)** Correlation coefficient distributions between gene pairs within GTEx v8 Esophagus Muscularis

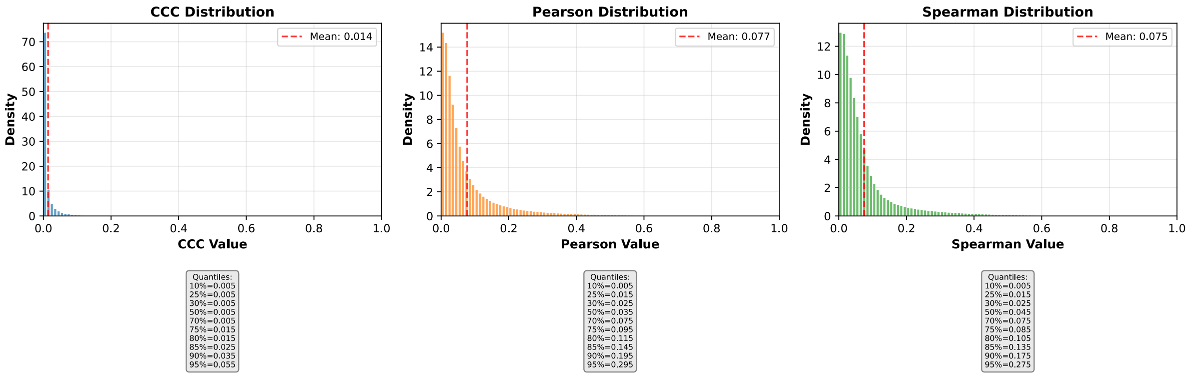


**b)** Corresponding cumulative histogram

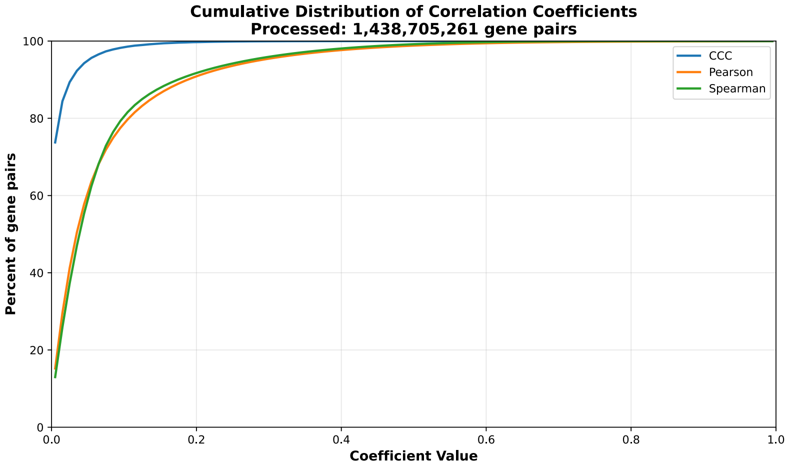


**c)** UpSet plot using top and bottom 30% correlations

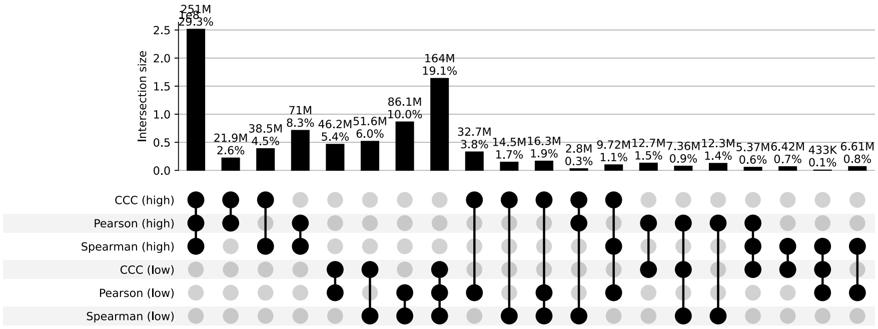


**d)** UpSet plot using permutation-based statistical thresholds

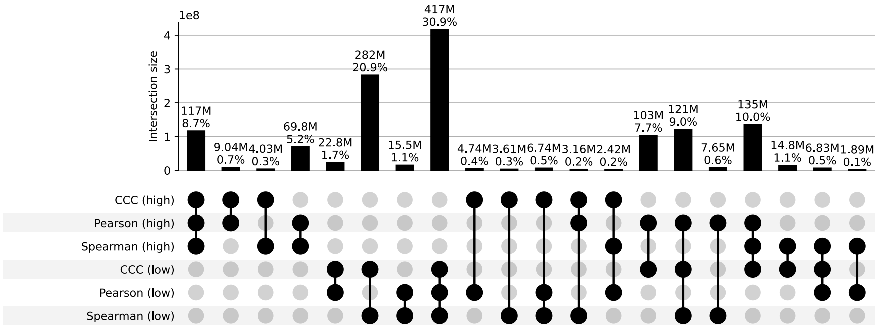


**Figure S31: Distribution and UpSet plots for GTEx v8 esophagus muscularis.**

#### Fallopian Tube

**a)** Correlation coefficient distributions between gene pairs within GTEx v8 Fallopian Tube

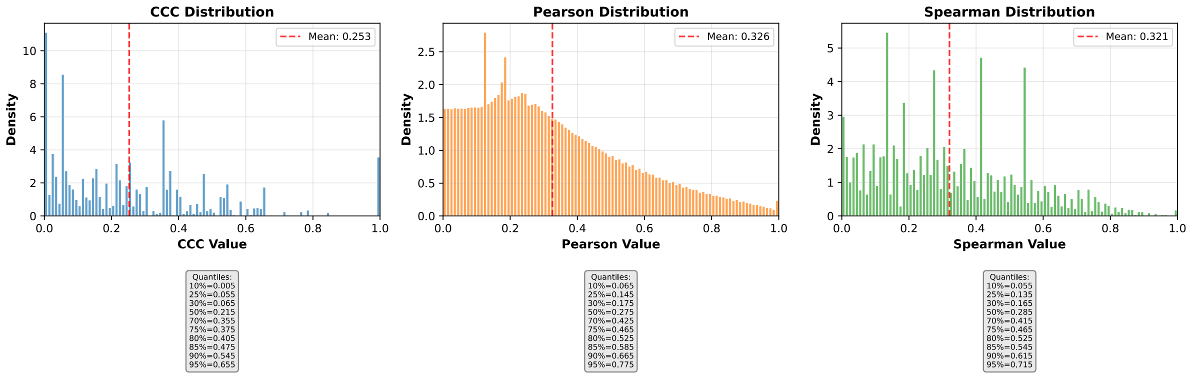


**b)** Corresponding cumulative histogram

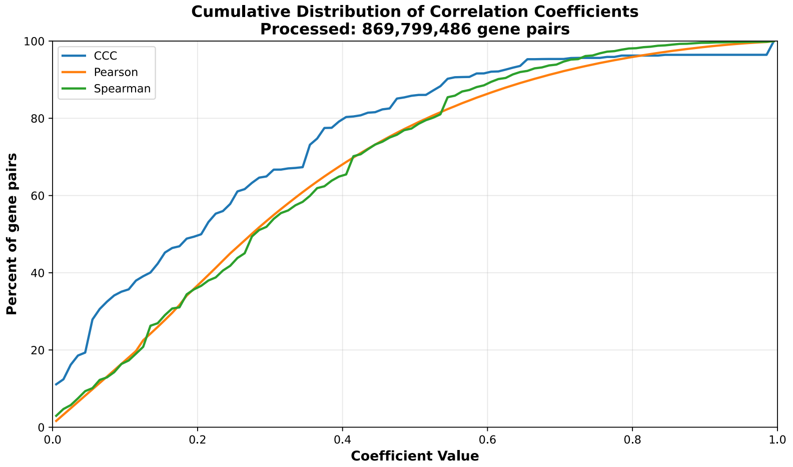


**c)** UpSet plot using top and bottom 30% correlations

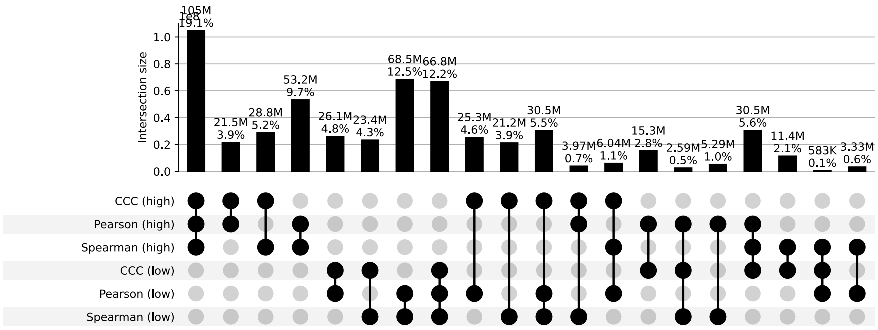


**d)** UpSet plot using permutation-based statistical thresholds

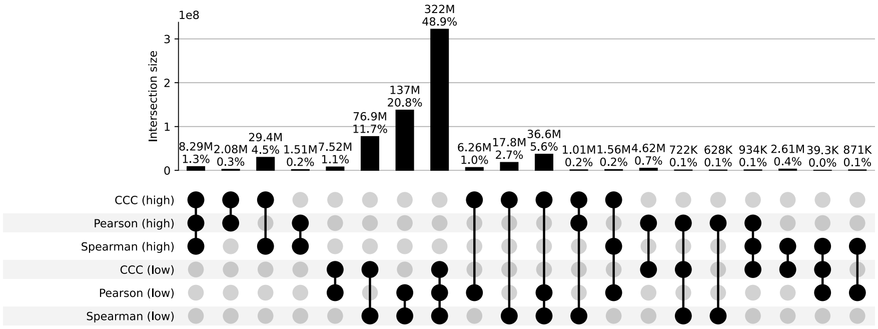


**Figure S32: Distribution and UpSet plots for GTEx v8 fallopian tube.**

#### Heart Atrial Appendage

**a)** Correlation coefficient distributions between gene pairs within GTEx v8 Heart Atrial Appendage

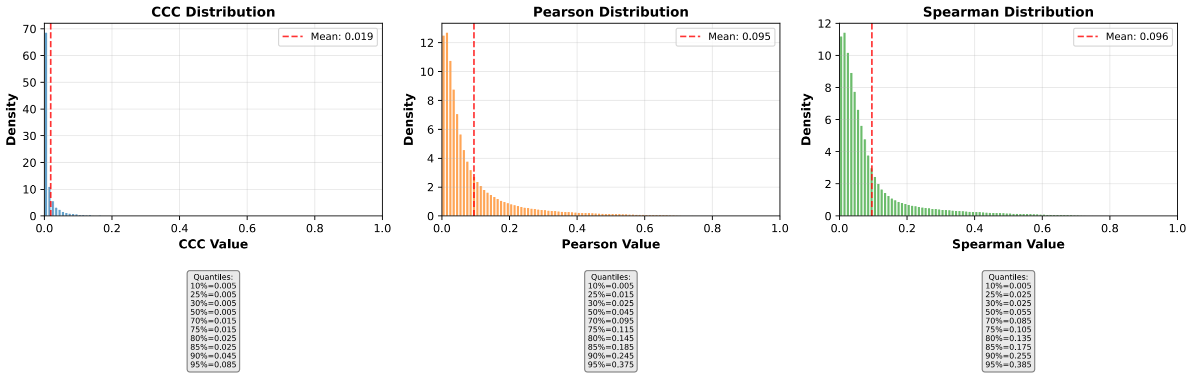


**b)** Corresponding cumulative histogram

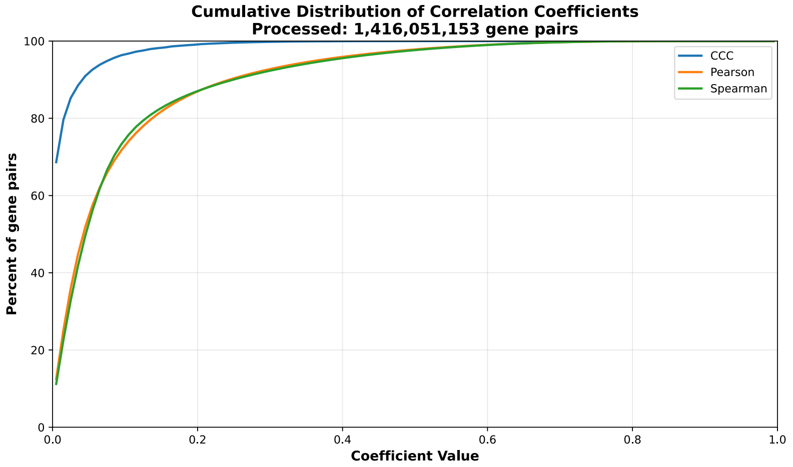


**c)** UpSet plot using top and bottom 30% correlations

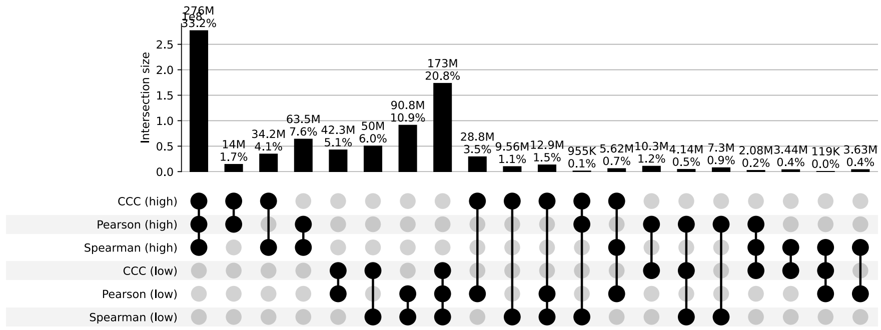


**d)** UpSet plot using permutation-based statistical thresholds

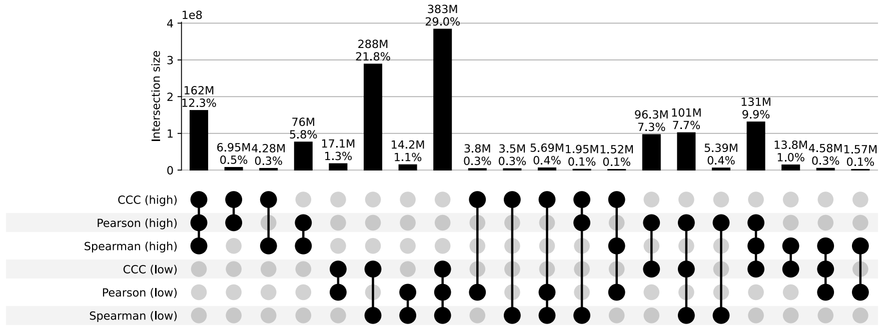


**Figure S33: Distribution and UpSet plots for GTEx v8 heart atrial appendage.**

#### Heart Left Ventricle

**a)** Correlation coefficient distributions between gene pairs within GTEx v8 Heart Left Ventricle

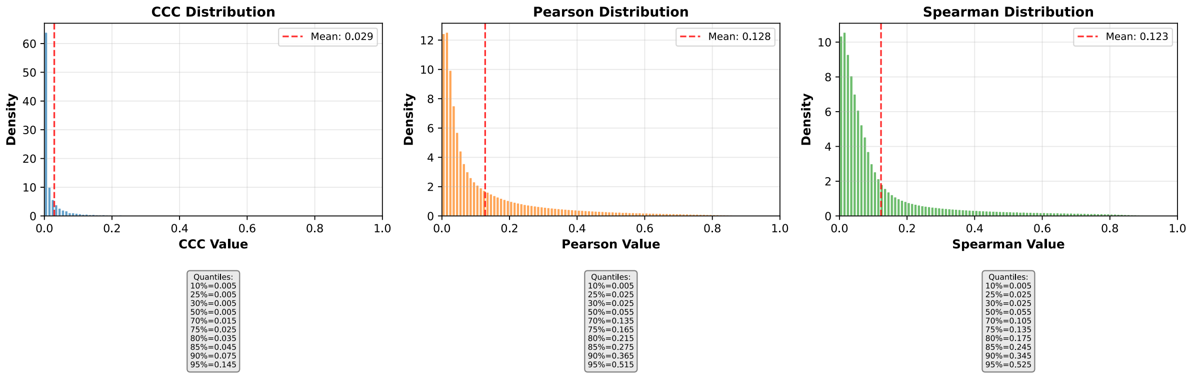


**b)** Corresponding cumulative histogram

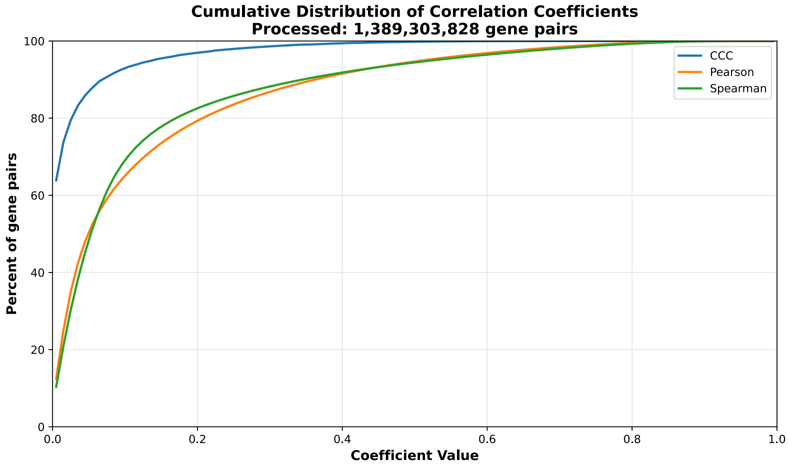


**c)** UpSet plot using top and bottom 30% correlations

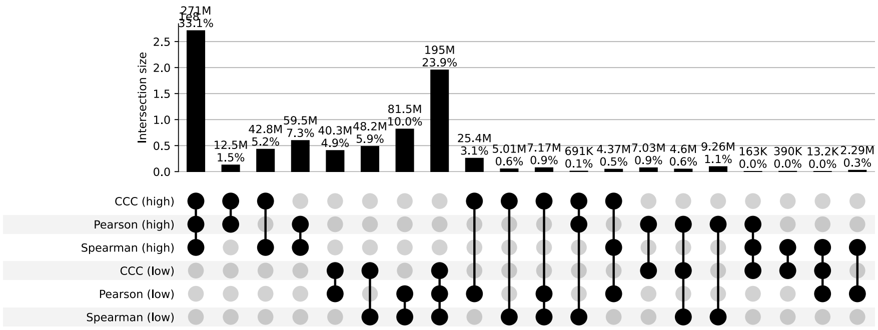


**d)** UpSet plot using permutation-based statistical thresholds

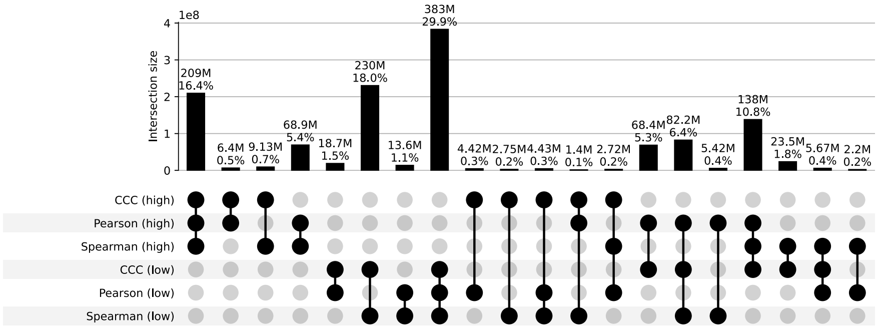


**Figure S34: Distribution and UpSet plots for GTEx v8 heart left ventricle.**

#### Kidney Cortex

**a)** Correlation coefficient distributions between gene pairs within GTEx v8 Kidney Cortex

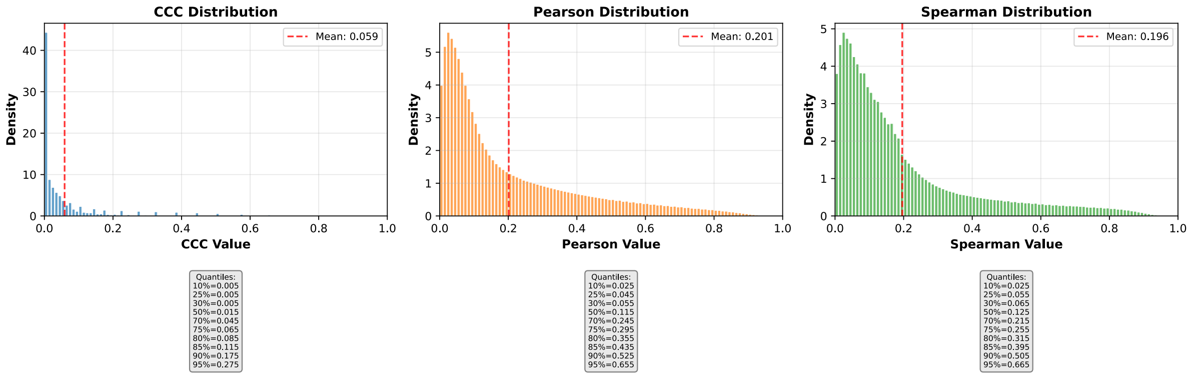


**b)** Corresponding cumulative histogram

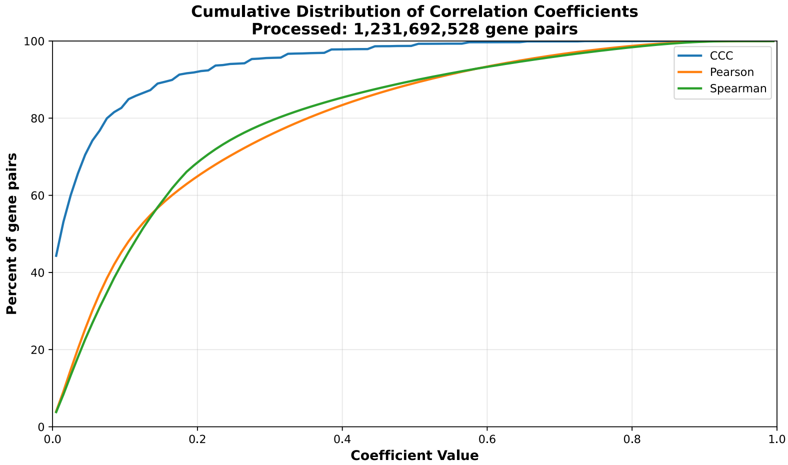


**c)** UpSet plot using top and bottom 30% correlations

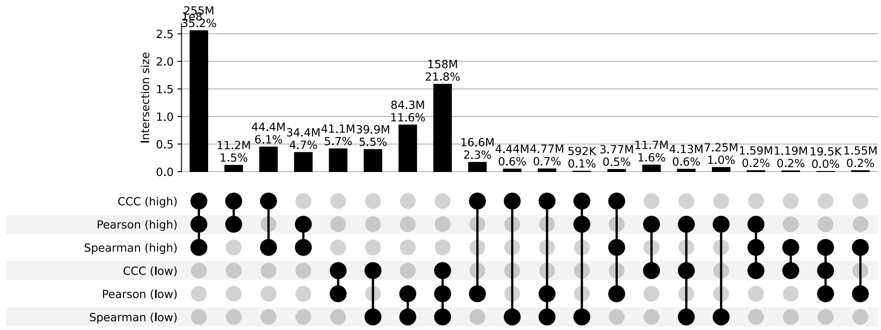


**d)** UpSet plot using permutation-based statistical thresholds

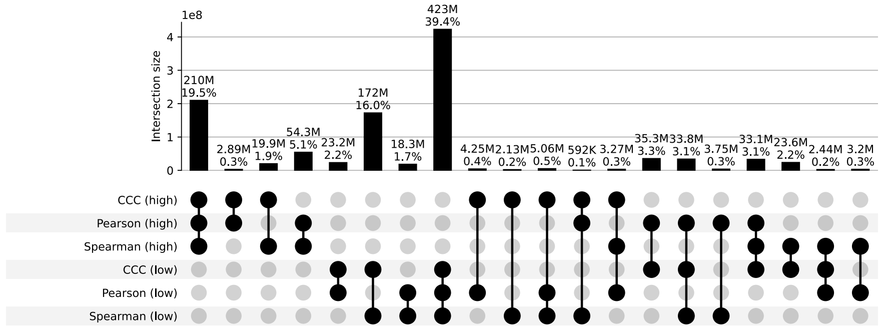


**Figure S35: Distribution and UpSet plots for GTEx v8 kidney cortex.**

#### Kidney Medulla

**a)** Correlation coefficient distributions between gene pairs within GTEx v8 Kidney Medulla

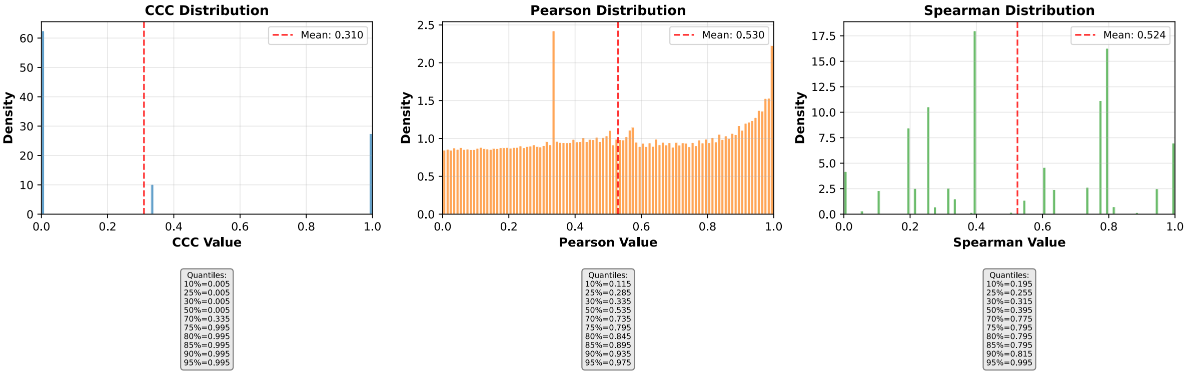


**b)** Corresponding cumulative histogram

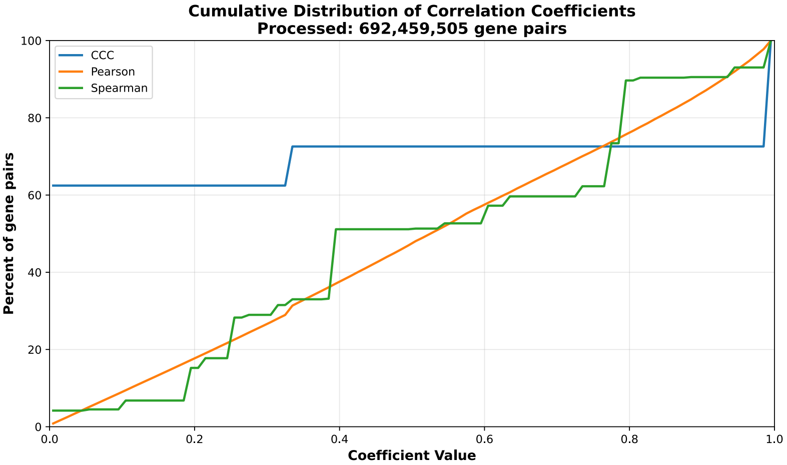


**c)** UpSet plot using top and bottom 30% correlations

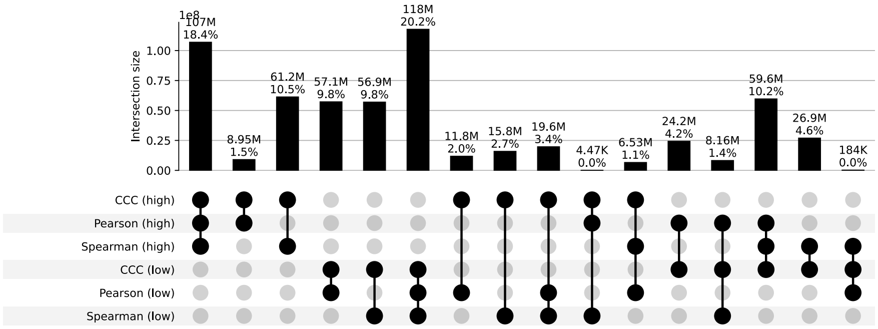


**d)** UpSet plot using permutation-based statistical thresholds

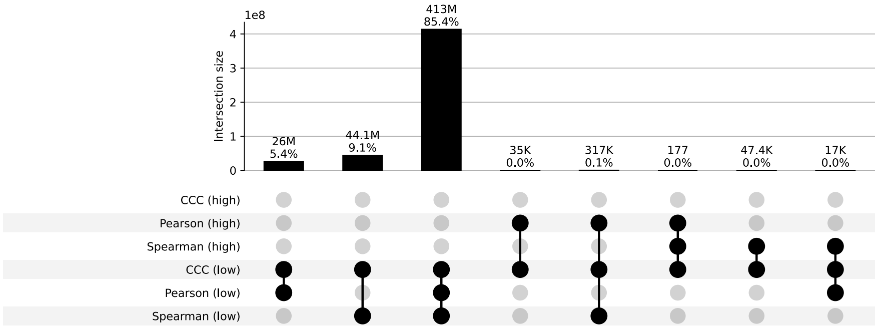


**Figure S36: Distribution and UpSet plots for GTEx v8 kidney medulla.**

#### Liver

**a)** Correlation coefficient distributions between gene pairs within GTEx v8 Liver

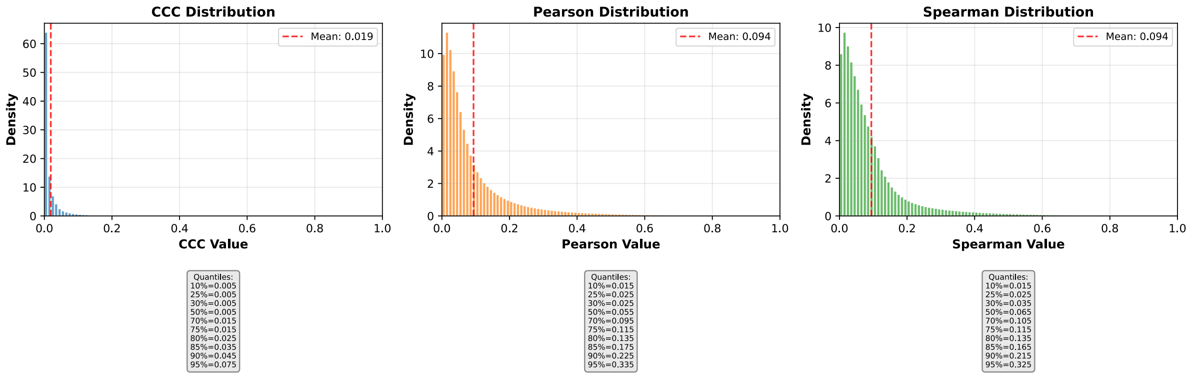


**b)** Corresponding cumulative histogram

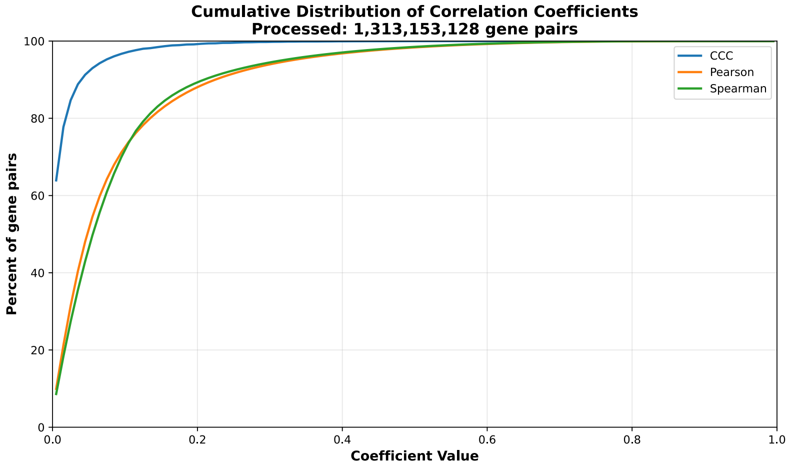


**c)** UpSet plot using top and bottom 30% correlations

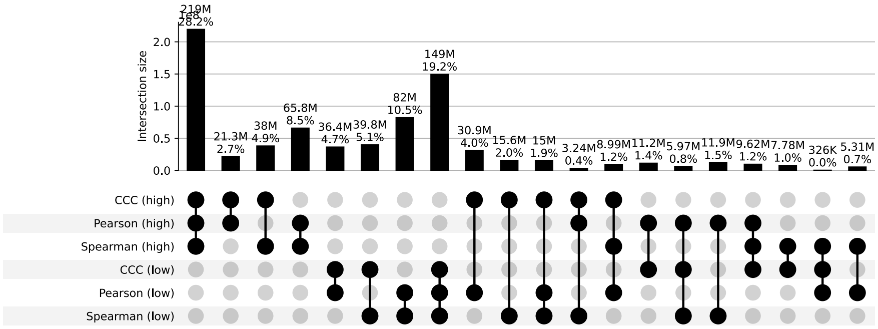


**d)** UpSet plot using permutation-based statistical thresholds

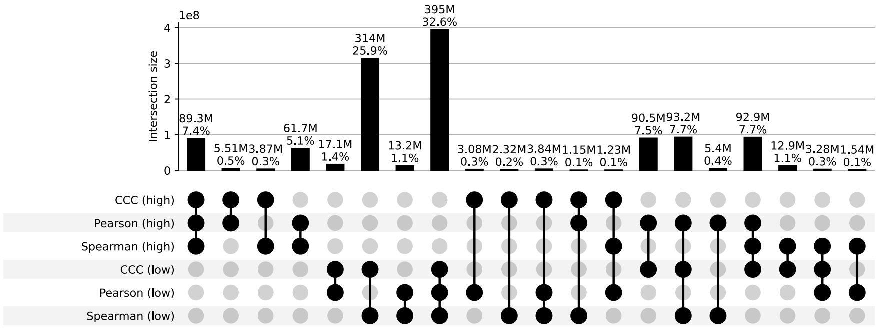


**Figure S37: Distribution and UpSet plots for GTEx v8 liver.**

#### Lung

**a)** Correlation coefficient distributions between gene pairs within GTEx v8 Lung

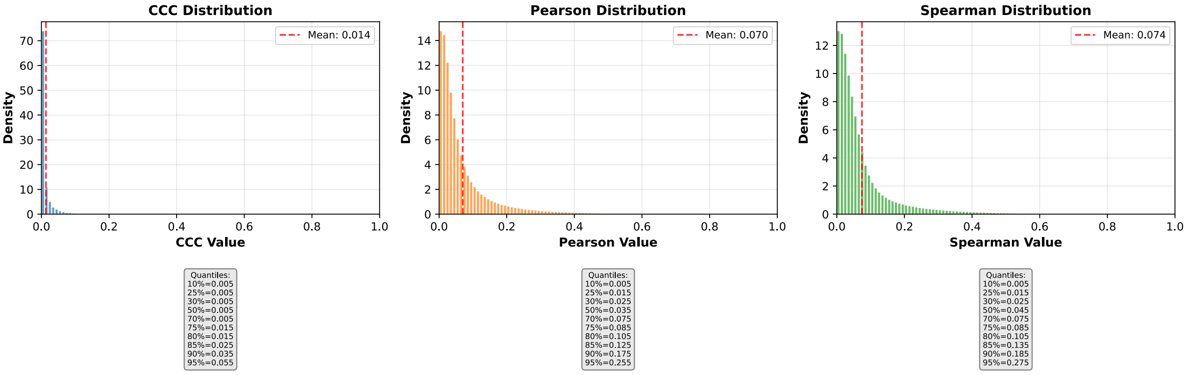


**b)** Corresponding cumulative histogram

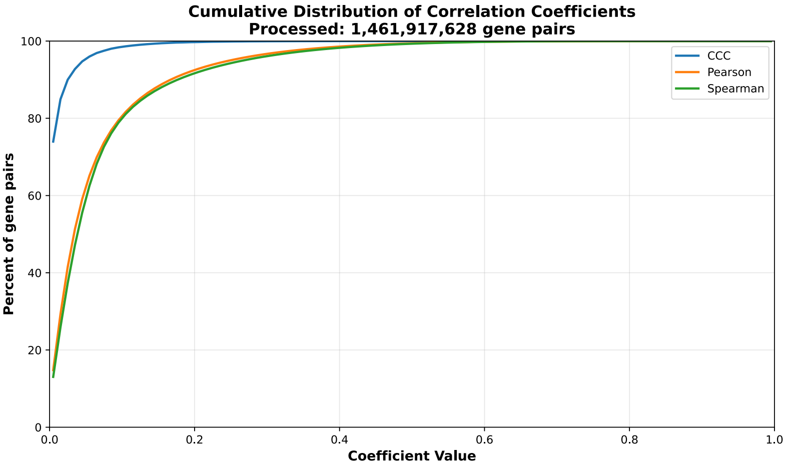


**c)** UpSet plot using top and bottom 30% correlations

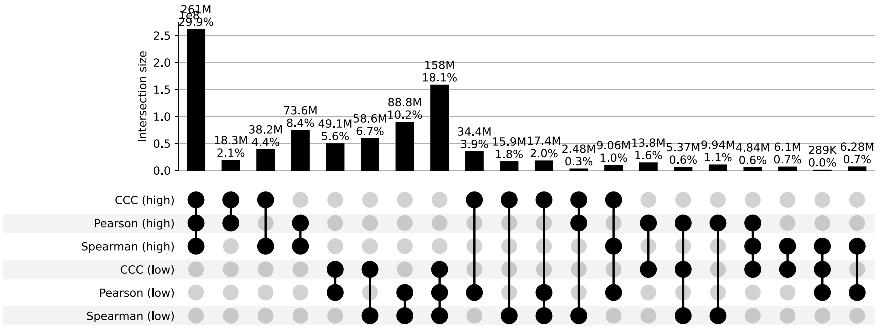


**d)** UpSet plot using permutation-based statistical thresholds

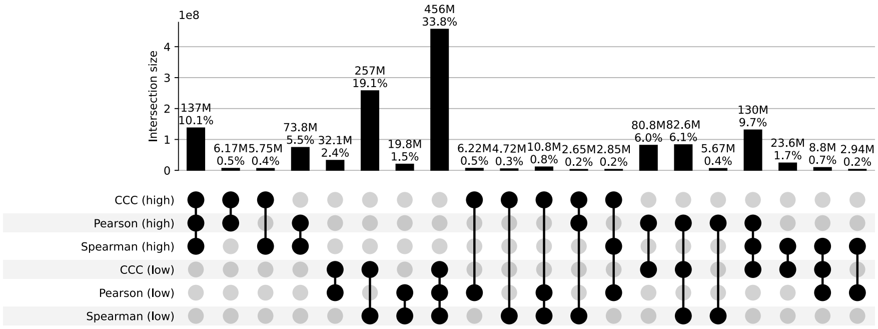


**Figure S38: Distribution and UpSet plots for GTEx v8 lung.**

#### Minor Salivary Gland

**a)** Correlation coefficient distributions between gene pairs within GTEx v8 Minor Salivary Gland

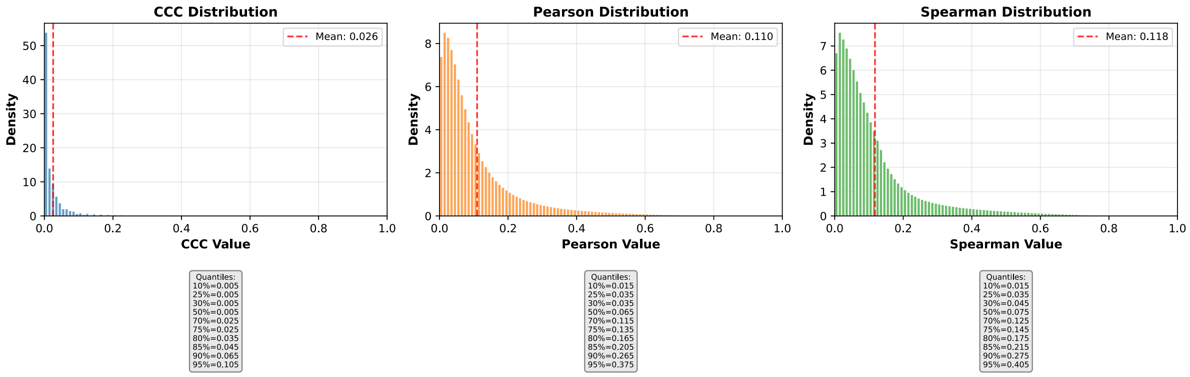


**b)** Corresponding cumulative histogram

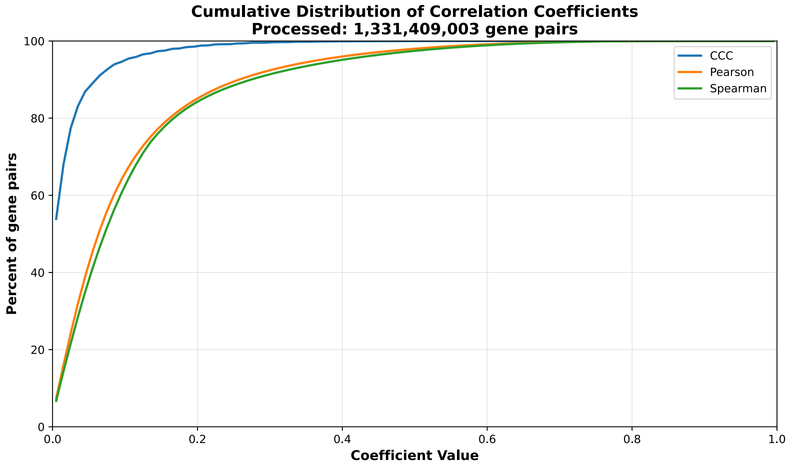


**c)** UpSet plot using top and bottom 30% correlations

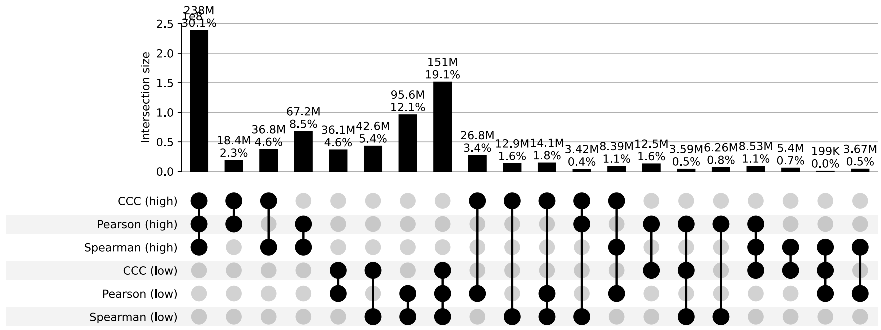


**d)** UpSet plot using permutation-based statistical thresholds

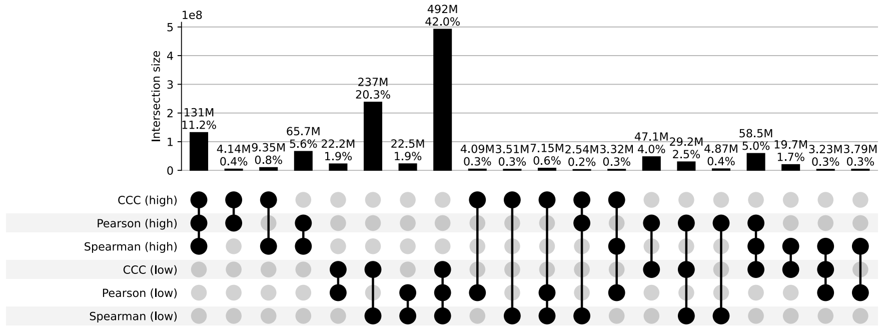


**Figure S39: Distribution and UpSet plots for GTEx v8 minor salivary gland.**

#### Muscle Skeletal

**a)** Correlation coefficient distributions between gene pairs within GTEx v8 Muscle Skeletal

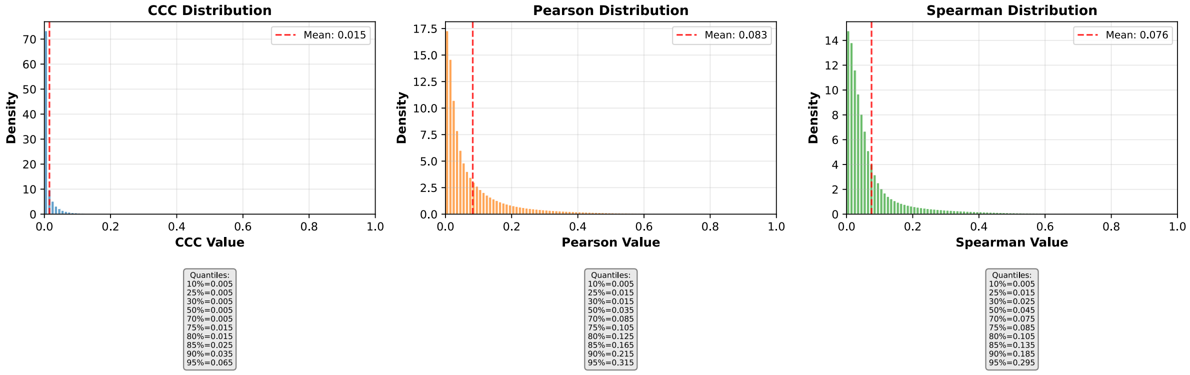


**b)** Corresponding cumulative histogram

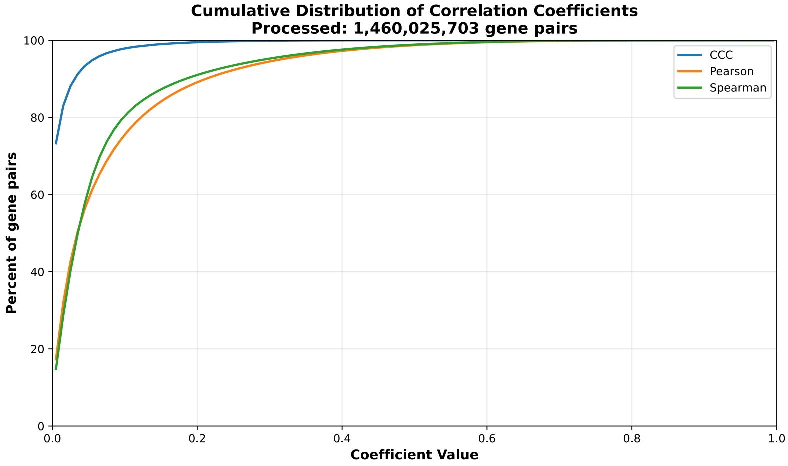


**c)** UpSet plot using top and bottom 30% correlations

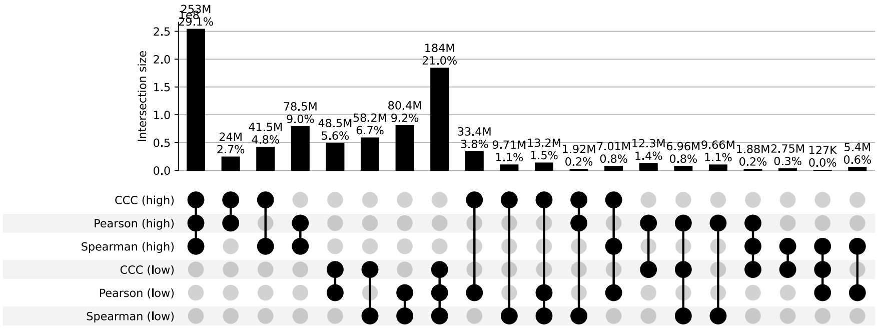


**d)** UpSet plot using permutation-based statistical thresholds

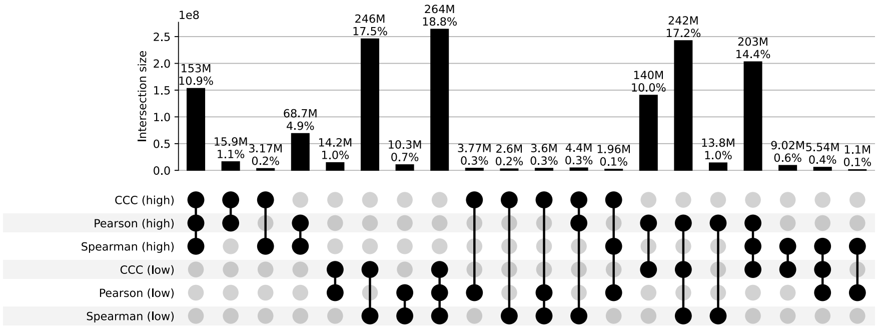


**Figure S40: Distribution and UpSet plots for GTEx v8 muscle skeletal.**

#### Nerve Tibial

**a)** Correlation coefficient distributions between gene pairs within GTEx v8 Nerve Tibial

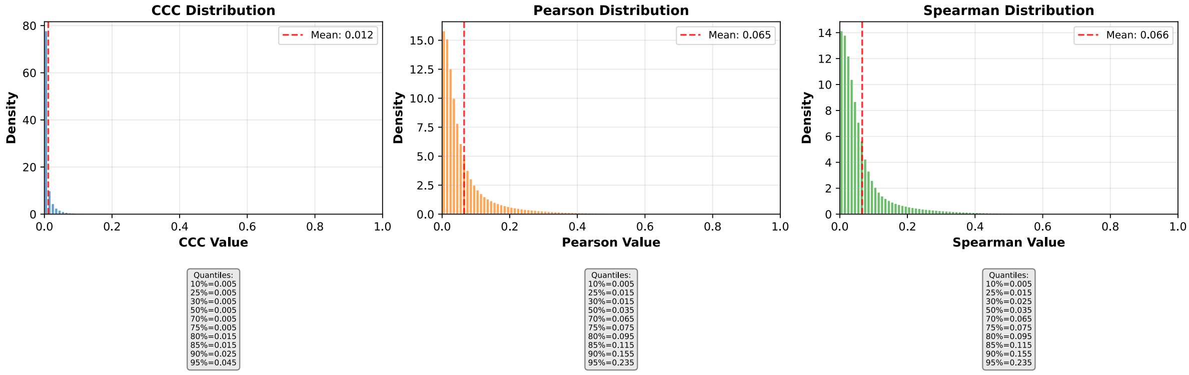


**b)** Corresponding cumulative histogram

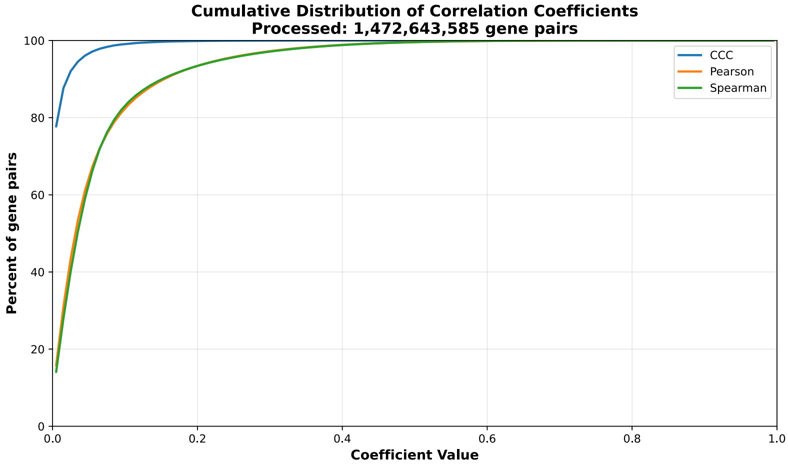


**c)** UpSet plot using top and bottom 30% correlations

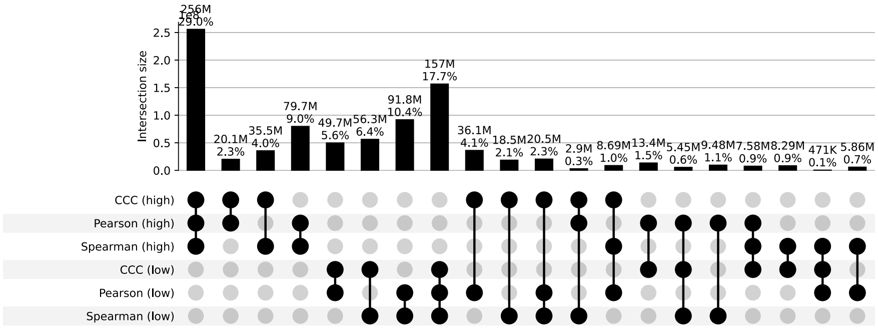


**d)** UpSet plot using permutation-based statistical thresholds

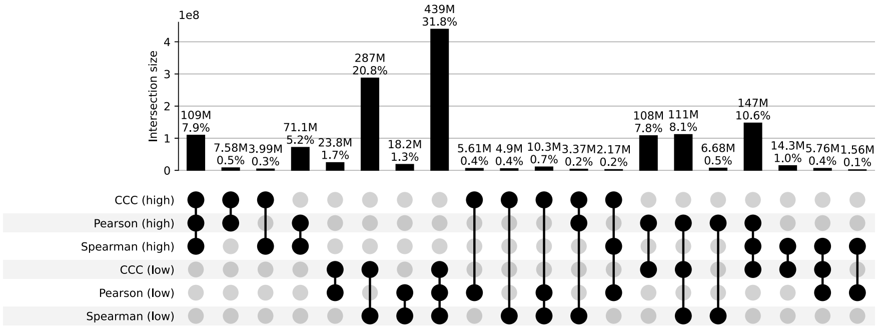


**Figure S41: Distribution and UpSet plots for GTEx v8 nerve tibial.**

#### Ovary

**a)** Correlation coefficient distributions between gene pairs within GTEx v8 Ovary

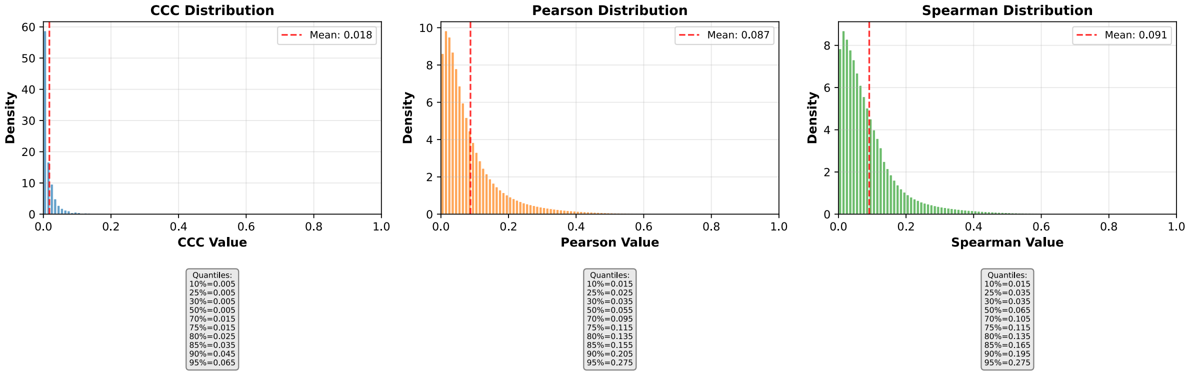


**b)** Corresponding cumulative histogram

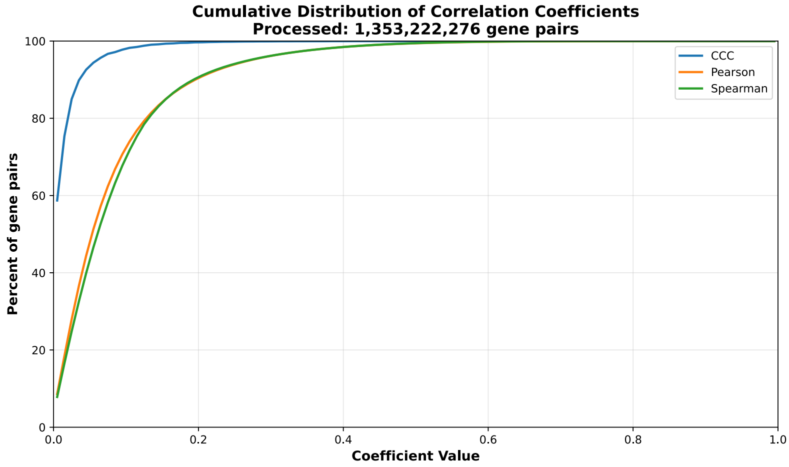


**c)** UpSet plot using top and bottom 30% correlations

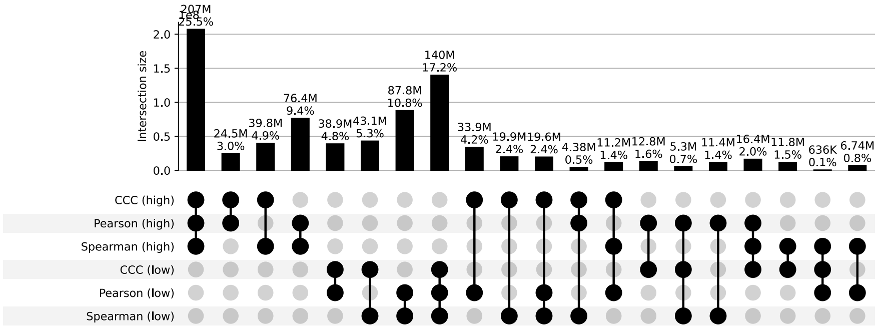


**d)** UpSet plot using permutation-based statistical thresholds

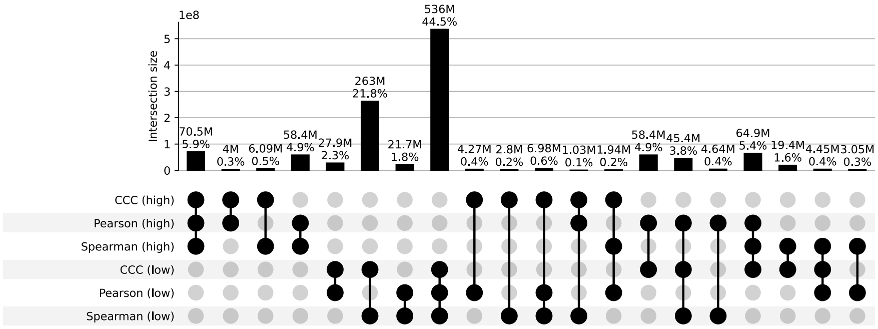


**Figure S42: Distribution and UpSet plots for GTEx v8 ovary.**

#### Pancreas

**a)** Correlation coefficient distributions between gene pairs within GTEx v8 Pancreas

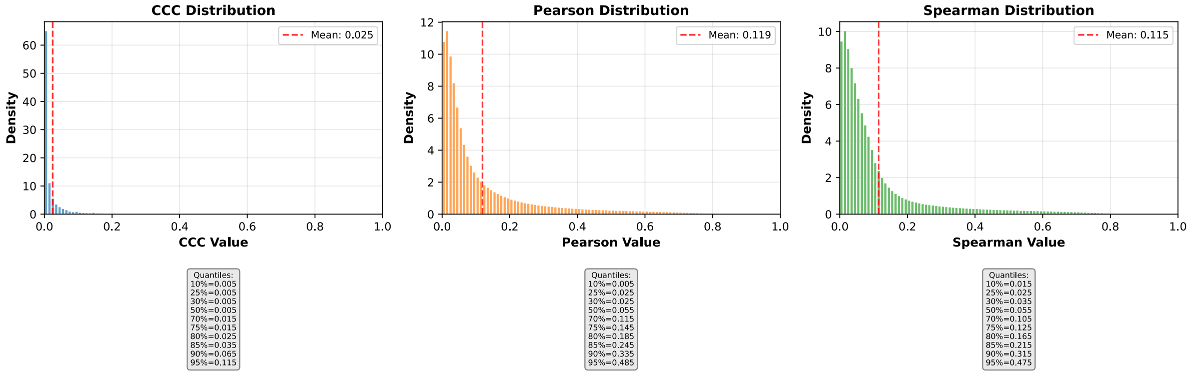


**b)** Corresponding cumulative histogram

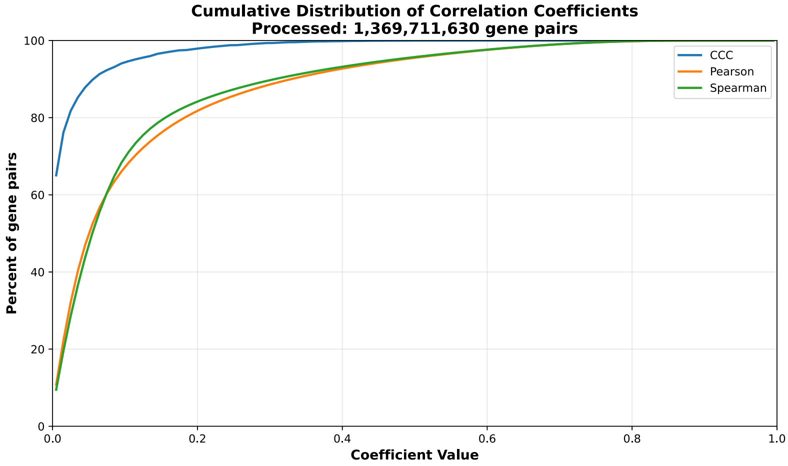


**c)** UpSet plot using top and bottom 30% correlations

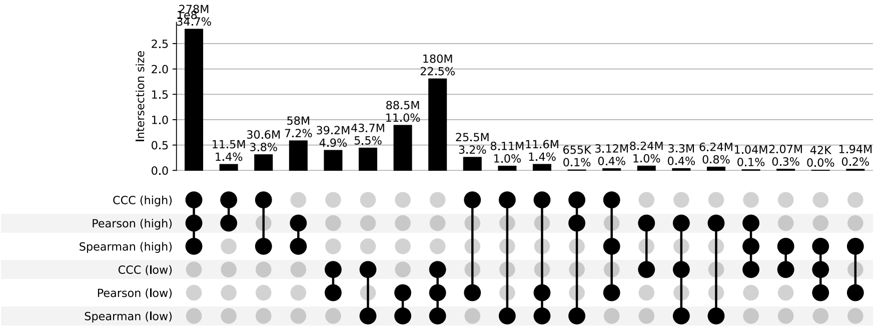


**d)** UpSet plot using permutation-based statistical thresholds

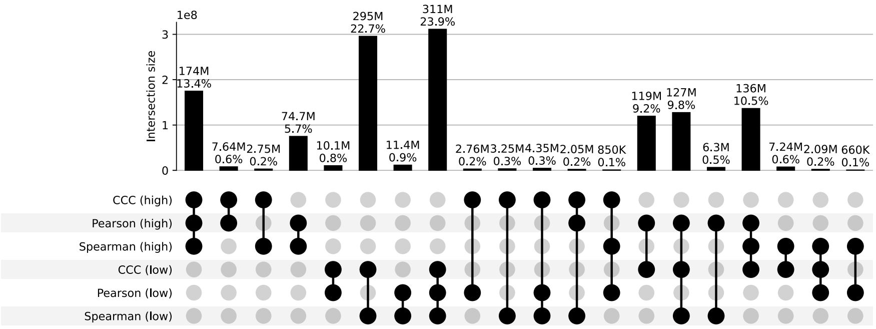


**Figure S43: Distribution and UpSet plots for GTEx v8 pancreas.**

#### Pituitary

**a)** Correlation coefficient distributions between gene pairs within GTEx v8 Pituitary

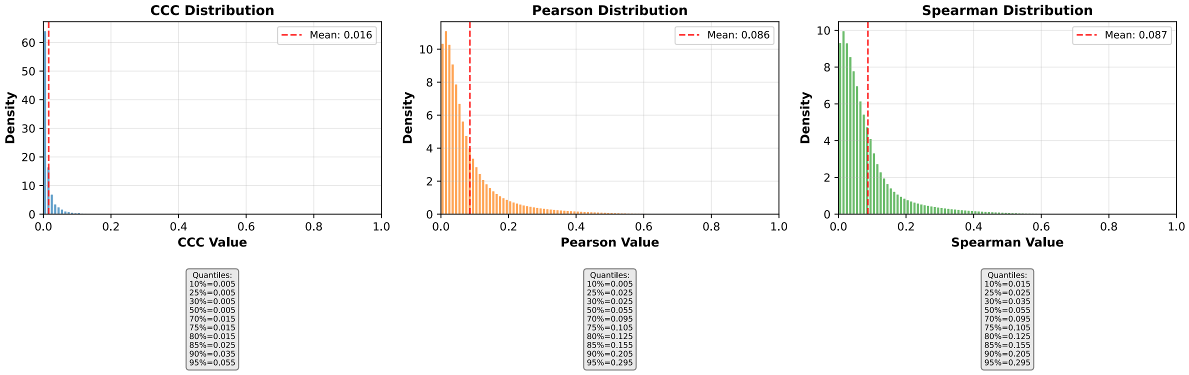


**b)** Corresponding cumulative histogram

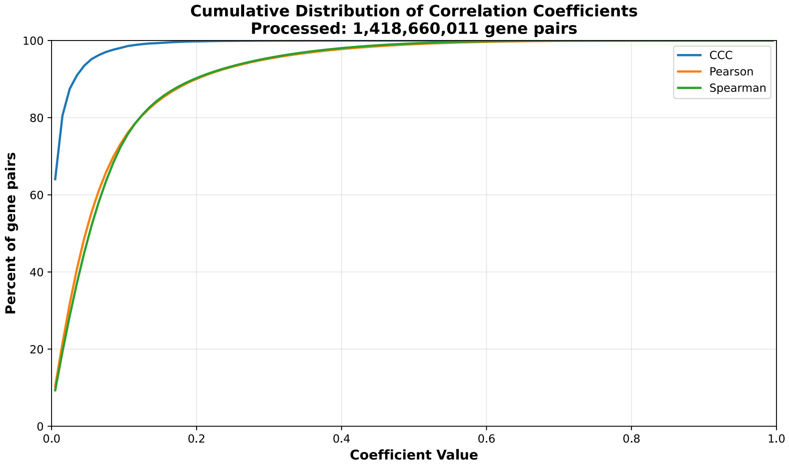


**c)** UpSet plot using top and bottom 30% correlations

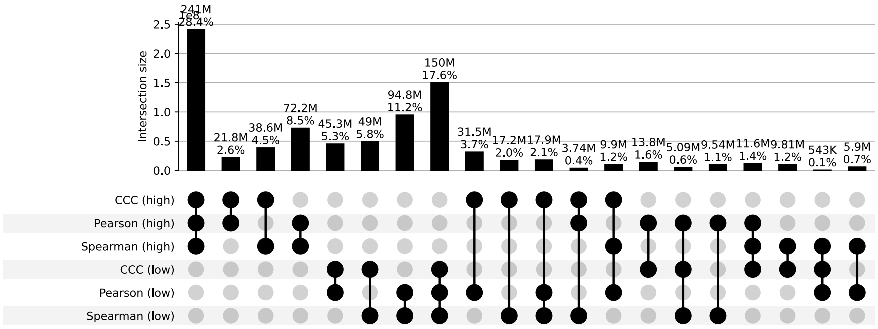


**d)** UpSet plot using permutation-based statistical thresholds

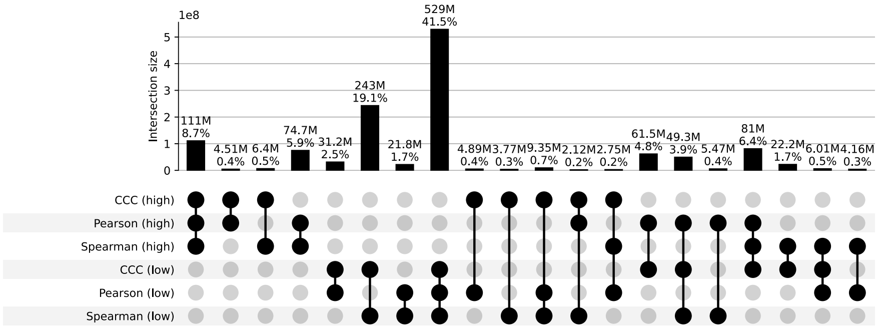


**Figure S44: Distribution and UpSet plots for GTEx v8 pituitary.**

#### Prostate

**a)** Correlation coefficient distributions between gene pairs within GTEx v8 Prostate

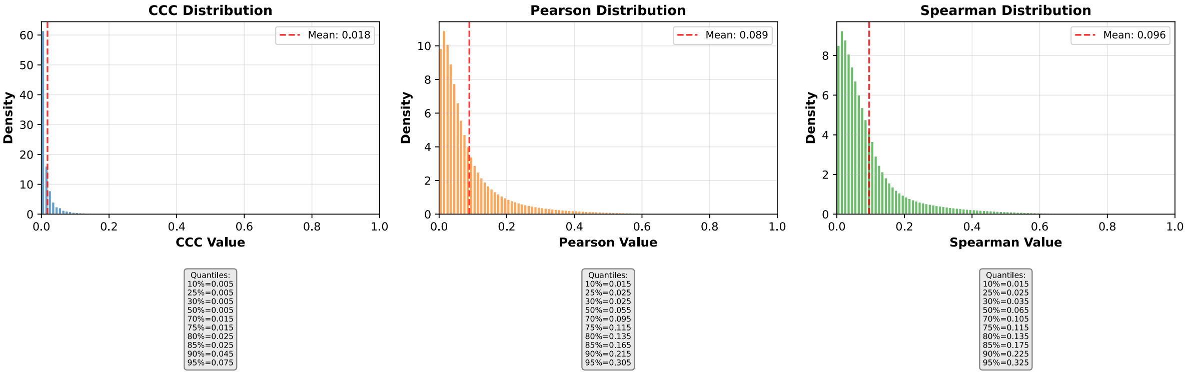


**b)** Corresponding cumulative histogram

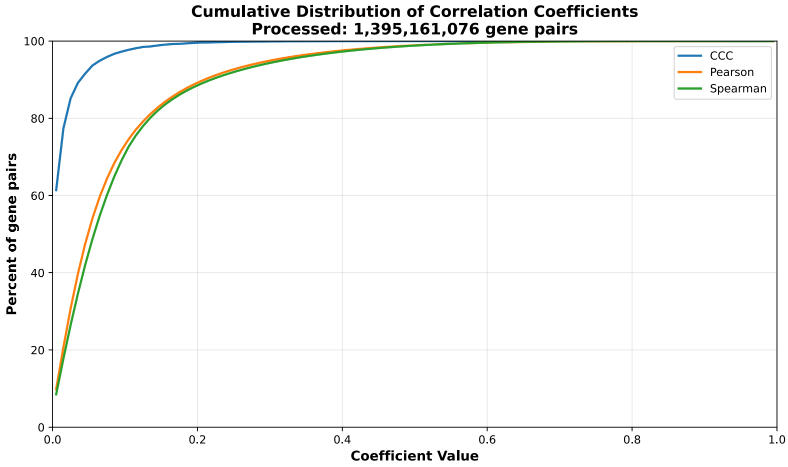


**c)** UpSet plot using top and bottom 30% correlations

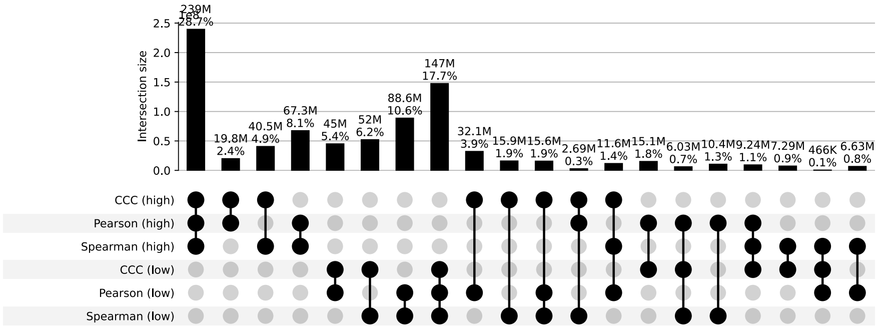


**d)** UpSet plot using permutation-based statistical thresholds

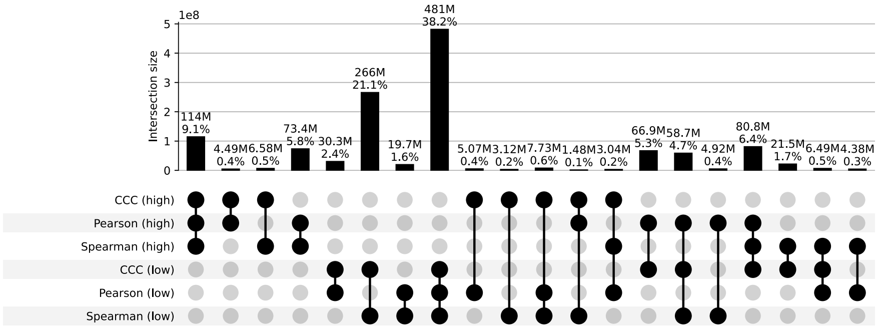


**Figure S45: Distribution and UpSet plots for GTEx v8 prostate.**

### Skin Not Sun Exposed Suprapubic

**a)** Correlation coefficient distributions between gene pairs within GTEx v8 Skin Not Sun Exposed Suprapubic

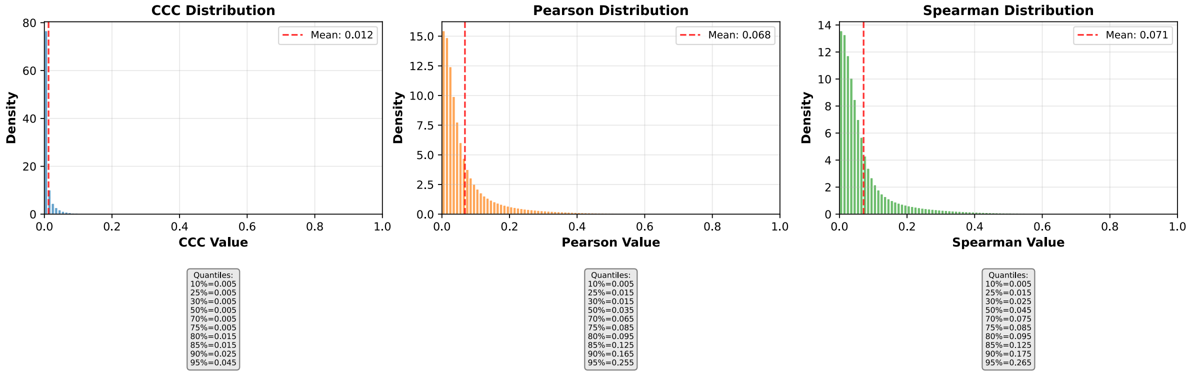


**b)** Corresponding cumulative histogram

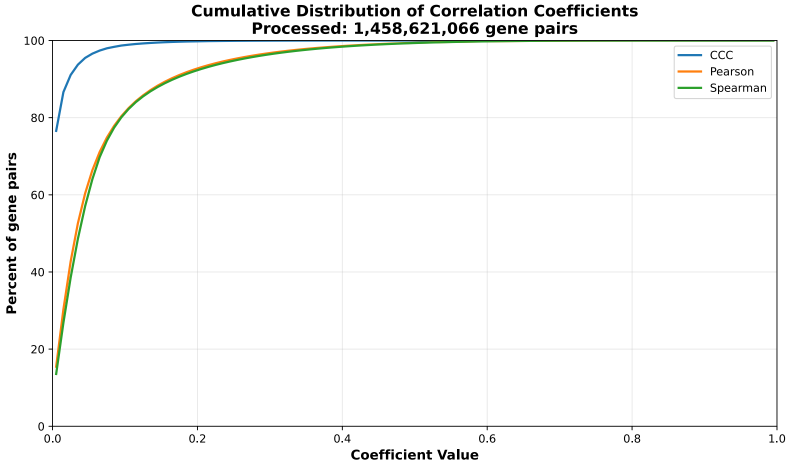


**c)** UpSet plot using top and bottom 30% correlations

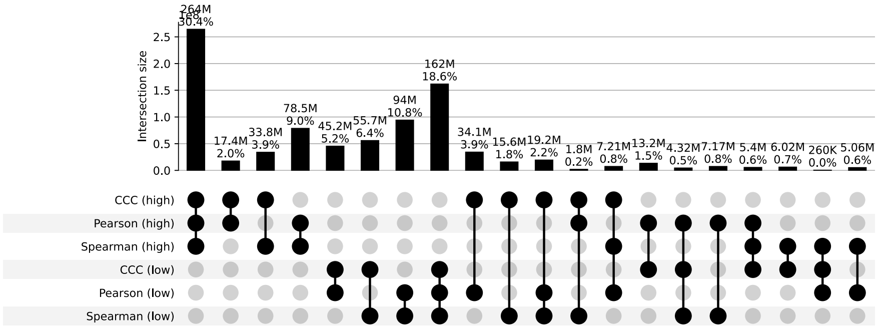


**d)** UpSet plot using permutation-based statistical thresholds

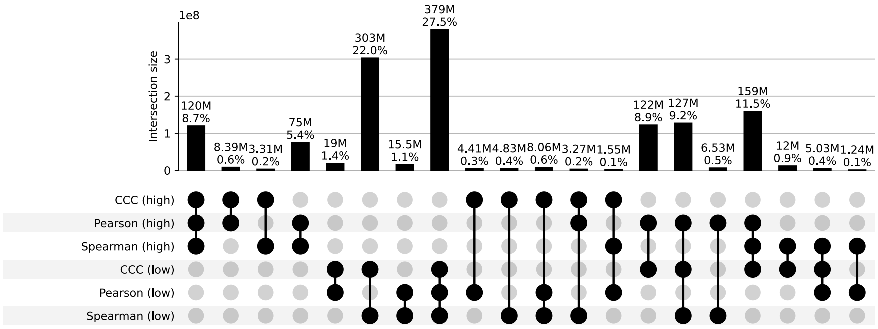


**Figure S46: Distribution and UpSet plots for GTEx v8 skin not sun exposed suprapubic.**

#### Skin Sun Exposed Lower Leg

**a)** Correlation coefficient distributions between gene pairs within GTEx v8 Skin Sun Exposed Lower Leg

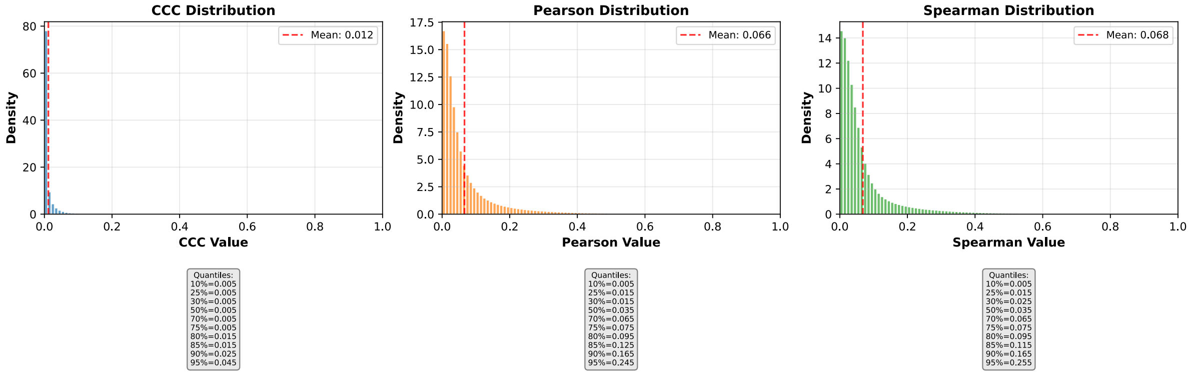


**b)** Corresponding cumulative histogram

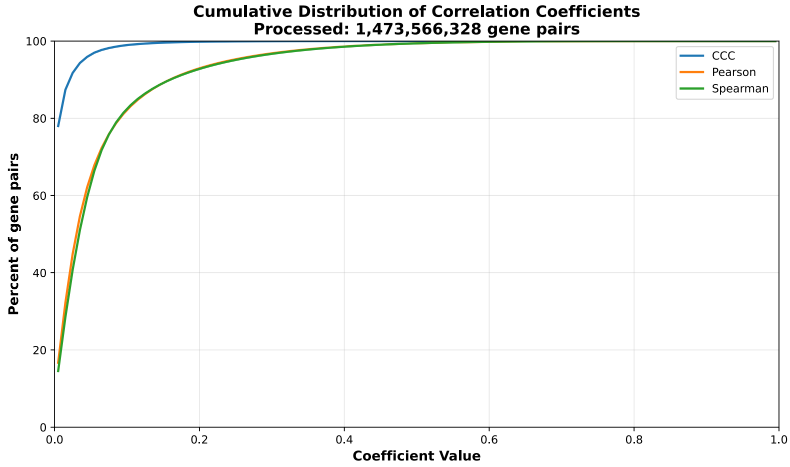


**c)** UpSet plot using top and bottom 30% correlations

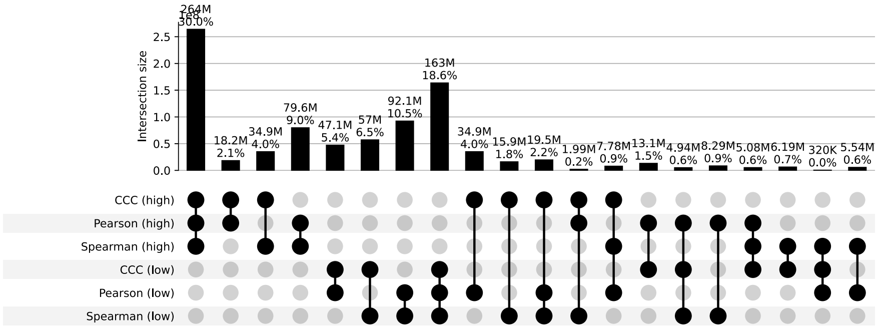


**d)** UpSet plot using permutation-based statistical thresholds

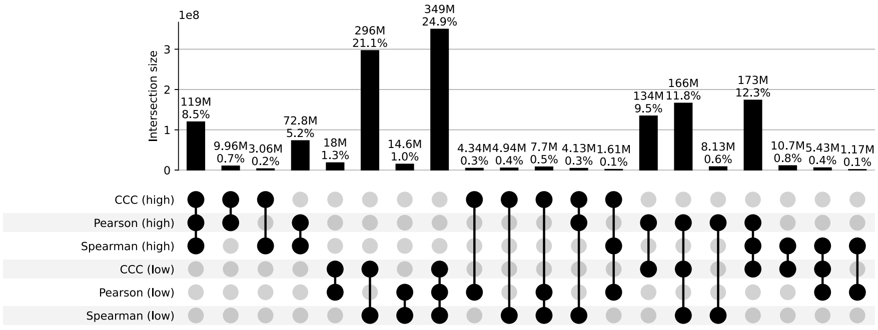


**Figure S47: Distribution and UpSet plots for GTEx v8 skin sun exposed lower leg.**

#### Small Intestine Terminal Ileum

**a)** Correlation coefficient distributions between gene pairs within GTEx v8 Small Intestine Terminal Ileum

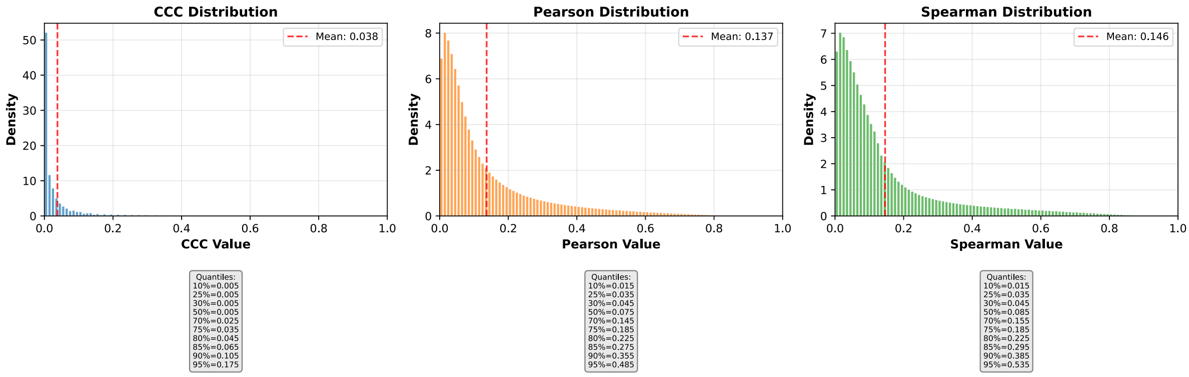


**b)** Corresponding cumulative histogram

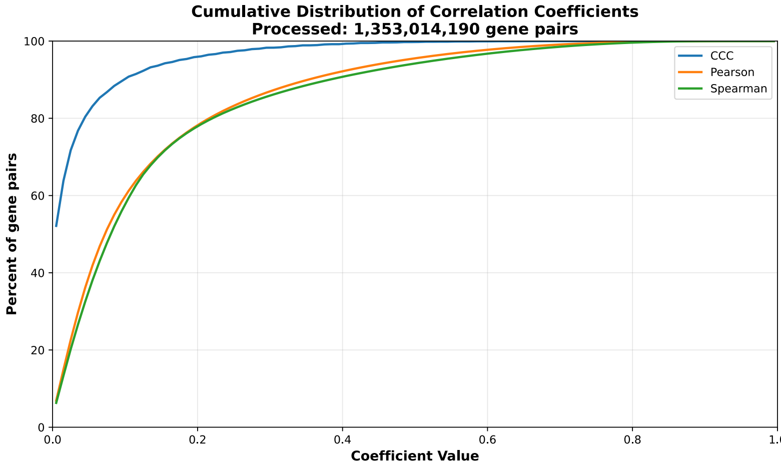


**c)** UpSet plot using top and bottom 30% correlations

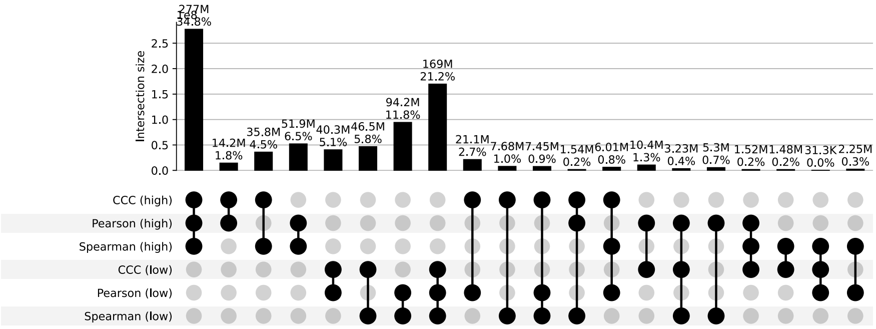


**d)** UpSet plot using permutation-based statistical thresholds

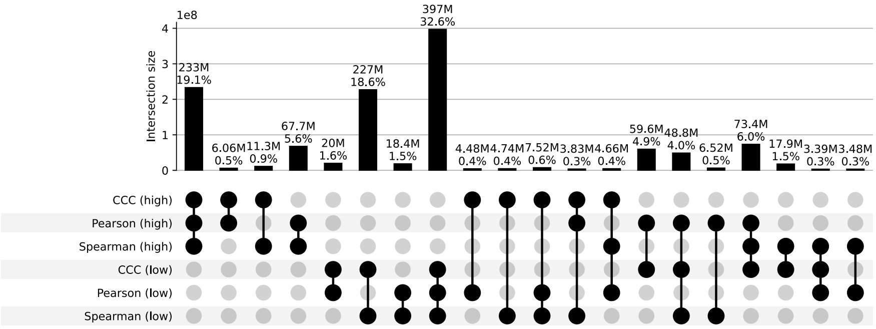


**Figure S48: Distribution and UpSet plots for GTEx v8 small intestine terminal ileum.**

#### Spleen

**a)** Correlation coefficient distributions between gene pairs within GTEx v8 Spleen

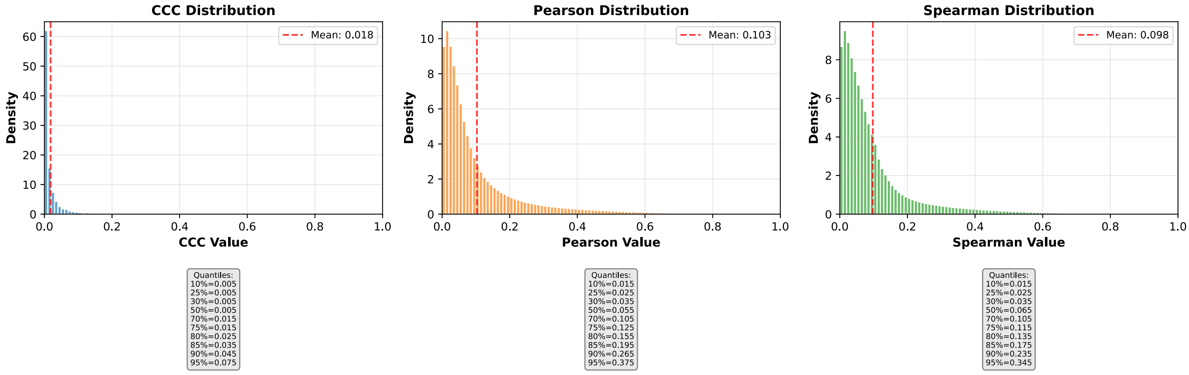


**b)** Corresponding cumulative histogram

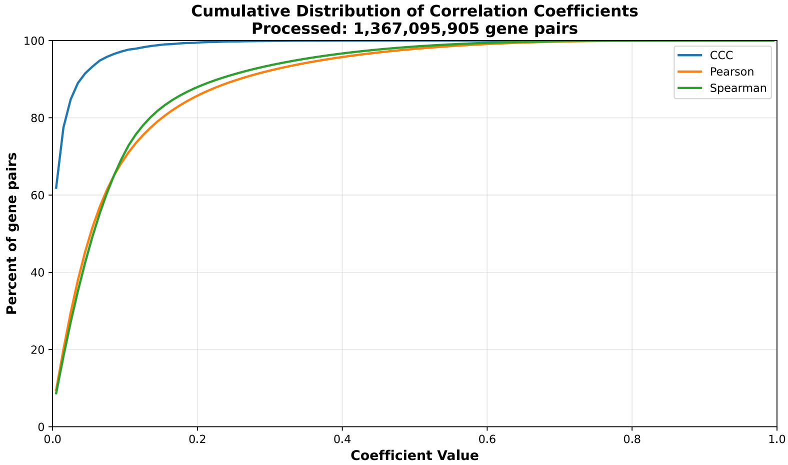


**c)** UpSet plot using top and bottom 30% correlations

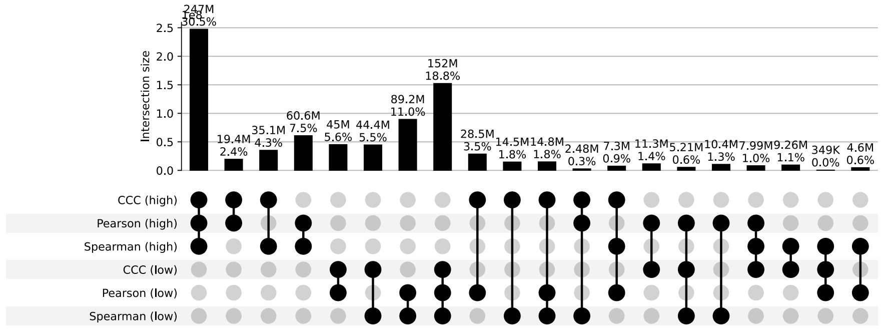


**d)** UpSet plot using permutation-based statistical thresholds

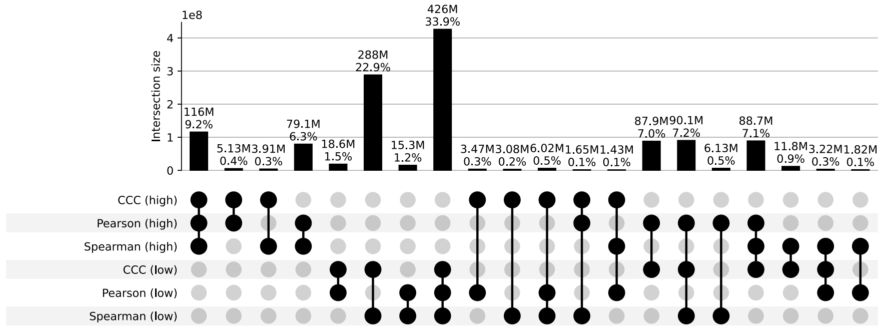


**Figure S49: Distribution and UpSet plots for GTEx v8 spleen.**

#### Stomach

**a)** Correlation coefficient distributions between gene pairs within GTEx v8 Stomach

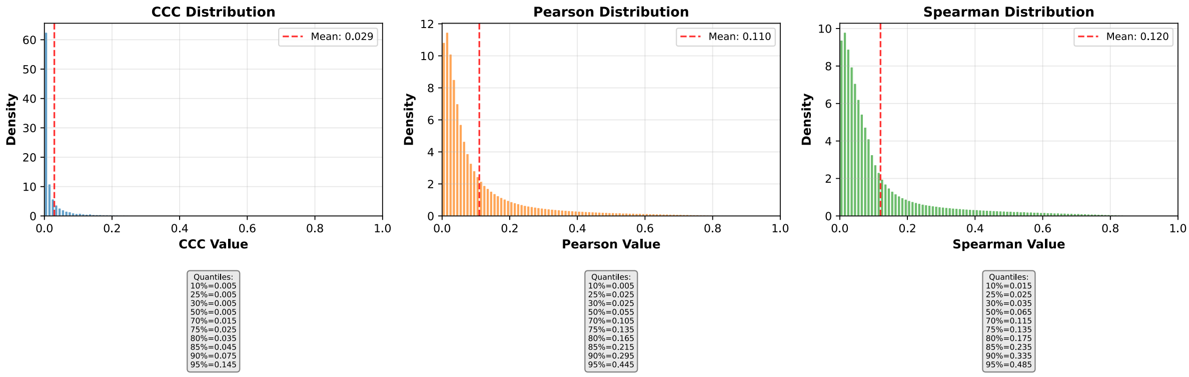


**b)** Corresponding cumulative histogram

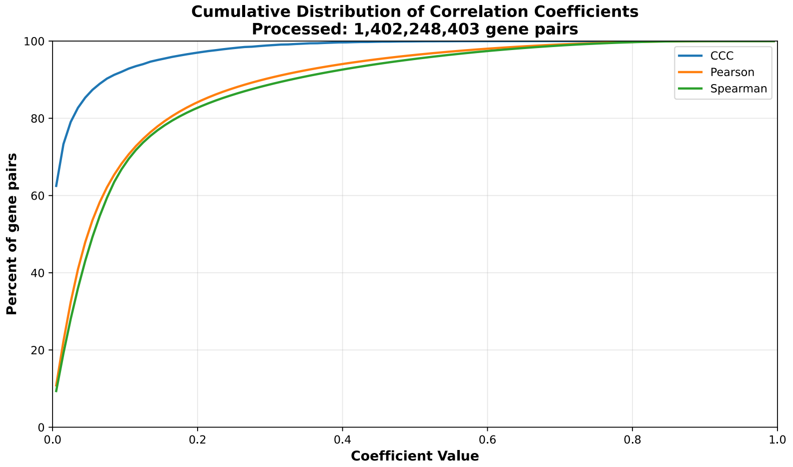


**c)** UpSet plot using top and bottom 30% correlations

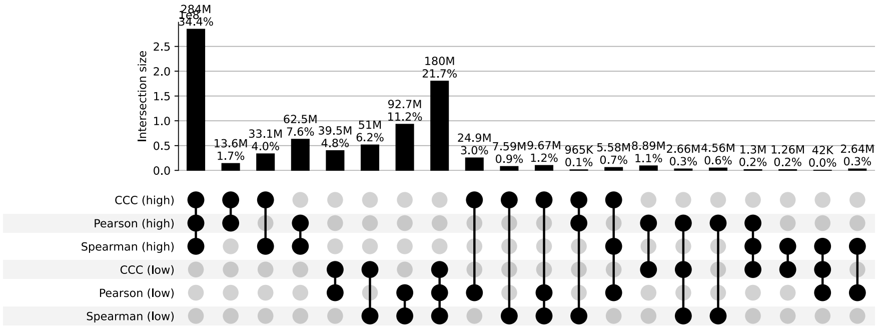


**d)** UpSet plot using permutation-based statistical thresholds

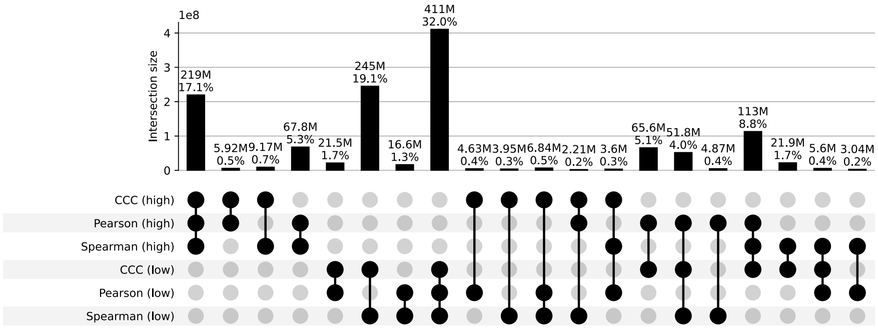


**Figure S50: Distribution and UpSet plots for GTEx v8 stomach.**

#### Testis

**a)** Correlation coefficient distributions between gene pairs within GTEx v8 Testis

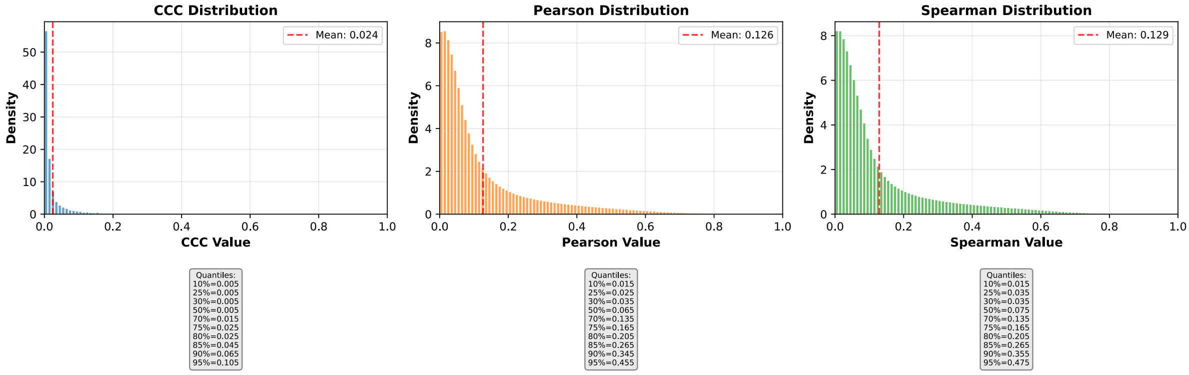


**b)** Corresponding cumulative histogram

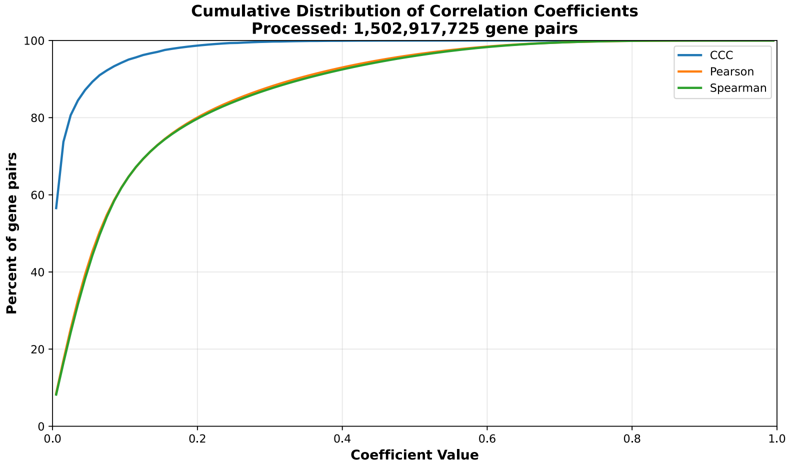


**c)** UpSet plot using top and bottom 30% correlations

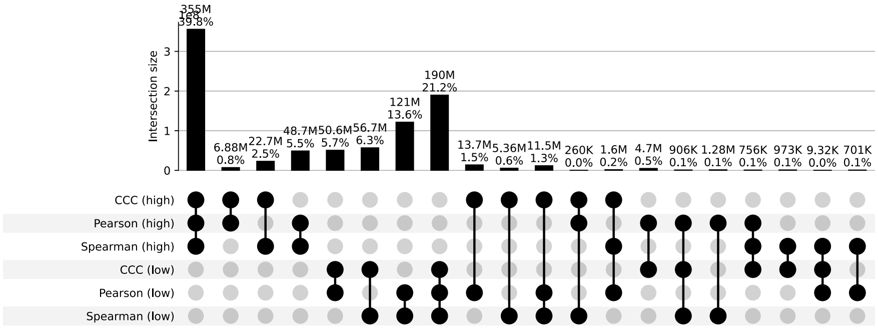


**d)** UpSet plot using permutation-based statistical thresholds

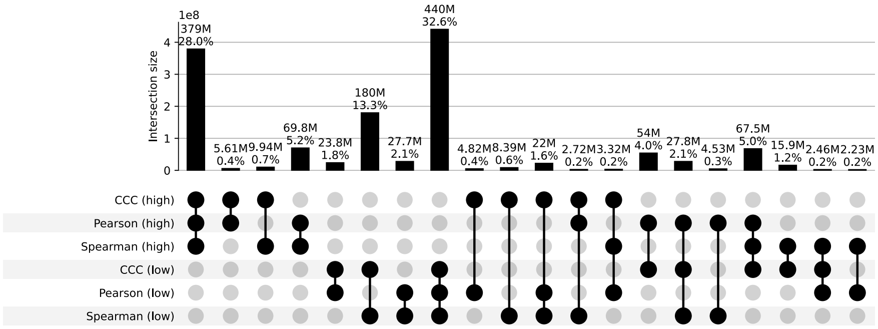


**Figure S51: Distribution and UpSet plots for GTEx v8 testis.**

#### Thyroid

**a)** Correlation coefficient distributions between gene pairs within GTEx v8 Thyroid

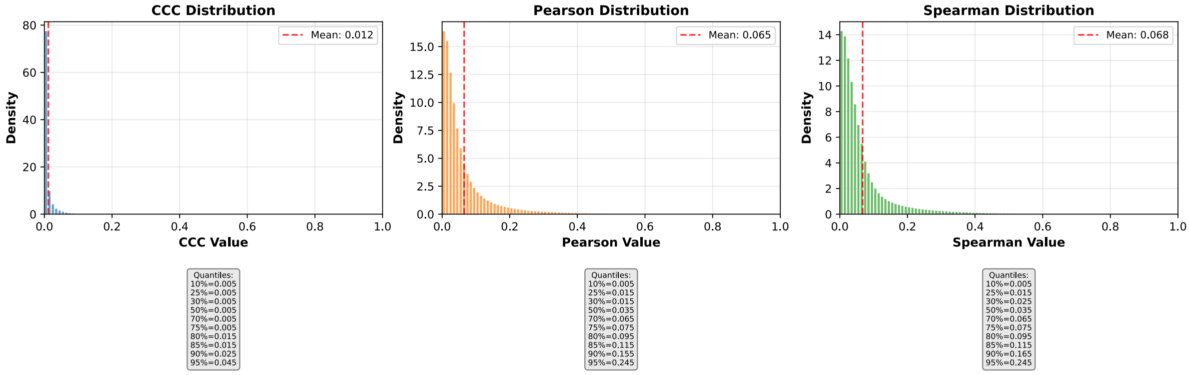


**b)** Corresponding cumulative histogram

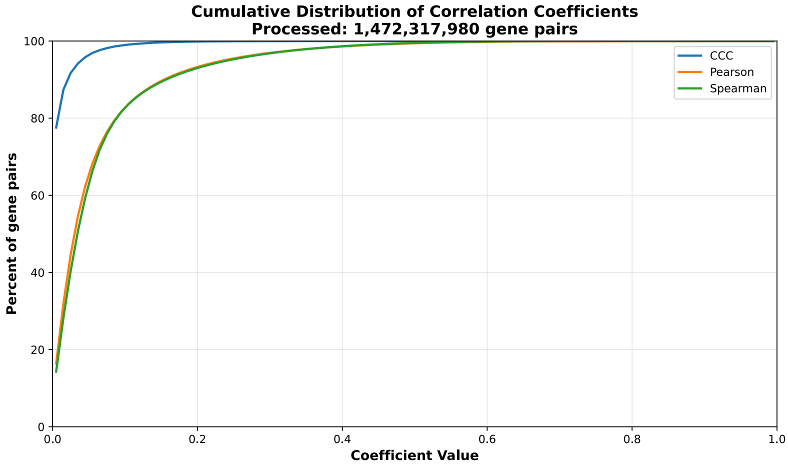


**c)** UpSet plot using top and bottom 30% correlations

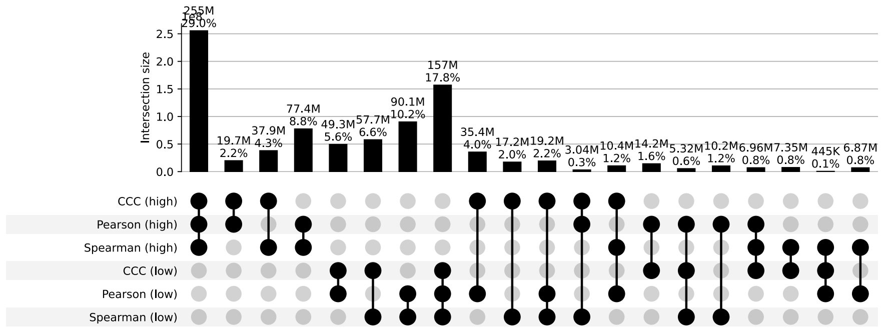


**d)** UpSet plot using permutation-based statistical thresholds

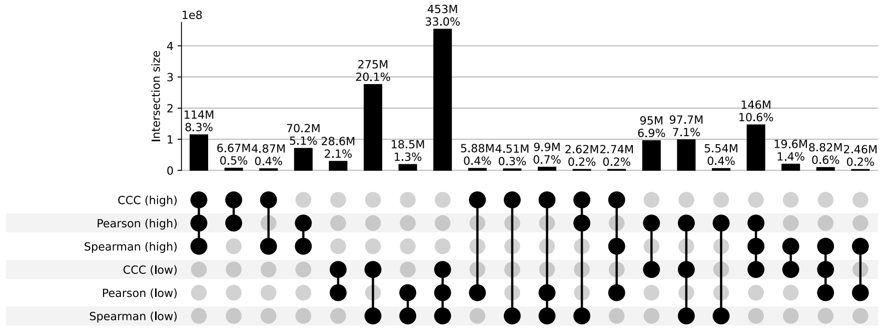


**Figure S52: Distribution and UpSet plots for GTEx v8 thyroid.**

#### Uterus

**a)** Correlation coefficient distributions between gene pairs within GTEx v8 Uterus

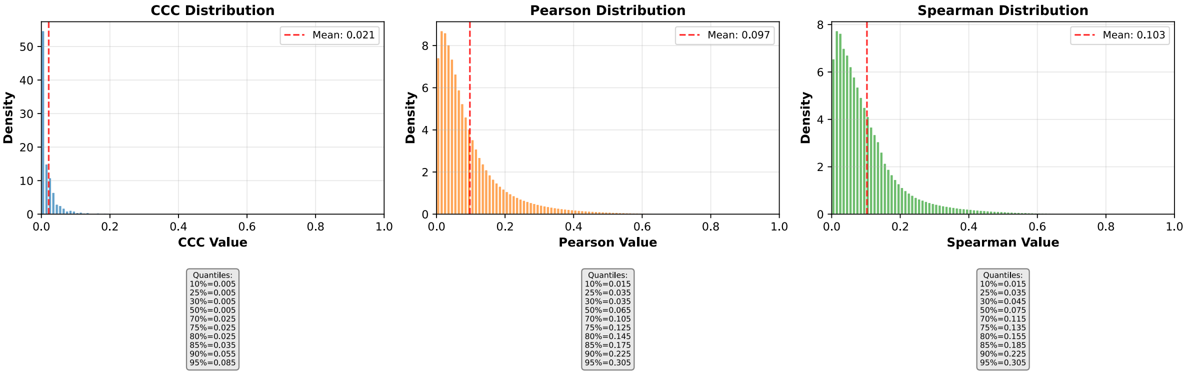


**b)** Corresponding cumulative histogram

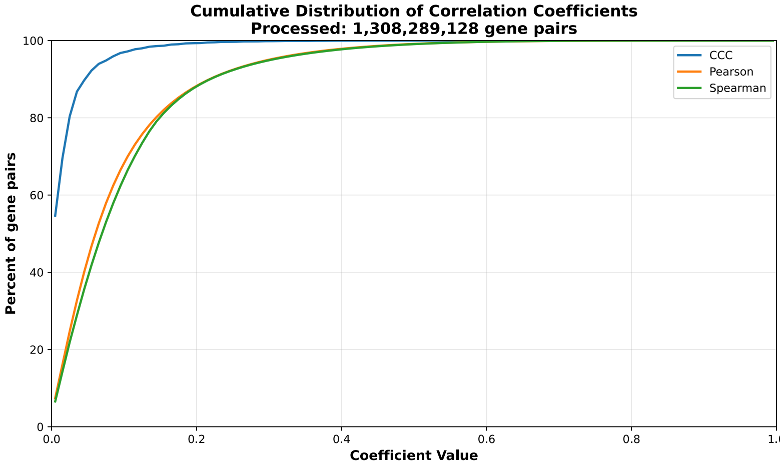


**c)** UpSet plot using top and bottom 30% correlations

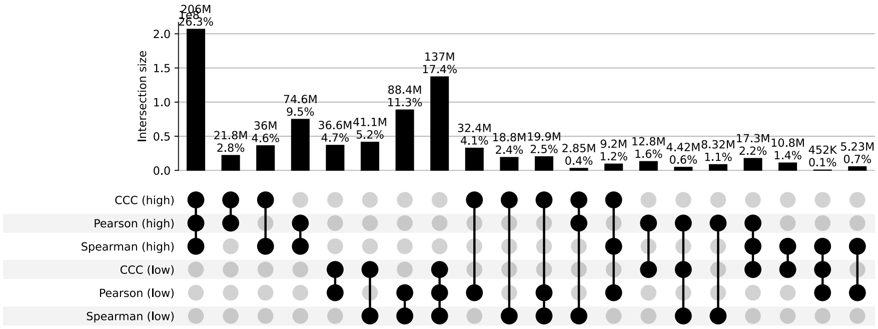


**d)** UpSet plot using permutation-based statistical thresholds

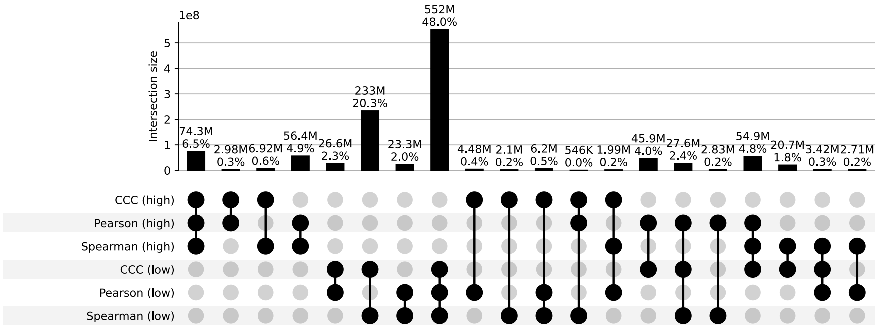


**Figure S53: Distribution and UpSet plots for GTEx v8 uterus.**

#### Vagina

**a)** Correlation coefficient distributions between gene pairs within GTEx v8 Vagina

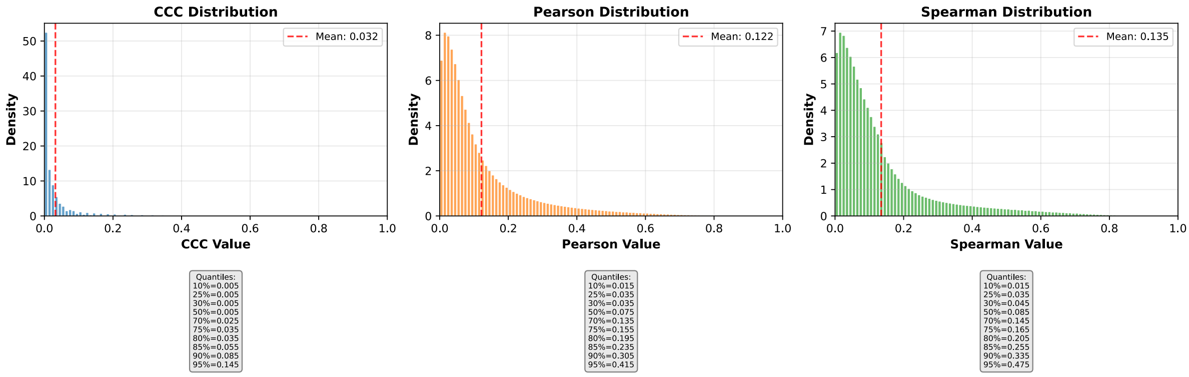


**b)** Corresponding cumulative histogram

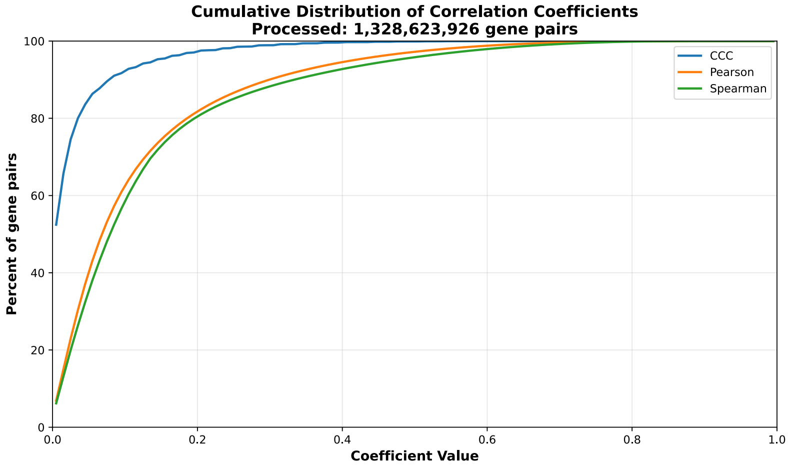


**c)** UpSet plot using top and bottom 30% correlations

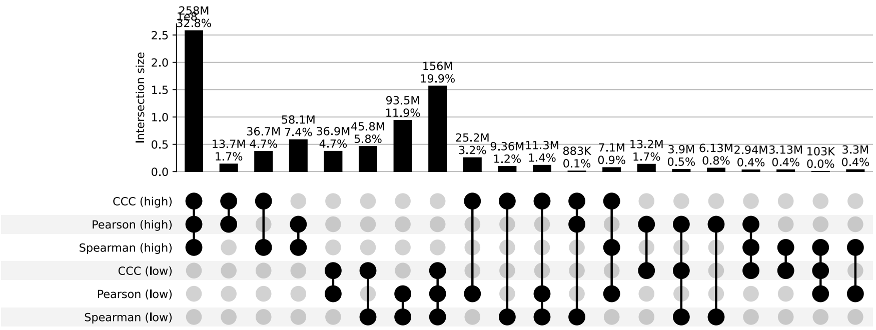


**d)** UpSet plot using permutation-based statistical thresholds

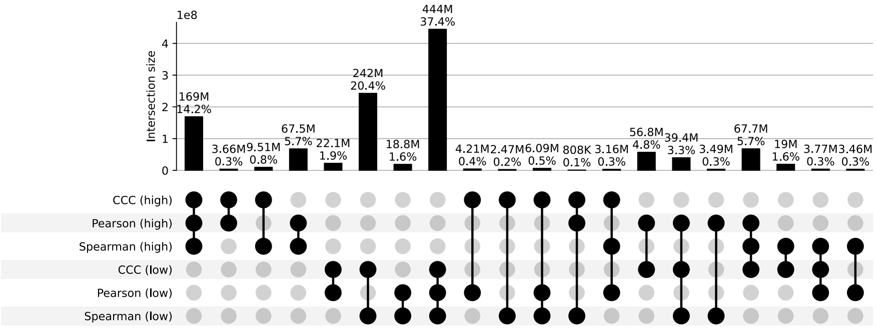


**Figure S54: Distribution and UpSet plots for GTEx v8 vagina.**

#### Whole Blood

**a)** Correlation coefficient distributions between gene pairs within GTEx v8 Whole Blood

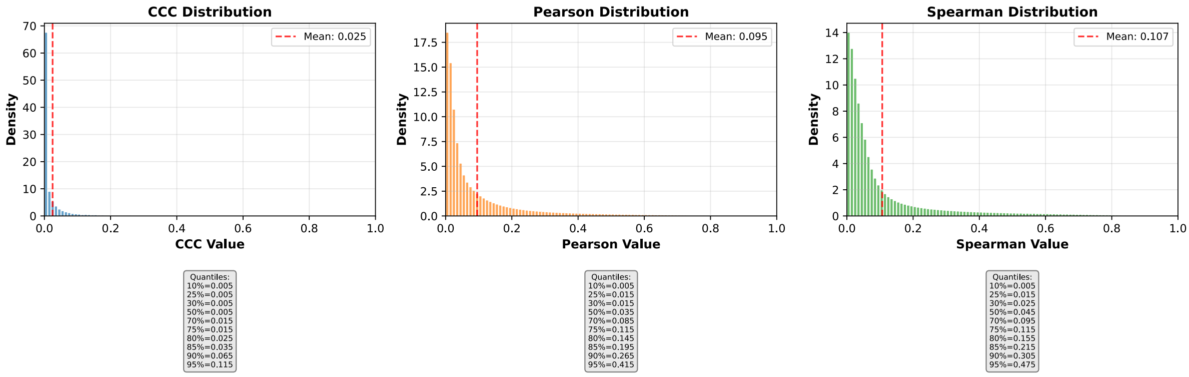


**b)** Corresponding cumulative histogram

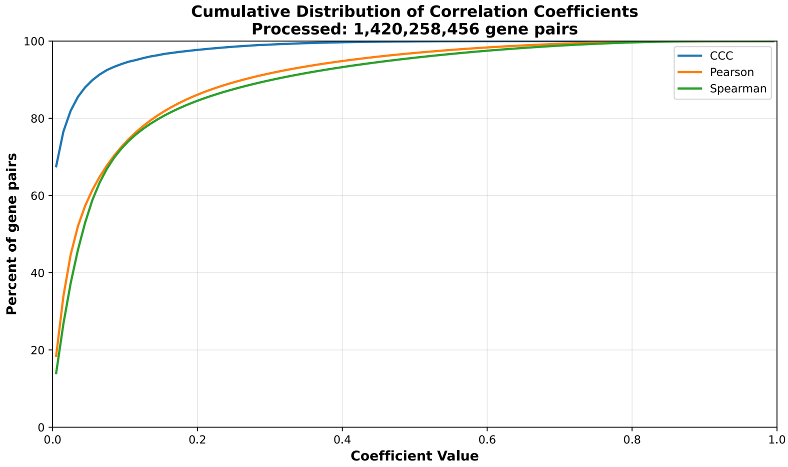


**c)** UpSet plot using top and bottom 30% correlations

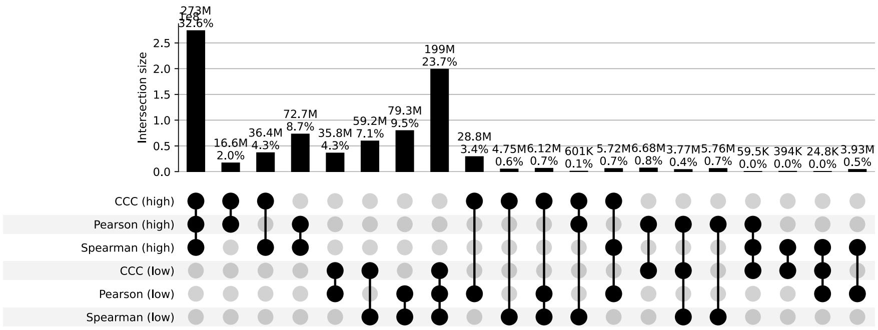


**d)** UpSet plot using permutation-based statistical thresholds

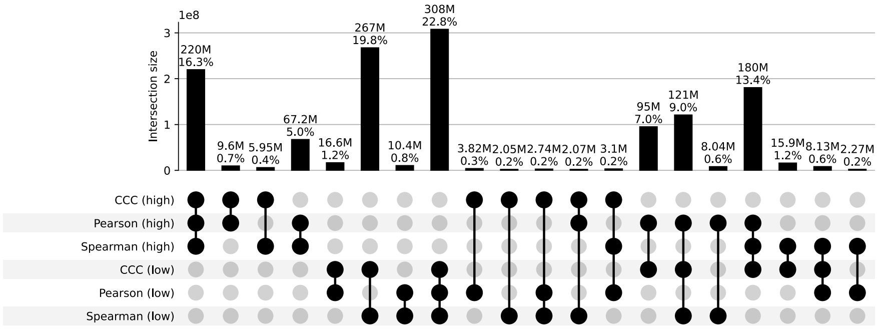


**Figure S55: Distribution and UpSet plots for GTEx v8 whole blood.**
